# Structural basis of half-site reactivity in the catalytic α-subunit of Class Ib ribonucleotide reductases

**DOI:** 10.64898/2025.12.21.695763

**Authors:** Lumbini R. Yadav, Shruti B. Chauhan, Manali Joshi, Shekhar C. Mande

**Affiliations:** National Centre for Cell Science, SPPU Campus, Ganeshkhind, Pune-411007, India; Bioinformatics Centre, Savitribai Phule Pune University, Ganeshkhind, Pune-411007

**Keywords:** RNRs -Ribonucleotide reductase, Mth-*Mycobacterium thermoresistibile*, α_2_ subunit, asymmetry, half site reactivity

## Abstract

Ribonucleotide reductases (RNRs) employ radical chemistry to generate deoxyribonucleotides required for DNA synthesis and repair. A notable feature of RNRs is half-site reactivity, where, despite the enzyme being a symmetric α₂ dimer, only one active site is catalytically active at a time while the other remains in a “poised” state for substrate binding. This phenomenon is tightly linked to the asymmetric α₂β₂ interaction required for radical transfer. Here, we determined cryo-EM structures of the α-subunit in the apo and holo states, i.e., the complex bound to TTP (effector) and GDP (substrate). The structures reveal asymmetric binding of the effector TTP and the substrate GDP across the dimer, with concomitant stabilization of loops surrounding the ligand-binding site. Interestingly, this asymmetry leads to well-resolved N-terminal density for ∼150 residues in the substrate-bound subunit, but weak density for this region in the effector-bound monomer. N-terminal domains are unresolved in both monomers of the apo structure. Isothermal titration calorimetry supports asymmetric binding of pyrimidine effectors with micromolar affinities. Molecular dynamics simulations and three-dimensional variability analysis reveal synchronous motions of loop 2, which together with the N-terminal domain drive alternate opening and closing of the active sites in the two monomers. These conformational dynamics provide key insights into the mechanistic basis of half-site reactivity. Together, these findings provide new insights into the structural dynamics and thermodynamic principles governing regulation and half-site activity in Class Ib RNRs.

**Significance statement:** Ribonucleotide reductases (RNRs) are essential enzymes that supply the building blocks required for DNA synthesis and repair, yet the structural basis of their half-site reactivity has remained unclear. Using cryo-electron microscopy, calorimetry, molecular dynamics simulations, and conformational variability analysis, we show that the catalytic α-subunit of a Class Ib RNR exhibits asymmetric nucleotide binding and coordinated conformational dynamics between the two monomers. These motions drive alternating opening and closing of the active sites and are linked to differential stabilization of the N-terminal region. Our findings suggest that asymmetric conformational gating and N-terminal sampling regulate productive interaction with the radical-generating β-subunit, providing a mechanistic framework for understanding half-site reactivity and allosteric regulation in RNRs.

## 1. Introduction

Ribonucleotide reductases (RNR) catalyse the de novo synthesis of deoxy-ribonucleoside diphosphates from the corresponding ribonucleoside diphosphates. Since deoxy-ribonucleotides are the building blocks of DNA, this enzyme is considered to be essential for DNA synthesis, repair and the existence of any DNA based life form ^1,2^. The reduction of ribonucleotides to deoxy-ribonucleotides involves free radical chemistry, where the radical is maintained within the enzyme and is used when required ^3,4^ . Based on the radical type, co-factor and their oxygen dependency, RNRs have been classified into three major classes I, II and III. Class I RNRs have been further sub-grouped into different subclasses, namely Ia-Ie, based on their subunit architecture and the metal cofactor utilised ^5,6,7,1^

Class I RNRs are composed of two subunits; a large catalytic α-subunit that contains substrate-binding site and a binding site for the effector nucleotide, and a small β subunit harbouring a stable tyrosyl radical. The subunit organization in this class of enzymes is of α_2_β_2_ type, forming an asymmetric assembly during radical transfer from the β to the α-subunit ^8,9,10,11^. The subunits in Class I RNRs are structurally identical, but differ in subtle domain architecture and dependence on ATP for regulation of their activity. Structurally, the catalytic subunit of all RNRs share a highly conserved α/β barrel, a catalytic site and two potential allosteric sites ^11^. With the exception of Class Ib and some Class II enzymes, the allosteric centres are highly conserved among the different RNR classes ^12,5,1^.

Three-dimensional structures of several α and β proteins have been determined from both Classes Ia and Ib ^13, 11, 8, 14,15^. The α-subunit is composed of a N-terminal helical domain consisting of ∼220 residues in Class Ia and ∼150 residues in Class Ib, followed by a α/β barrel domain of ∼480 residues, and a 70-residue αβααβ C-terminal domain. The active site is located in the center of a deep cleft between the N-terminal and the barrel domains. The catalytic reaction is initiated by a free radical transfer from a stable tyrosyl radical in the β subunit to an active site cysteine that gets oxidized to form a transient cysteinyl radical. This radical is involved in the abstraction of 3’-hydrogen from the ribose ring of the ribonucleoside diphosphate substrate ^16^. Substantial conformational rearrangements take place during the catalytic cycle within the α-subunit and concomitantly during complex formation of the α and β subunits.

Many elegant studies have yielded insights into the mechanism of radical formation in the β subunit, mode of association between the two subunits, mechanism of activity, specificity regulation and the structural basis of radical transfer between the two subunits ^9,17,18,19,20,21, 22,18^. One of the most intriguing observations, however, is the formation of an asymmetric complex between the two subunits despite both existing as dimers. This asymmetry has led to the proposal that the enzyme operates through half-site reactivity, wherein only one active site is catalytically engaged at a time. Such behaviour may arise either from asymmetric assembly between the α and β subunits or from intrinsic asymmetry within the α-subunit itself, such that one active site binds substrate while the other concurrently releases product. The conformational principles underlying this half-site reactivity within α-subunit have remained unknown, but are now being addressed at the structural level ^23^. Consequently, several important questions remain unresolved, including the structural basis of half-site reactivity ^24^, the mechanism of allosteric regulation in Class Ib RNRs ^11^, and how the β dimer switches between alternative modes of association with α-subunit ^9^.

In this study, we employed cryo-electron microscopy (cryo-EM) to determine the structures of the catalytic α-subunit of Class Ib ribonucleotide reductase from *M. thermoresistibile* (Mth) in both apo (ligand-free) and holo (TTP/GDP-bound) states. Structural comparisons reveal pronounced conformational heterogeneity, particularly within the N-terminal domain and regulatory loops surrounding the catalytic pocket and specificity sites. Ligand binding induces stabilization of these loops and reshapes the catalytic pocket. Notably, the holo structure exhibits asymmetric binding of the effector and substrate nucleotides across the α-dimer. This asymmetry is reflected in differential N-terminal domain stability, with the substrate-bound monomer or the monomer opposite the effector-bound chain displaying greater structural stability. Isothermal titration calorimetry (ITC) further supports this asymmetry, showing distinct effector binding stoichiometries: purine nucleotides bind at one molecule per monomer, whereas pyrimidine nucleotides exhibit approximately half-site occupancy (∼0.5 per α-dimer), despite both classes binding with micromolar affinities. Three-dimensional variability analysis and atomistic molecular dynamics simulations reveal extensive conformational heterogeneity within the α-dimer, especially in the N-terminal domain, suggesting that conformational sampling may precede productive engagement with the β-subunit. These analyses also provide a structural framework for half-site reactivity, wherein alternating opening and closing of active sites appear to be coordinated through concerted motions of loop 2 across the dimer interface.

Together, these findings offer mechanistic insights into the structural dynamics and regulation of the α-subunit during deoxyribonucleoside synthesis. Given the evolutionary relationship between Mth and *Mycobacterium tuberculosis*, these structures further provide a foundation for future structure-based drug discovery targeting mycobacterial diseases ^25^

## 2. Results

The Class Ib RNRs comprise a dimer of the catalytic α-subunit (NrdE), a dimer of the radical generating β-subunit (NrdF), and the accessory subunit, NrdI. The α-subunit is known to exhibit half-site reactivity, and interestingly, its interaction with the NrdF subunit is also asymmetric ^26, 23^. To investigate the allosteric regulation and functional dynamics of the Mth α-subunit, we employed an integrated structural, computational, and biophysical approach. This enabled determination of effector nucleotide-binding affinities, determination of cryo-EM structures in apo and holo forms, and characterization of inter-domain dynamics using molecular dynamics simulations. SEC-MALS analysis of the purified Mth α-subunit revealed a predominant species of ∼120 kDa indicating a concentration-dependent monomer–dimer equilibrium that shifts toward the dimer, consistent with its functional role **(Supplementary Figure 1).**

### 2.1. Thermodynamics of ligand (effector nucleotide/dNTPs) binding to α-subunit

Isothermal titration calorimetry (ITC) analysis of deoxyribonucleotide binding revealed distinct stoichiometries for purine and pyrimidine effectors. The purine nucleotides, dATP and dGTP, exhibited stoichiometry consistent with one binding site per α-subunit. Interestingly, pyrimidine nucleotides displayed a stoichiometry of 0.5 nucleotide per α-subunit, (i.e.one nucleotide binding per dimer of the α-subunit) (**Figure 1A and 1B)**. The binding affinities for dGTP and dATP were determined to be 4.6 and 1.1 µM, respectively, whereas dCTP, dTTP, and TTP bound with affinities 6.6, 3.3, and 0.64 µM, respectively. Binding of all nucleotides was enthalpically driven, as evidenced by the negative binding enthalpies observed for each interaction. The entropy of deoxynucleotide binding ranges from −10.5 to −73.6 cal/mol/deg suggesting a favourable entropy contribution. Interestingly, TTP shows a very small entropy change. The favorable entropy observed in deoxynucleotides, despite ordering of α-subunit loop upon ligand association, may result from the displacement of highly structured solvent molecules, thereby compensating for this ordering (**Figure 1C).**

**Figure 1:**
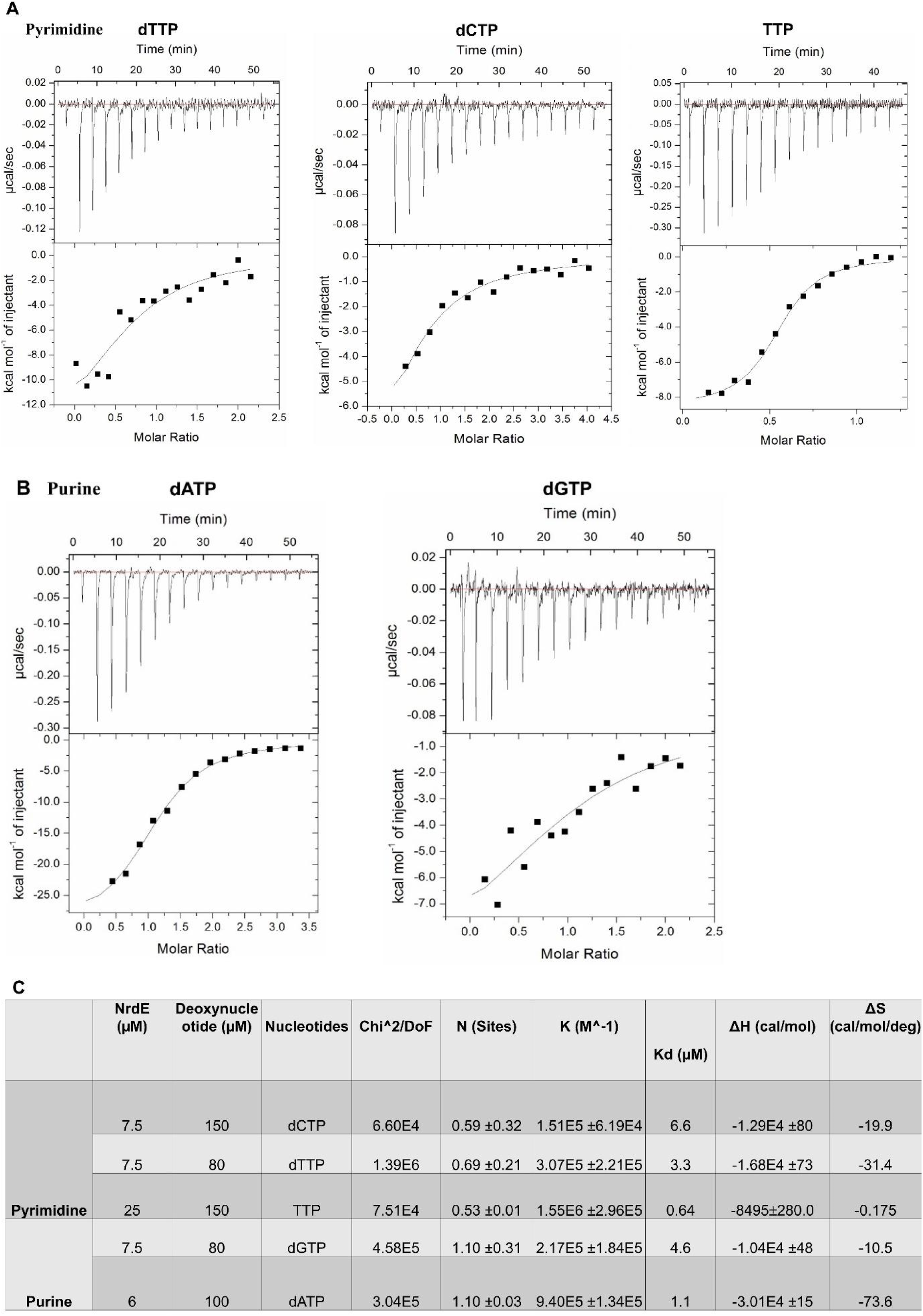
Interaction of α-subunit with effector nucleotides: ITC binding isotherm of deoxyribonucleotides/TTP with α-subunit with top panel showing heat differences obtained after injections of deoxyribonucleotide into α subunit. The lower panel shows the enthalpy changes, corrected for heats of dilution. Data were fitted using a one-site binding model. **A.** Isotherm of Pyrimidine dTTP, TTP and dCTP and **B.** Isotherm of Purine dATP and dGTP. **C.** Thermodynamics parameters derived from heat change is summarized.

### 2.2. Cryo-EM structures of apo and holo α-subunits

Structures of the α-subunit in the nucleotide-bound (holo) and ligand-free (apo) states were determined by single particle cryo-electron microscopy using ∼583,000 and ∼126,000 particles respectively **(Table 1**, **Supplementary Figures 2A and 3A).** The apo α-subunit structure was obtained at a nominal resolution of 3 Å **(Supplementary Figures 2B, 2C).** In he apo structure, the density corresponding to the N-terminal region was weak and fragmented, preventing reliable model building in this region. Consequently, ∼35 residues from the N-terminal region were not modelled **(Figure 2B and Supplementary 2B)**. To provide structural overview of this region and highlight the organization of different loops, the AlphaFold3-predicted structure is shown ^27^ **(Figure 2A).**

**Figure 2:**
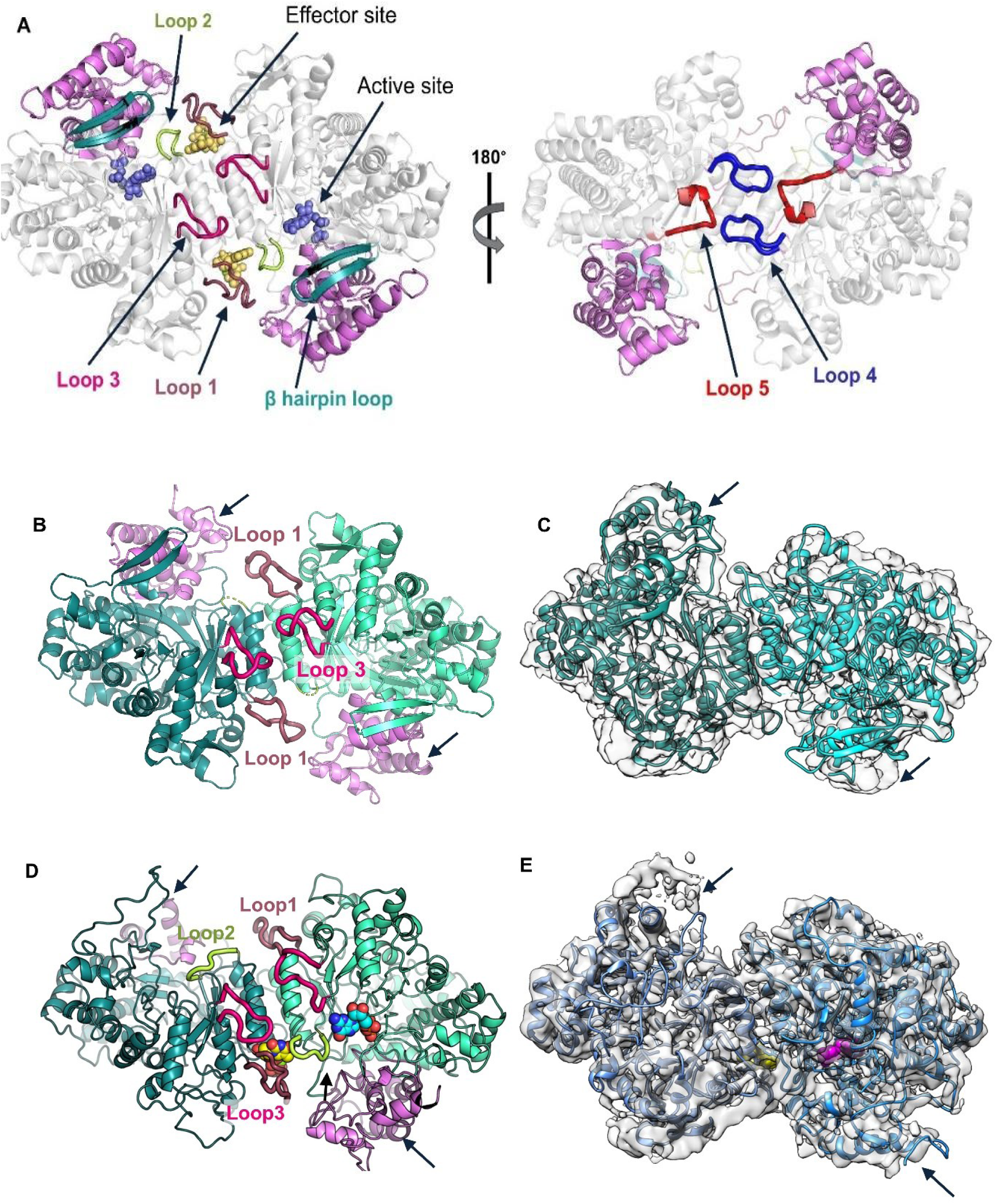
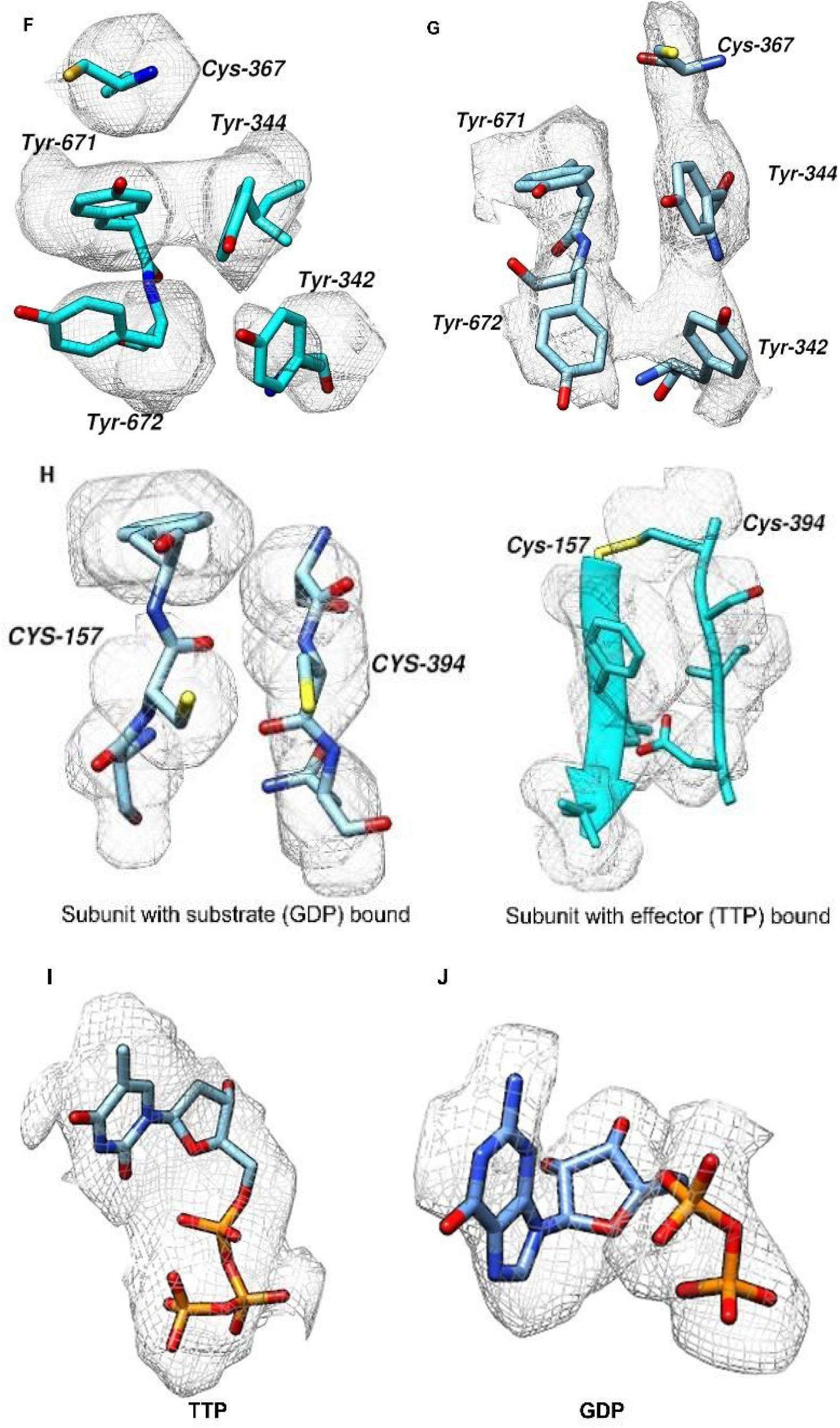
Overview of structures of Mth α-subunit in apo and holo form: **A.** AlphaFold3 model of Mth α-subunit showing different loops involved in effector and substrate binding (loop 1-loop 5) as indicated by arrow. The potential effector and substrate binding site are shown in yellow and blue spheres respectively. The N-terminal region is shown in violet colour. **B.** Cryo-EM structure of apo α-subunit with short N-terminal region indicated in violet. C. Map model overlay of Apo α-subunit. The map sharpened using half map in DeepEMhancer is represented as a transparent surface and the model is shown in cyan. The map contour is 0.04 as displayed in Chimera. **D.** Cryo-EM structure of holo3 α_-_subunit. Yellow and Cyan color spheres in holo density represent effector (TTP) and substrate (GDP) nucleotide binding regions respectively. **E.** Map-model overlay of holo α-subunit. A map sharpened using CryoSPARC (B-factor of −40 Å²) is shown as a transparent surface with a model fitted colored in blue. The map contour is 0.13 as displayed in Chimera. In effector bound monomer the N-terminal region (∼100 residue missing) was not modelled due to weak and fragmented density and substrate bound monomer show density N-terminal density. N-terminal region is indicated by a black arrow. **F.** The density encompassing side chains involved in the active site region of apo α-subunit. The DeepEMhancer sharpened map contoured at 0.01σ with radius cut-off of 2 Å was used to map the density. **G.** The density map of the active site region of holo α-subunit. The CryoSPARC sharpened holo α-subunit map was contoured at 0. 14σ. Colour zone tool of chimera with radius cut-off of 2Å was used to map the active site density. The cut-off radius was chosen to provide an optimal balance between specificity and completeness in mapping density around the atomic coordinates. **H.** Density for cysteine at the catalytic site involved in disulphide bond formation shown for monomer with substrate (GDP) and monomer with effector (TTP). The cryoSPARC sharpened map contoured at level of 0.13 in Chimera with radius cut-off of 2 Å was used to map the density. Difference densities corresponding to the **I.** TTP nucleotide and **J.** GDP nucleotide in the active site of the holo 2 structure are shown. The cryo-EM sharpened map generated in cryoSPARC was contoured at a level of 0.13 and visualized in UCSF Chimera. A protein-only map was generated using the Color Zone function with a radius cutoff of 2 Å. Difference density map shown were calculated by subtracting the protein-only map from the original sharpened map.

**Figure 3:**
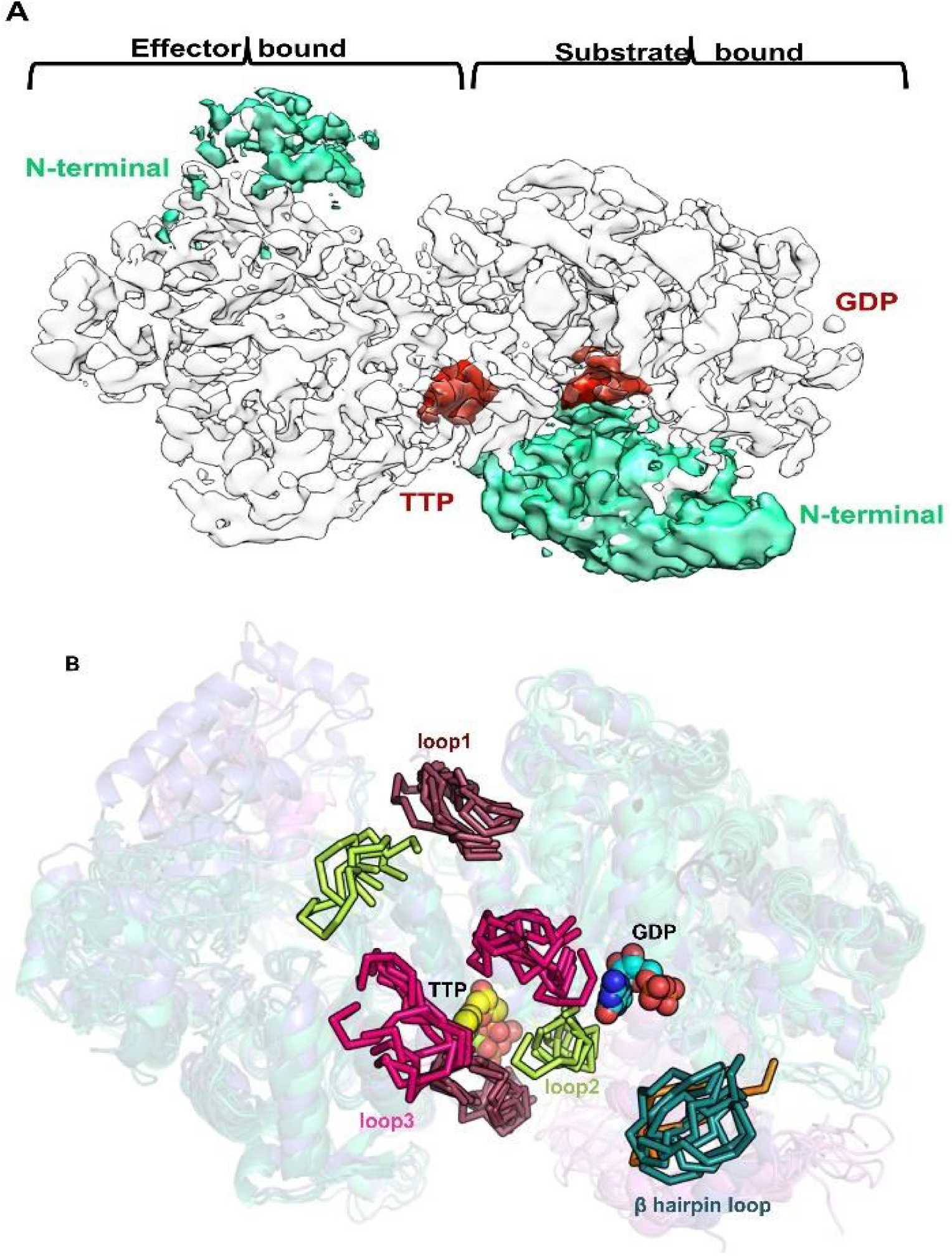
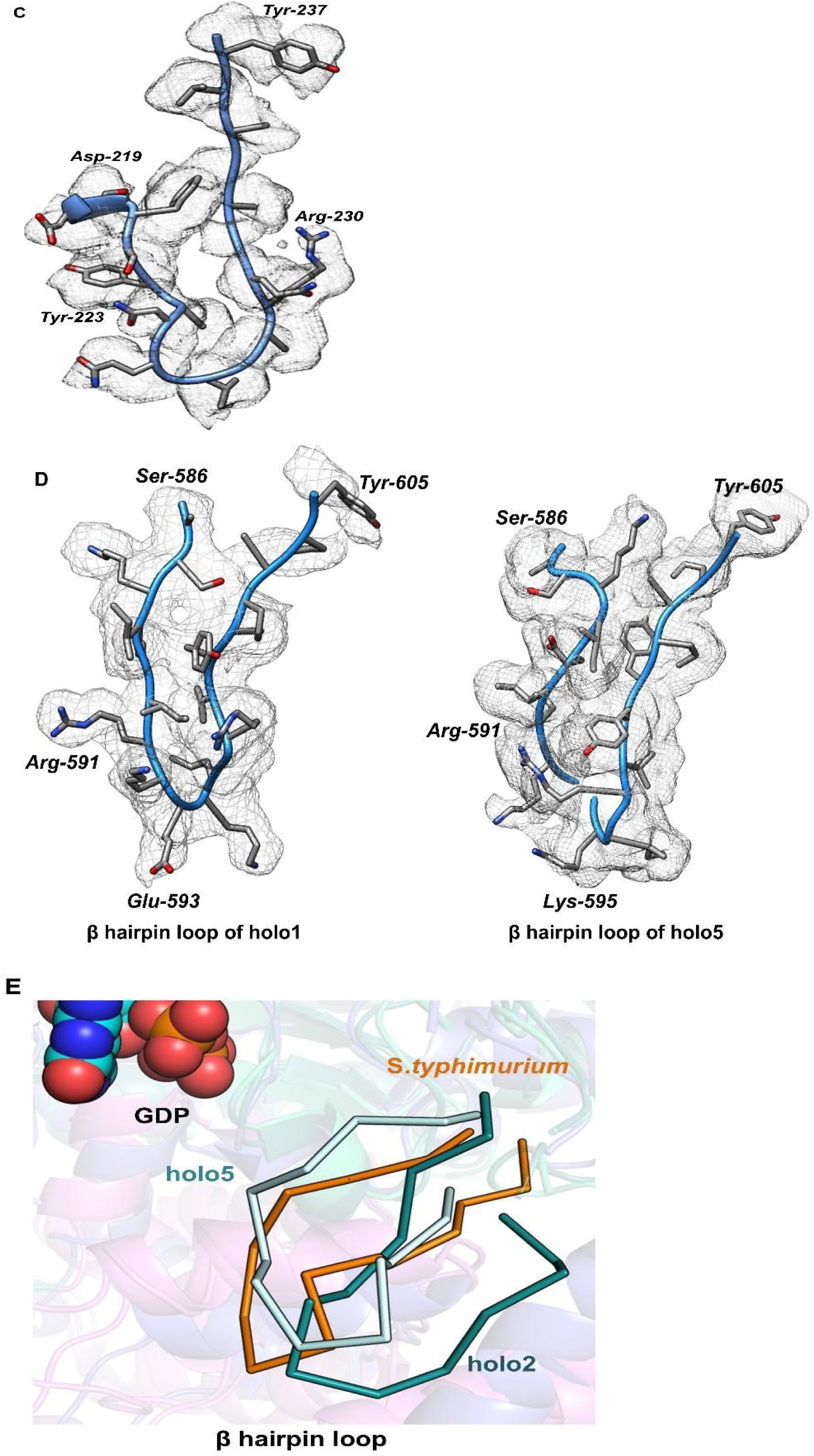
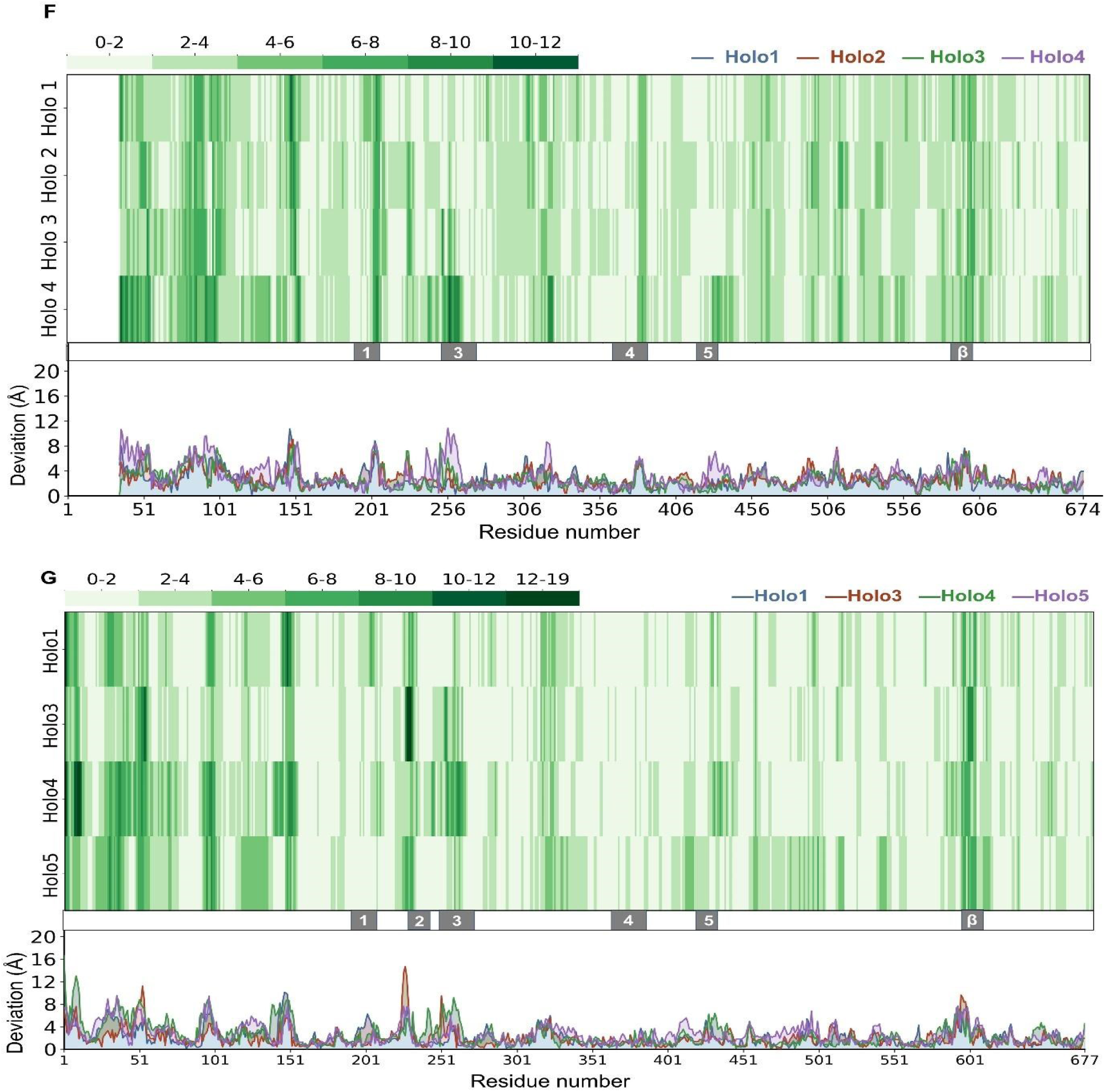
Comparison of apo and holo α-subunit structures: **A.** Representative holo2 map demonstrating asymmetry observed in N-terminal density (cyan), TTP and GDP (red) binding. N-terminal density is poor in all the holo structures in TTP bound chain. **B.** Superposition of holo structures demonstrating variability observed in loop at activity site, specificity site and β-hairpin loop. **C.** Representative cryo-EM density for the loop2 region in the holo3 structure. The density corresponding to loop2 was visualized using the color zone tool in Chimera with a radius cut-off of 2 Å. The cryoSPARC-sharpened map of holo3 was contoured at a level of 0.008. **D.** Representative density of the β-hairpin loop in holo1 (substrate- and effector-bound structure) and holo5 (effector-bound structure). The color zone tool in Chimera was applied with a radius cut-off of 2 Å. Map contour levels were set to 0.09 for holo1 and 0.03 for holo5. **E.** The flexible β-hairpin loop region is highlighted to show the conformational variation among the holo forms with TTP/GDP (cyan; holo2), TTP alone (light cyan; holo5), and the *S. typhimurium* structure. The β-hairpin loop (residues 580–604) in holo2 exhibits a distinct conformation compared to that of *S. typhimurium* (shown in orange). In contrast, the TTP-bound structure (holo5) adopts a conformation similar to that of *S. typhimurium*. The plots show per-residue deviation of substrate-bound holo NrdE chains. Grey shaded regions labelled 1–5 and β denote Loop 1–5 and the β-hairpin loop regions across the residue axis. Coloured lines represent different holo structures, while the green bar indicates the heatmap deviation range in Å. **F.** Deviations were calculated relative to chain A of apo NrdE. Blue, red, green, and purple lines correspond to Holo1, Holo2, Holo3, and Holo4, respectively. Loop 2 (residues 226–230) was excluded due to missing residues in apo NrdE. G. Deviations were calculated relative to the substrate-bound chain of Holo2. Blue, red, green, and purple lines correspond to Holo1, Holo3, Holo4, and Holo5, respectively.

**Table 1:**
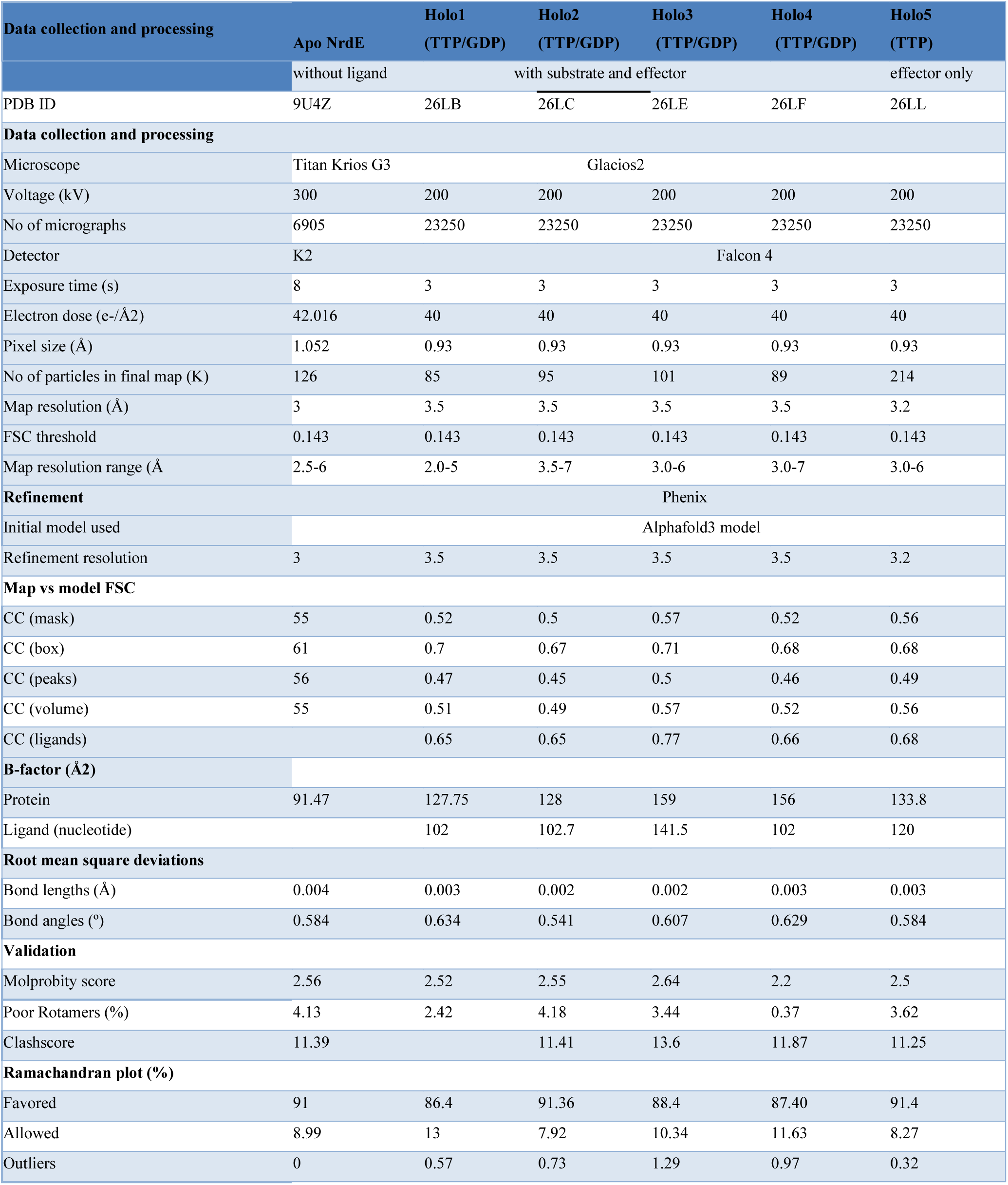
Cryo-EM Data collection and processing details: Cryo-EM data collection and map details of α-subunit in apo and holo form with refinement and validation statistics details.

The holo α-subunit structure was determined in the presence of effector (TTP) and substrate (GDP) nucleotides. The dimer contains binding sites for two TTP and two GDP molecules, however previous studies suggest that two effector (TTP) and only one substrate (GDP) are typically observed per dimer ^27,22,13^ . GDP is expected to bind to only one catalytic site due to half-site reactivity. However, the similarity between the α and α′ subunits and averaging over many particles can hide this difference, making weak GDP density appear on both active sites. We therefore performed focused 3D classification using a mask with two TTP and one GDP. Unexpectedly, the resulting map showed TTP density only in one monomer, and as expected, substrate binding was also restricted to a single monomer within the α-subunit dimer. Therefore, a final focused mask for spatially proximal TTP–GDP pair was generated, followed by 3D classification into six classes. Of the six classes, two classes lacked well-defined density at the substrate-binding site and were therefore merged as an effector-bound class (holo5). The five distinct classes obtained (named as holo1-holo5), each containing between ∼80K–200K particles, were used for structure determination. All classes were independently refined to resolutions of ∼3.2–3.5 Å. **(Table 1; Supplementary Figure. 3E).**

All the distinct holo classes, as well as the apo α-subunit, adopt the canonical α-subunit architecture. Overlay of atomic models with the corresponding cryo-EM maps demonstrates a fit of the models within the reconstructed maps **(Figure 2C and 2E and Supplementary Figure 4)**. Overall, the cryo-EM maps are in good agreement with the refined atomic models, with well-resolved density for both backbone and side-chain atoms. However, the holo maps exhibit anisotropic map features, likely due to preferred particle orientation. **(Supplementary Figures 5A and 5B)**. **Figures 2B and 2D** show the overall cryo-EM structures of the α-subunit in the apo and holo states, respectively. In both structures, portions of the N-terminal region could not be modelled because of insufficient or fragmented density (**Figure 2C and 2E, 3A and Supplementary Figure 4).**

The α-subunit contains a redox-active cysteine pair in the C-terminal region that is required for enzyme reactivation, as well as a catalytic cysteine trio at the active site that is essential for nucleotide reduction. The C-terminal cysteine pair lacks observable electron density in both the apo and holo structures, consistent with previous studies ^13,11, 8^.. At the active site, cysteines are oxidized in the apo form with both monomers displaying disulphide bond formation. In contrast, the holo structure exhibits asymmetric redox states within the dimer, with the substrate-bound monomer showing reduced catalytic cysteines, whereas the effector-bound monomer retaining oxidized disulphide state **(Figure 2H)**. Similar asymmetry in redox states was observed in the holo5 structure. . Clear density is also observed for catalytic residue Cys 367, as well as for clusters of Tyr residues involved in radical transfer and stabilization, including Tyr-671, Tyr-672, together with neighbouring Tyr-342 and Tyr-344 **(Figure 2F and 2G).** Density corresponding to active site substrate GDP is present in all the four 3D classes of holo α-subunit structure (**Supplementary Figures 5C, 5D, 5E and Figure 2J)** whereas density for the specificity site effector TTP is observed in all the 5 holo classes **(Supplementary Figures 5C, 5D, 5E 5F and Figure 2I).** No clear Mg²⁺ density is observed at the effector-binding site, although it is included in the model. Notably, substantial variation in density at both TTP and GDP binding sites is evident across the five holo classes.

### 2.3. Comparative analysis of apo and holo structures of α-subunit

Cross-structure RMSD analysis revealed that all holo structures are relatively similar to one another, with RMSD values ranging from 1.9–2.5 Å. In contrast, comparisons between the apo and holo structures yielded larger RMSD values (2.2–3.0 Å) indicating conformational differences between the ligand-free and ligand-bound states (**Supplementary table 1**). Superposition of the holo and apo structures with an overall RMSD of ∼3 Å suggests that ligand binding does not induce major changes in the global fold, but instead gives rise to localized variability and structural asymmetry **(Figure 3A**). The regions displaying the greatest variability include loops surrounding the active and specificity sites, β hairpin loop (aa ∼580 to 604), loops 4 and 5, and the N-terminal region **(Figure 3B**).

The loops proximal to the specificity and active sites are commonly known as loop 1, loop 2 and loop 3. Electron density corresponding to loops 1 and 3 is observed in both the holo and apo α-subunit structures, whereas loop 2 is resolved only in the holo state **(Figure 3C).** Specifically, loop 2 density is clearly detectable only in holo1–holo4 structures, which contain both effector and substrate nucleotides, while it is poor in holo5, which contains only TTP **(Figure 5)**. In addition, the effector bound chain exhibits comparatively poor/absent loop 2 density in all holo classes. These observations suggest that loop 2 becomes stabilized upon simultaneous binding of both effector and substrate nucleotides and function as a structural bridge linking the specificity and catalytic sites.

Given asymmetric ligand binding in the holo structures, each chain (effector-bound and substrate-bound chains) was individually compared with the apo form. . Comparison of apo structure with the holo substrate-bound chain revealed pronounced conformational variations mainly in loops 1, 3, 4, and 5, whereas comparison with the holo effector-bound chain showed variations mainly in loops 1 and 3 **(Figure 3F and supplementary Figure 6A).** Loop 4 and loop 5 in the effector-bound chain display increased conformational variability relative to the substrate-bound chain, and both loops are linked to the active site cysteines, suggesting a role in allosteric communication.

Other than these loops, weak density is observed for the β-hairpin loop (residues ∼580–604) in the substrate bound chain of the holo structures **(Figure 3D).** This region remains poorly resolved in the effector-bound chain across all five holo classes. In the substrate bound chain of holo1-holo4, the β-hairpin adopts an orientation away from the active site relative to the corresponding *S. typhimurium* structure **(Figure 3E)** ^8^. In contrast, this loop in the holo 5 structure exhibits an orientation more similar to that observed in the *S. typhimurium* structure. However, due to the limited resolution in this region, a detailed structural comparison was not feasible **(Figure 3E)**.

Overall, these structures reveal substantial conformational heterogeneity within the nucleotide-binding pocket and N-terminal region of the RNR α-subunit, highlighting the dynamic nature of the enzyme during ligand binding and catalysis.

### 2.4. Flexibility at the N-terminal domain of Mth α-subunit

The N-terminal region appears highly flexible in both the apo and holo α**-**subunit structures, as indicated by consistently lower local resolution in this region **(Supplementary Figure 2B and 3D**). Superposition of the unsharpened cryo-EM maps of the holo and apo forms revealed a marked absence of density in the N-terminal region of the effector-bound chain in the holo structure. In contrast, this region exhibited only weak and fragmented density in both chains of dimer of the apo structure **(Figure 4A).** Consequently, the N-terminal region exhibits lower average cross-correlation (CC) values for both the chains in apo structure **(Supplementary Figure 7A).** In the holo structures, asymmetric nucleotide binding is observed, with TTP occupying the regulatory site in one monomer and GDP occupying the catalytic site in the opposite monomer (**Figure 4B**). This asymmetry is accompanied by differential stabilization of the N-terminal region, where the substrate bound monomer has density present, however, the effector bound monomer has ∼100 residues missing in N-terminal region **(Figure 3A and Supplementary Figure 7B).** Interestingly, in holo5, which contains only the effector nucleotide, the monomer opposite the effector-bound chain also exhibits intact N-terminal density **(Figure 4C).**

**Figure 4:**
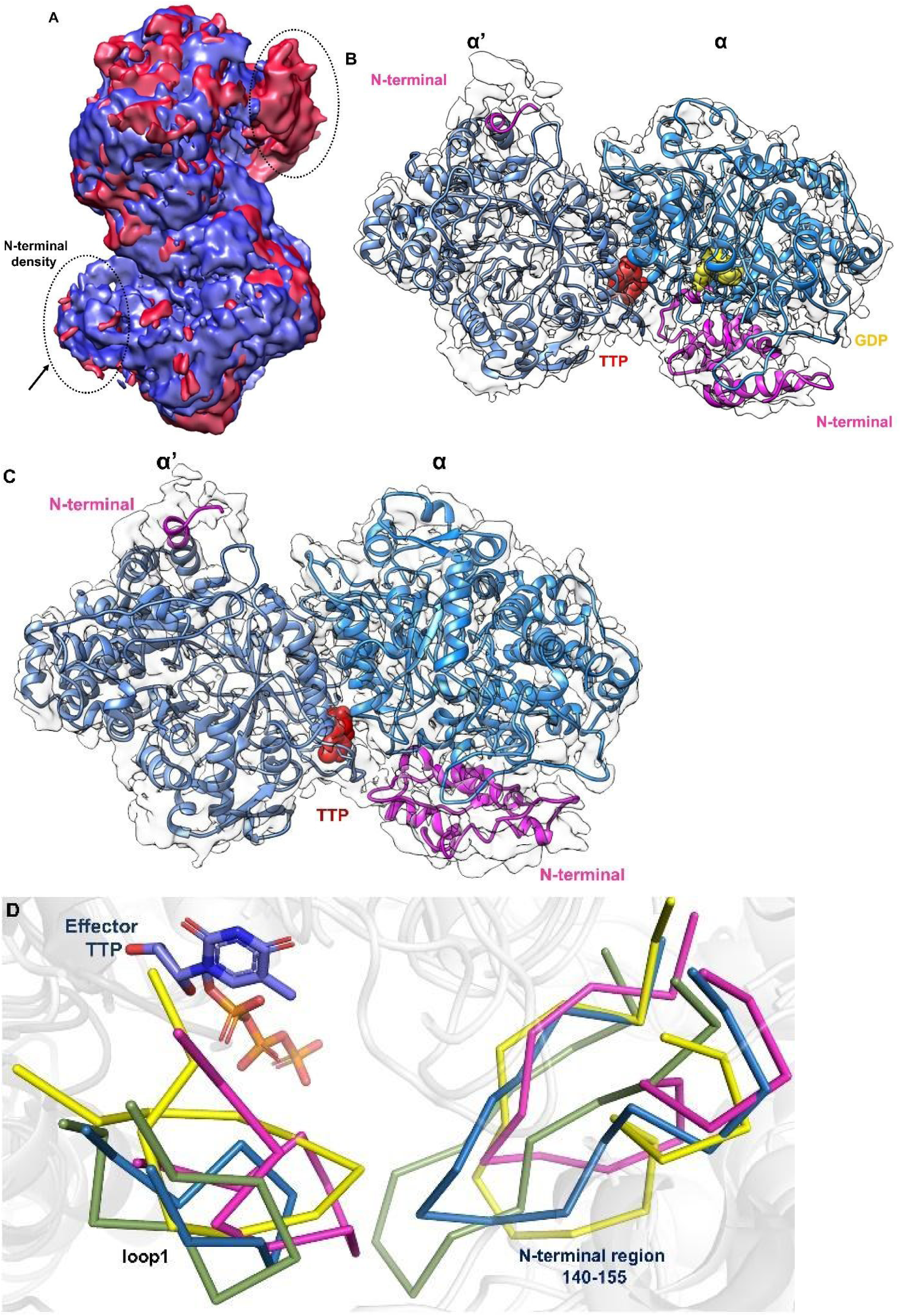
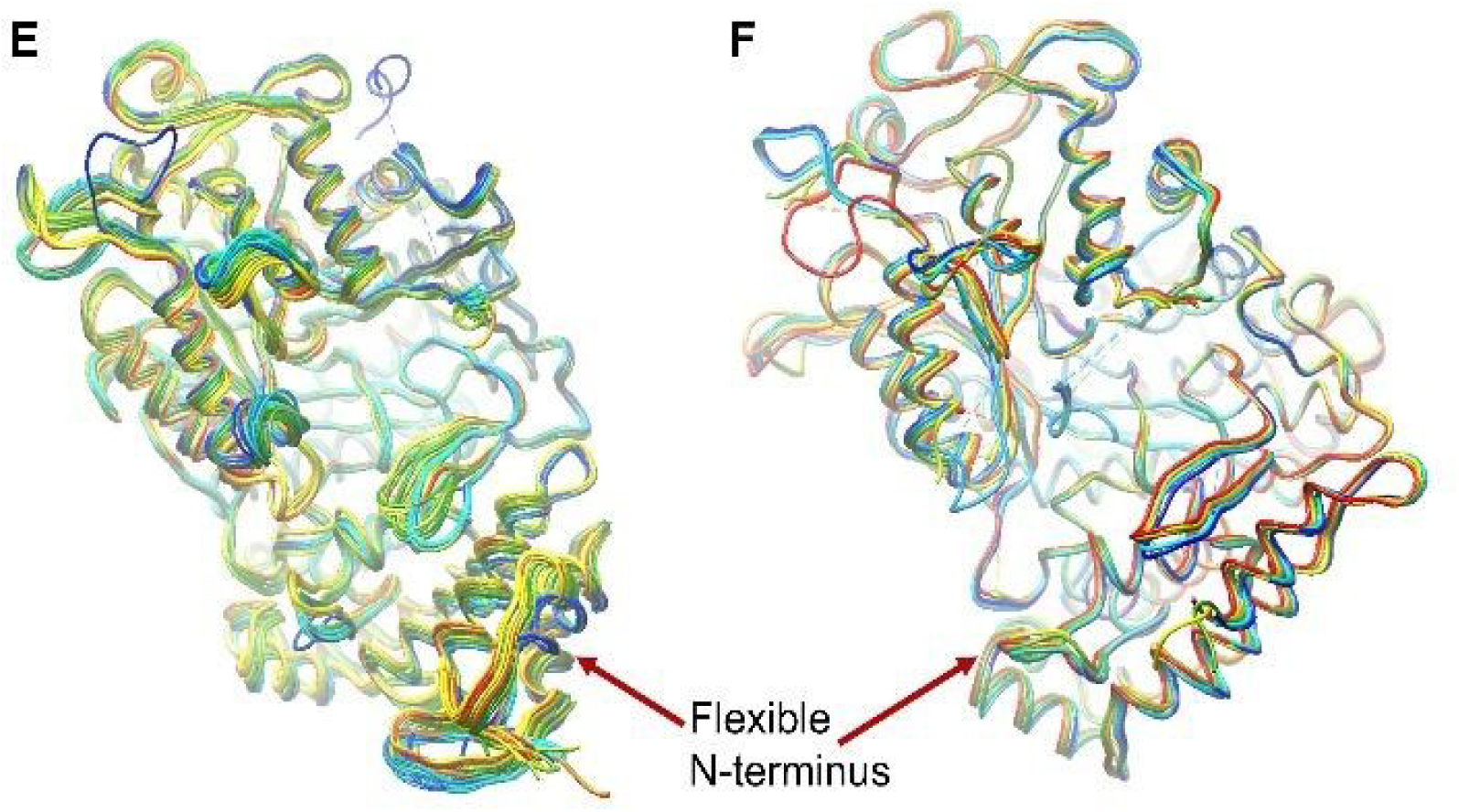
Flexibility in N-terminus of α-subunit: **A.** Superposition of an unsharpened map of apo (blue) and holo (red) α-subunit highlighting conformational variability in the N-terminal region. Dotted circles indicate the flexible N-terminal segment, and the arrow marks the absence of corresponding density in the holo form. B. Cryo-EM structure of the holo2 with TTP (red) and GDP (yellow) showing presence of N-terminal density (magenta) in substrate bound subunit. **C.** Cryo-EM structure of the holo5 class containing only TTP, showing N-terminal density (magenta) in the other monomer**. D.** Structural comparison of Loop1 and the stabilized N-terminal region illustrating conformational variability and increased ordering associated with effector binding. PDBFlex highlighted region of intrinsic flexibility in the different cluster analysed of reported α structures, **B.** in Class Ia, **C.** in Class Ib ribonucleotide reductases.

Comparative analysis of all the holo and apo structures highlights coordinated structural variations in loop1 and N-terminal region. In the apo structure, loop1 region exhibits weak density, whereas in the holo form this loop becomes stabilized upon effector nucleotide binding. Stabilization of loop1 appears to facilitate interactions with the region surrounding Gln 150 in the N-terminal of opposite chain **(Figure 4D)**. In addition, reduction of active site disulphide shifts the region near residue 150 towards the loop1, further stabilising the N-terminal region. Upon substrate binding, the region near residue 140-150 becomes additionally stabilized through interactions with the substrate, which in turn reinforces the substrate binding within the pocket formed by loop 2 and the N-terminal region. Collectively, these observations suggest that effector binding to loop1, together with the reduction of active site cysteines, promotes stabilisation of the N terminal region required for productive substrate binding.

Limited proteolysis also showed rapid degradation of the α-subunit within 15 min, indicating flexible, protease-accessible terminal regions (Supplementary Section 2.3). Proteolytic resistance increased upon β-subunit complex formation or nucleotide binding, suggests stabilization of flexible regions such as loop 1 **(Supplementary Figure 7C).** PDBFlex analyses were performed to analyse the intrinsic flexibility in the different deposited α-subunit structures ^29^. It indicated global flexibility and presence of local flexibility in the N-terminal and different loop regions within clusters of Class Ia **(Figure 4E)** and Class Ib of α-subunit **(Figure 4F).**

### 2.5. Loop2 Dynamics Regulate Substrate Binding and Active-Site Closure

Comparison of structures of different holo 3D classes reveals that the principal conformational variations involve the orientations of the substrate and effector nucleotides, loop 2, and the N-terminal region. Across the four classes (holo1–holo4) containing both substrate and effector, the effector nucleotide TTP occupies a conserved cleft in which the nucleobase is stabilized through π–π stacking interactions with Tyr223, while the phosphate groups are coordinated by Arg194 **(Supplementary Figure 8A)**. TTP binding is associated with increased ordering of loop 2 and promotes a closed conformation of the active site. In holo5, which contains only the effector nucleotide, TTP is stabilized primarily through interactions with Arg194 and Lys201. The catalytic substrate-binding site for GDP is formed by residues Cys157, Cys367, Cys394, Tyr671, and Tyr672, together with residues from the N-terminal region (∼140) and the flexible loop 2 segment, including Arg230 and Gln231 **(Supplementary Figure 8B)**. These residues collectively define the catalytic and substrate-binding pocket. The guanine base of GDP is stabilized through interactions with Arg230, which adopts a conserved inward-facing orientation consistent with its role in substrate coordination within the catalytic pocket **(Figure 5B, 5D and Supplementary Figure 8B)**

Loop 2 exhibits pronounced conformational heterogeneity across holo1–holo4 (**Figure 5A-D)**. The dynamic positioning of Arg230, Gln226, Gln231, and Tyr223, together with differences in GDP and TTP orientation, suggests that the nucleotide-binding pocket samples multiple conformations to facilitate substrate recognition, allosteric regulation, and catalysis. Structural comparison of the four holo states indicates a continuum extending from an open conformation (holo4) to progressively closed conformations in the order holo2, holo1, and holo3. In holo4, GDP is positioned away from the catalytic centre, and Arg230 adopts an outward-facing orientation, leaving the catalytic pocket relatively open (**Figure 5E and 5J)**. In holo1 and holo2, GDP moves closer to the catalytic cysteines, although Arg230 forms direct interactions with GDP only in holo1 (**Figure 5B and 5C)**. Relative to holo2, loop 2 in holo1 is repositioned by approximately 5.9 Å. In the most closed state, holo3, GDP is deeply buried within the catalytic pocket and stabilized by both Arg230 and Gln231 (**Figure 5D and 5I)**. In holo3, loop 2 undergoes a displacement of approximately 13.6 Å relative to holo2 towards active site (**Figure 5F)**. These observations suggest that loop 2 undergoes substantial rearrangement during the transition from open to closed states, thereby guiding the substrate into the catalytic pocket **(Figure 5 and supplementary video 2)**.

**Figure 5:**
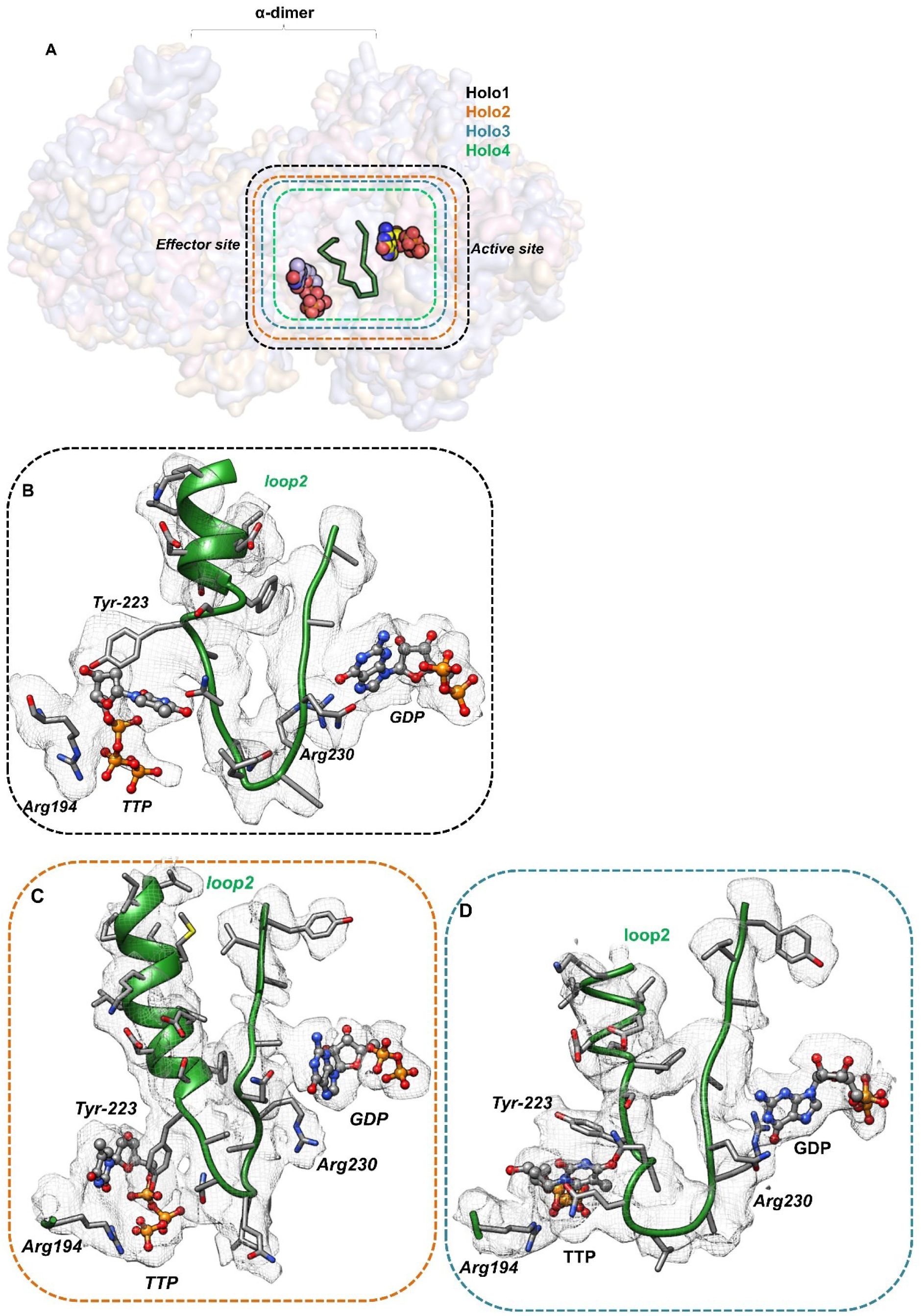
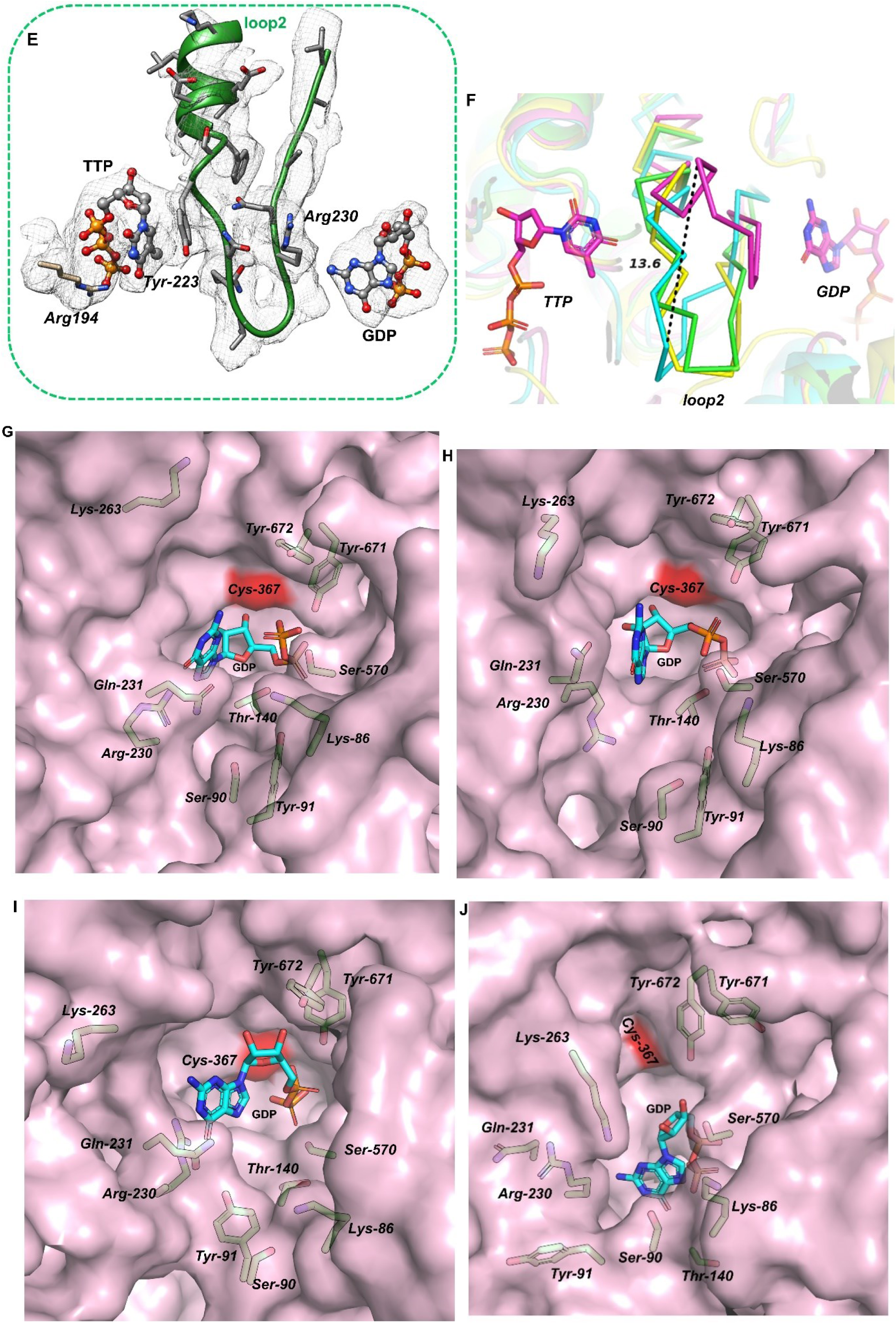
Loop2 Dynamics Regulate Substrate Binding and Active-Site Closure. **A.** Superposed holo α-subunit structures (holo1-holo4) shown in surface representation with effector and active site shown in sphere and stick models with loop2 region highlighted in green. The dotted-line boxes represent the colors assigned to different holo structures, which are described in detail in panel B–E. The cryo-EM sharpened map generated in cryoSPARC was used to visualize structural features of the loop 2 region in UCSF Chimera. A protein-only map was generated using the Color Zone function with a radius cutoff of 2 Å. Difference density maps were then calculated by subtracting the protein-only map from the original sharpened map; **B.** loop2 region of holo1 contoured at a level 0.1 (black box), **C.** loop2 region of holo2 contoured at a level 0.1 (orange box), **D.** loop2 region of holo3 contoured at a level 0.1 (blue box), **E.** loop2 region of holo4 contoured at a level 0.14 (green box). **F.** The loop 2 regions of holo1–holo4 are shown in ribbon representation to highlight structural variations among the holo classes. The distance between residue Gln226 in the two extreme conformational classes, holo3 and holo2, is indicated by a black dotted line and is measured in Å. Surface representation of the active-site pocket with key residues displayed as sticks, highlighting loop 2 and the N-terminal region for **G.** Holo1, **H.** Holo2, **I.** Holo3, and **J.** Holo4.

In addition to loop 2 movements, residues near the active-site pocket and the N-terminal region surrounding residue 140 undergo conformational rearrangements that create space for substrate binding and positioning (**Figure 5G-J)**. In holo4, Lys263 from loop 3 occludes the substrate-binding pocket (**Figure 5J)**, whereas in holo1, holo2, and holo3 it adopts an alternative orientation that avoids steric clashes and permits substrate accommodation (**Figure 5G-I)**. The N-terminal region near residue 140 in holo1–3 shifts toward the catalytic pocket, contributing to substrate positioning and stabilization. Similarly, a slight reorientation of the Ser570-containing loop in holo3 places it closer to the substrate-binding site, potentially enhancing substrate stabilization (**Figure 5G-I)**.

Together, the coordinated motions of loop 2, Lys263, Ser570, and the N-terminal region appear to act as a dynamic conformational gate regulating substrate entry, stabilization, and catalytic progression within the active site.

### 2.6. Molecular Dynamics simulations of the α-subunit

MD simulation for the apo-structure was carried out in seven replicates of 500 ns each, resulting in a total period of 3.5 µs. The simulation trajectory was analysed to characterize residue correlations arising from the dynamic behavior of the α-dimer. Average RMSD upon stabilization for the three replicates showed higher than expected value of ∼6 Å **(Supplementary Figure 9A).** The RMSD for the core and the C-terminal domain lie within a range of 3-6 Å, whereas the RMSD of N-terminal domain varies between 4-11 Å, clearly suggesting that the major contribution to the high deviation arises from the N-terminal domain **(Supplementary Figure 9B).** High root mean square fluctuations (RMSF) are observed in the region of amino acid residues 378-386 (named as loop4) and the β-hairpin loop **(Supplementary Figure 9C).** Similarly, loop 1 (aa 196-206) and loop 2 (aa 225-232), which interact with the effector and the substrate nucleotide respectively, exhibit elevated RMSF highlighting their conformational flexibility. Residue range 254-261 (loop3), 430-438 (named as loop 5) and 500-522 also show significant mobility. These fluctuations are in general concurrence with the variability observed in the Cryo-EM data.

Dynamic cross-correlation matrix (DCCM) analysis revealed regions within each chain that exhibit strong positive correlations among residues belonging to the same structural segment. Based on these correlation patterns, the α-subunit was partitioned into three major domains, namely, the N-terminal domain (residues 1–147), the core domain (residues 148–396), and the C-terminal domain (residues 397–678) (**Figure 6A**). Interestingly, the DCCM analysis further showed that the N-terminal and core domains of one monomer are positively correlated with the corresponding domains of the opposing monomer respectively. In contrast, the core domain displays negative correlations with the N-terminal domains of in each chain (**Figure 6A**).

**Figure 6:**
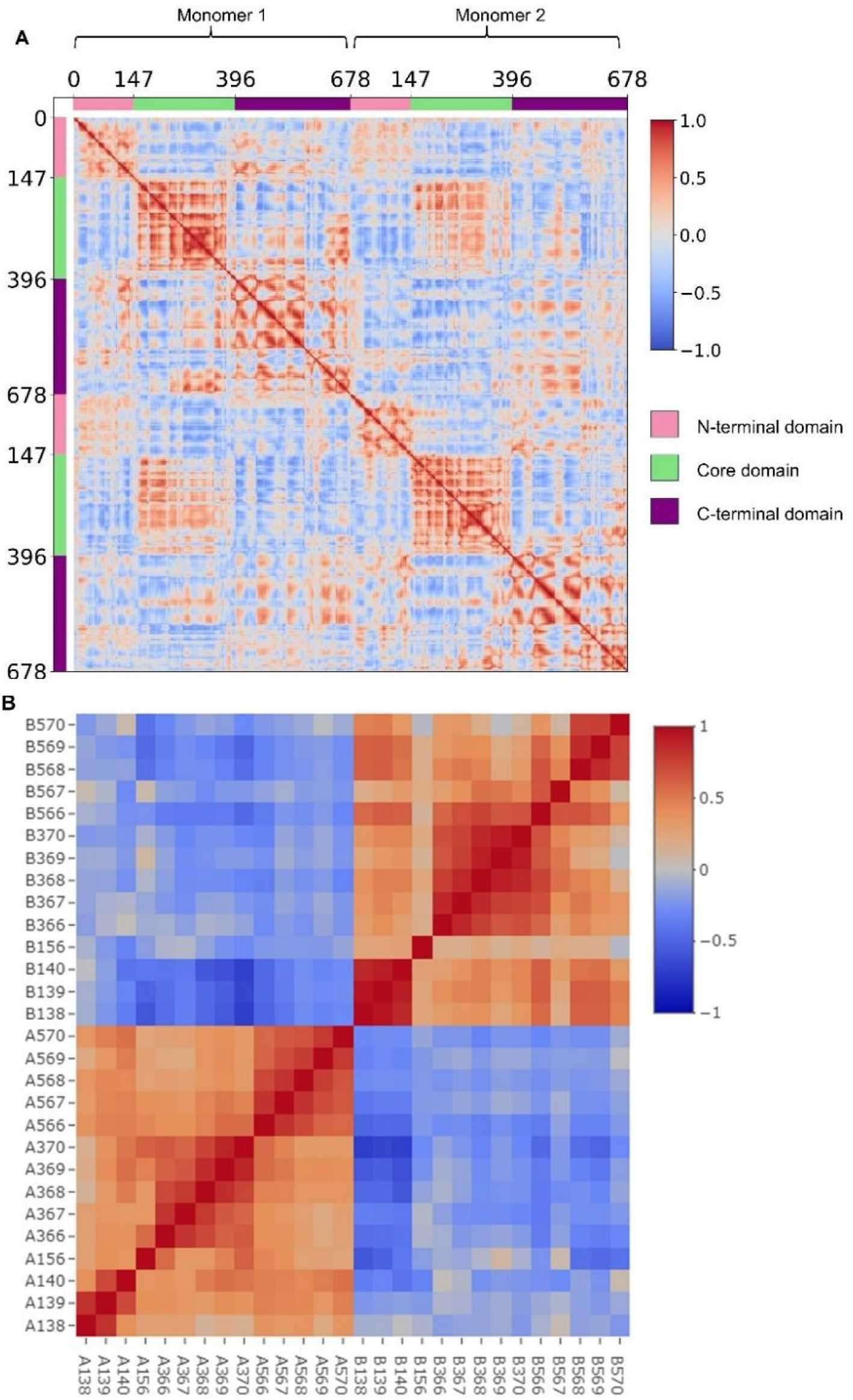
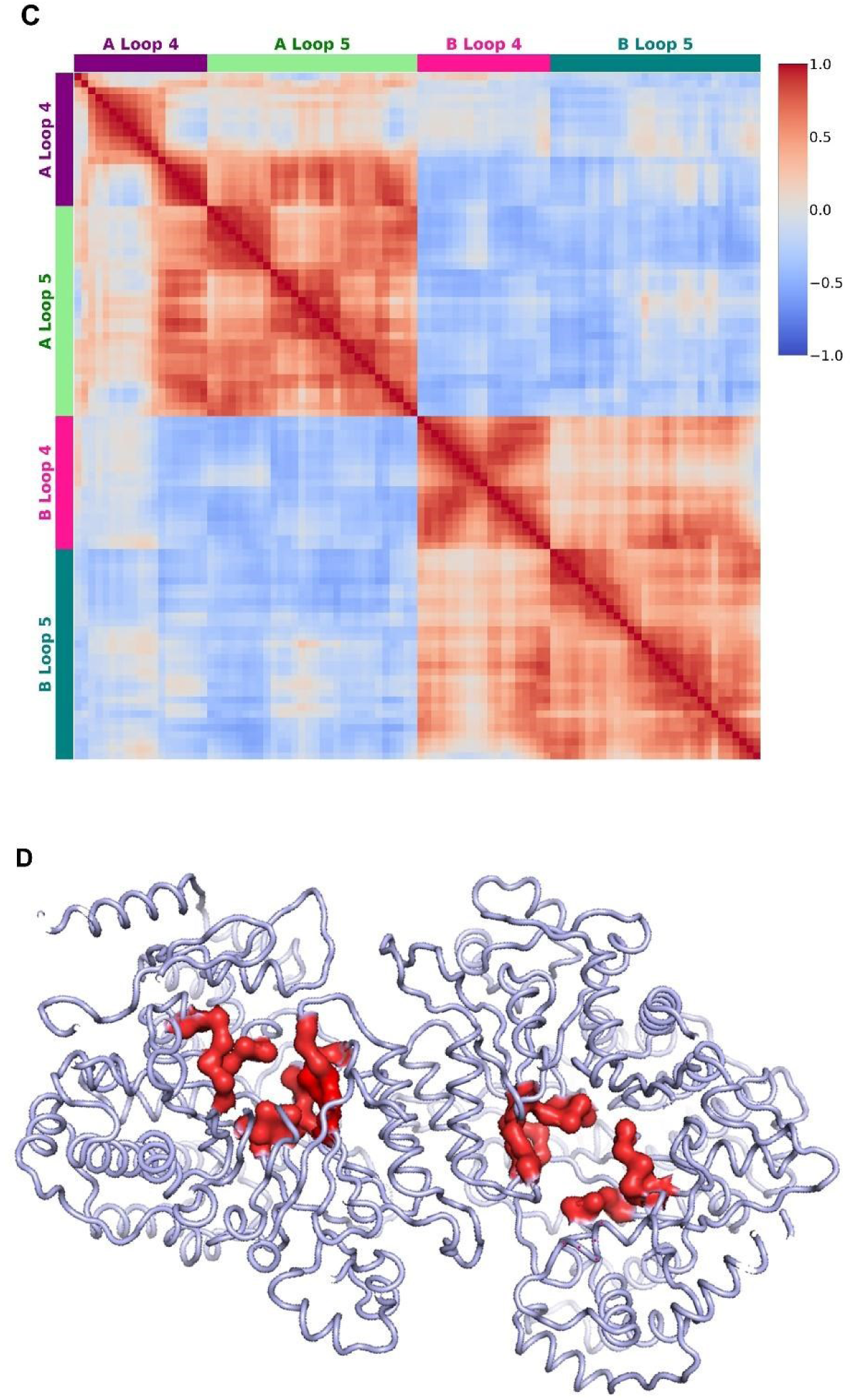
Correlated and anti-correlated motions revealed by DCCM and Principal Component Analysis: **A.** The plot shows intra-chain and cross-chain correlations derived from dynamic cross correlation matrix (DCCM) for the trajectory of a representative simulation replicate. The domain boundaries are marked as N-terminal domain (pink), core domain (green) and C-terminal domain (purple). The x and y axes represent residues of monomer 1 followed by those of monomer 2 of the dimer. The plot is symmetric along the diagonal. The color bar on the right shows correlations ranges from -1 (blue) to 1 (red), **B.** The DCCM for residues present at the active site binding pocket with x and y axes showing residue numbers along with the chain name. The top left and bottom right quarters represent cross-chain correlations whereas the top right and bottom left quarters show intra chain correlations for chain B and chain A respectively, **C.** DCCM specific to Loop 4 and Loop 5 regions. The DCCM highlights Loop 4 (373–391) and Loop 5 (418–447), with purple and pink bars representing Loop 4 of chains A and B, respectively, and green and teal bars representing Loop 5 of chains A and B, respectively. D. Visual analysis of the trajectory along principal component axis 1 shows significant movement of the active site pocket residues highlighted in red (includes the residues shown in (B)). The left monomer tends to have a closed active site while the right monomer has open active site indicating anti-correlation between the two active site pockets of the dimer.

The core domains of the two monomers undergo coordinated motions, whereas residues associated with the substrate-binding site exhibit pronounced cross-chain anti-correlations, as evident from the DCCM analysis (**Figure 6B and 6D**). In particular, loop 4 of one monomer is negatively correlated with loop 5 of the opposing monomer, and vice versa, suggesting that these loops move toward one another in an alternating manner across the dimer interface. By contrast, loop 4 and loop 5 within the same monomer display predominantly positive correlations, indicating coordinated motion in the same direction (**Figure 6C**). These correlated and anti-correlated motions suggest a dynamic inter-subunit communication network that may contribute to coordinated active-site regulation and half-site reactivity in the α-dimer. To further investigate these correlations, the dominant collective motions of the α dimer backbone were identified by principal component analysis of the simulation trajectory. The motions along the first three principal component axes constituted more than 48% of the α dynamics. The projection of the motions along the first principal component axis evidently shows that the opening of the active site in one chain is concomitant with the closing of the active site in the other chain **(Figure 6D) (Supplementary video 1).**

## 3. Discussion

The α-subunit of RNRs serves not only a catalytic function but also plays a central role in allosteric regulation. Here, we present structural and mechanistic insights into the Mth Class Ib RNR α-subunit using cryo-EM, biophysical, and computational approaches. The apo, holo1- 4 (TTP/GDP-bound), and holo5 (TTP-only) structures reveal variability in effector and substrate binding, loop 2 dynamics, and N-terminal organization. Loop 2 emerges as a key regulatory element coordinating substrate accommodation and active-site closure. Despite forming a dimer, the α-subunit exhibits pronounced asymmetry in ligand binding, catalytic-site redox states, and N-terminal density. Dynamic Cross-Correlation Matrix (DCCM) and 3D Variability Analysis (3DVA) analyses further demonstrate anti-correlated motions between substrate-binding regions of opposite monomers, indicating coupling between opening of one active site and closure of the other. Together, these findings highlight intrinsic conformational asymmetry and inter-subunit communication as key determinants of half-site reactivity and catalytic regulation in the Mth α-dimer.

### 3.1. Asymmetric ligand binding and loop dynamics underlie half-site reactivity in α-subunit

The structure of Mth α-subunit reported here exists as a canonical dimer, with the dimer interface consisting of a 4-helix bundle, with 2 α-helices contributed by each monomer. The two monomers in apo bury an area of approximately 1438 Å^2^ upon dimer formation. The apo structure shows a higher interface area, whereas all holo structures exhibit reduced interface areas, indicating ligand-induced conformational changes. Among them, holo2 displays the greatest reduction, suggesting the strongest structural compaction or interface rearrangement. Near this dimer interface several loop regions participate in nucleotide binding and allosteric regulation. Among the three loops involved in nucleotide binding, density for loop 1 and loop 3 is present in both the holo and apo α subunits; whereas density for loop 2 is present only in the holo structures. Loop 1 is known to stabilize the effector, while loop 2 bridges and stabilizes both effector and substrate. Absence of density of loop 2 in the apo structure is corroborated by high mobility as seen in the MD simulations **(Supplementary Figure 9C).** Its presence in the holo structures, with close interactions with the nucleotides, is a strong indication that this loop is involved in conformational selection required for nucleotide binding **(Figure 5).** Low resolution density around the β-hairpin loop (residues 580 to 604) is present in holo α-subunit but is fragmented in the apo form. Interestingly, MD simulations also showed highest fluctuations in this region **(Supplementary Figure 9C).** As the β-hairpin loop region is involved in intimate interactions with the N-terminal domain, this loop might play an important role in initiating conformational sampling by the N-terminal domain upon substrate binding, in order to bind to the β subunit and form the α₂β₂ complex. A major reorientation in this loop was observed in the cryo-EM structure of Mth α_2_β_2_ complex ^23^ in comparison to *S. typhimurium* structure ^8^.

The α-subunit contains three nucleotide-binding sites: the catalytic site (GDP), the activity site (ATP), and the specificity site (TTP). In our cryo-EM holo structure, clear density is observed for GDP and TTP; however, the ATP-binding site could not be confidently modelled, although some diffuse density is present in this region. Interestingly, density for the effector nucleotide (TTP) at specificity site was observed only in single subunit of the dimer unlike previous study. The TTP effector binding site resembles previously reported structures; however, except in holo2, its triphosphate adopts an extended linear density rather than the octahedral coordination seen earlier ^13,11, 27, 39^.. TTP is stabilized by π–π stacking with Tyr223 and coordination by Arg194 (and Lys201 in holo5), promoting loop 2 ordering and active-site closure in substrate-bound states. On the other hand, density for the substrate nucleotide (GDP) is observed at only one active site, consistent with the half-site reactivity reported in previous studies. GDP adopts slightly different conformations across the four holo (holo1–4) states, with the holo2 conformation closely resembling previously reported structure ^13^. The GDP catalytic pocket is formed by Cys157, Cys367, Cys394, Tyr671, Tyr672, and loop 2 residues Arg230 and Gln231, with Arg230 stabilizing the guanine base through an inward-facing conformation **(Figure5)**. Notably, variation in density at the nucleotide-binding site (TTP and GDP) is observed across the five holo classes, indicating heterogeneity in ligand occupancy and/or local conformational differences. Earlier study by Zimanyi et al showed two distinct snapshots of substrate-bound states where a high affinity state is closed and ready for radical transfer whereas in low affinity state is not correctly oriented to undergo catalysis ^28^. We propose that the holo structures represent intermediate states of GDP binding during placement into the active site **(Figure5G-J)**.

The reported α-subunit X-ray structures, indicate a 1:1 stoichiometry at the specificity site ^14,28,33^; however, our ITC analysis reveals one binding site per monomer for purines and half-site occupancy for pyrimidines (one nucleotide per α dimer). This pyrimidine stoichiometry (TTP) at specificity site is consistent with our cryo-EM observations. To the best of our knowledge, this work provides the first ITC-based characterization of all five (TTP/4 dNTPs) effector nucleotide binding to RNRs. Notably, it also reports the first observation of pyrimidine effector binding restricted to a single monomer of the dimer. The reason for this difference in stoichiometry between pyrimidine and purine nucleotides remains unclear. In the GC-rich genome of *M. tuberculosis*, weak binding affinity of dGTP (4.6 µM) and dCTP (6.6 µM), may help maintain elevated intracellular levels of GC and facilitate dynamic responsiveness to the increased GC demand of the genome. TTP (0.64 µM) binds the α-subunit more tightly than dTTP (3.3 µM), with a lower entropic penalty, indicating a more efficient interaction likely due to its 2′-OH group. This suggests TTP is the preferred ligand, stabilizing the regulatory state, whereas dTTP may primarily modulate dNTP pool balance. The affinity values of dATP and dTTP obtained from our ITC experiments **(Figure 1)** are consistent with those reported for *S. typhimurium* ^30^. dATP binds with higher affinity (∼1 µM), supporting its role as a potent feedback inhibitor of RNR activity. Both ATP and dATP bind at the activity site to regulate enzyme function ^34,32^ but the tighter binding of dATP ensures effective control of dNTP production. At elevated concentrations (>1 mM), dATP inhibits Class Ia RNRs but not Class Ib enzymes ^33^. The difference in binding affinities between ATP and dATP is attributed to Tyr244, which sterically restricts nucleotides containing a 2′-OH group ^14,43^.

Other studies have also reported that the affinity value determined for various nucleotides varied in the range 0.1 -80 µM ^35^. SPR studies demonstrated that the effector nucleotides strengthen interaction between α and β subunits, while substrate show no effect in Class Ia RNRs. However, no such affinity enhancement was observed in Class Ib RNRs ^36^.

### 3.2. Conformational sampling of the N-terminal domain

Higher flexibility of the N-terminal domain in both the structures is one of the distinct features of Mth α-subunit. Notably, the N-terminal region is unresolved in effector-bound chain but is clearly visible in the other chain of dimer across all holo (1-5) structures **(Figure 3A).** In contrast, deviation around residue 150 is lower in the effector-bound chain than in substrate-bound chain, suggesting that substrate binding stabilizes the N-terminal region while increasing conformational heterogeneity near residue 150 **(Figure 3 and Supplementary Figure 6).** Such an enhanced flexibility of the N-terminal domain is evident from multiple observations:

a. On an average, the map–model correlation for the N-terminal region is lower in both monomers of the apo α structures. In contrast, in the holo (1-4) structures, the N-terminus is better resolved in the monomer with substrate (GDP) bound at the active site, whereas the effector (TTP) bound subunit shows poor N-terminal density **(Figure 4).** Interestingly, in holo5, where only the effector (TTP) is bound, the N-terminal region of the opposite monomer is also stabilized. Consistent with this, 3DVA of the apo α-subunit reveals an alternating presence and absence of the N-terminal density, indicating significant conformational flexibility **(Supplementary Figure 12C).**
b. The MD simulations show that the N-terminal domain has higher flexibility compared to the rest of the structure **(Supplementary Figure 9B).**
c. Limited proteolysis data revealed faster degradation of α subunit, suggesting a protease-accessible, flexible region near the terminus. Mass spectrometry data confirmed this to be N-terminal region. Other sequence and structure-based analysis predict flexibility in the N-terminal region although the protein is not intrinsically disordered **(Supplementary Figure 7).** Earlier studies also reported weak density for the N-terminal region ^15^.

The destabilized N-terminal region likely plays a critical role in facilitating active-site accessibility and closure, and mediating interactions with the β subunit. The stability of the N-terminal region is determined by the stability of loop 2 and loop 1, which depend on substrate and effector binding, respectively. Effector binding appears to stabilize the N-terminal domain, which in turn helps position the substrate and promotes active-site closure required for nucleoside reduction. Reports indicate movement of β subunit away from α-subunit is essential to release the reduced nucleotide ^13^. It is pertinent to note that α-subunit N-terminal regions in the X-ray structures show higher B-factors and cryo-EM structures indicate poor density and substantial flexibility ^13, 15^. For example, the B-factor plot for the *B. subtilis* α structure indicated a distinctly higher disorder in the N-terminal region ^11^ **(Supplementary Figure 10)**. Thus, collective evidence strongly suggests flexibility in the N-terminal segment of α likely plays a significant role in its function.

The intrinsic flexibility of the N-terminal domain is consistent with conformational sampling, a phenomenon known to play critical role in protein function ^37^. As the principal contact site of the β subunit, this flexibility in α may facilitate productive engagement with β, particularly when the substrate is bound at the active site. Under such conditions, conformational sampling could promote rapid α-β complex formation, positioning key residues from both subunits in close proximity to enable efficient radical transfer ^13^.

### 3.3. Structural insights into loop 2 dynamics coupled to substrate binding

The four holo structures obtained after focused classification showed asymmetric binding for one TTP and one GDP in the dimer. Interestingly, the density for TTP and GDP were present in the vicinity on either side of loop 2. Loop2 is characterised as important loop in specificity regulation ^15, 38^. In the absence of an allosteric effector, loop 2 is disordered and therefore absent in the electron-density maps^14,39,40^. Consistent with this, loop 2 density is absent/poor in our apo structure and effector bound chain of holo structure. However, upon binding of an effector and substrate, the loop becomes stabilized and adopts a well-defined conformation **(Figure 5).** Loop2 exhibits pronounced conformational heterogeneity across the four holo classes (holo1–holo4). Structural comparison reveals a continuum from an open conformation (holo4) to progressively closed states (holo2, holo1, and holo3), suggesting that loop 2 undergoes coordinated rearrangements during substrate entry, binding and catalytic progression. The dynamic positioning of loop2 residues (Arg230, Gln226, Gln231, and Tyr223), together with differences in GDP/TTP binding, suggests that the nucleotide-binding site samples multiple conformations to facilitate effector and substrate recognition, positioning and allosteric regulation. In holo1 and holo3, Arg230 interacts with the GDP nucleobase, whereas this interaction is absent in holo2 and holo4. In the most closed state, holo3, Gln231 additionally contributes to GDP stabilization, suggesting a cooperative role for Arg230 and Gln231 in maintaining the closed catalytic conformation **(Figure 5).**

Arg230, located near the substrate-binding pocket in Mth, appears to stabilize and position GDP during reduction. Similar roles for the equivalent arginine residue have been reported previously. Sequence alignment further shows that Arg230 and Gln226 of Loop 2 are highly conserved across species, supporting their functional importance in substrate recognition and positioning. Previous work by Zimanyi et al reveal how these residues of loop2 regulate substrate preference. The loop 2 residues, Arg293 and Gln294, remodel the active site depending on the identity of the bound allosteric effector ^38^. dATP binding reorients Gln294 toward the active site, favouring pyrimidine reduction (CDP/UDP), while TTP binding shifts Gln294 outward, enabling purine substrate (ADP/GDP) accommodation ^38^. Since our holo structures was determined in presence of effector pyrimidine and substrate purine, Gln 226 is not oriented into the active site pocket. Additional structures with diverse purine and pyrimidine combinations are needed to assess whether the observed regulatory mechanisms and asymmetry are broadly conserved or substrate/effector-dependent. Targeted mutagenesis of residues involved in loop 2 dynamics and the active site will be important to validate their roles in substrate binding, active-site closure, and allosteric communication.

In the *E. coli* class Ia structure, the Arg residue in loop 2 extends into the active-site pocket, where it directly interacts with the substrate phosphate group and stabilize the bound substrate ^38^. Similarly, in the *T. maritima* class II structure complexed with TTP/GDP, the arginine adopts an orientation similar to that observed in the *E. coli* structure ^41^. In contrast, the *S. cerevisiae* structure with dGTP/ADP, although the Arg side chain points toward the substrate phosphate group, it does not make direct contact with it ^39^. In another *E. coli* class Ia structure, the arginine side chain is instead oriented away from the active site ^15^. Notably, the class Ia structures of both *S. cerevisiae* and *E. coli* complexed with TTP/GDP show an arginine orientation resembling the Mth holo2 state, with the side chain positioned away from the active site. Structural information for class Ib ribonucleotide reductases is limited, largely restricted to S. *typhimurium* and *B. subtilis*. In *S. typhimurium*, effector-bound structures (dCTP/dATP) lacking substrate show loop 2 present, but Arg251 is oriented away from the active site, consistent with an open state, whereas in *B. subtilis*, loop 2 lacks observable density even in effector- and substrate-bound structures, indicating pronounced disorder ^11,38^.Together, these observations suggest that loop 2 undergoes substantial conformational rearrangements depending on the bound effector and substrate. Importantly, this study provides the first structural evidence of major loop 2 rearrangements associated with substrate entry into the active site.

### 3.4. Allostery in the α-subunit; Structural Basis of Inter-Monomer Communication

Coordinated motions across distinct domains and loops within the α-subunit play an important role in placement and catalysis of substrate within a single monomer and also mediate inter-monomer allosteric regulation. Within a monomer, binding of the effector nucleotide stabilizes loop 1 and partially loop 2, pre-organizing the enzyme into a catalytically primed state.. Across holo classes 1–4, structural variability in loop 2 is observed, with conformations that guide the substrate into the active-site pocket and maintain it in a reduction-competent orientation **(Figure 5)**. In the substrate-bound chain, loop 2 and the N-terminal region become ordered and move toward the active site, contributing to substrate locking and pocket closure, while loop 4 movement away from the active site helps position the substrate optimally for reduction **(Figure 4B, 4C and Supplementary Figure 13).**

3DVA and MD further reveal coordinated inter-monomer communication that also contribute in regulation of half-site reactivity **(Supplementary Figure 6 and 12).** Analysis of the α₂ model with the 3DVA map and MD simulation shows that the central interface α-helices exhibit coordinated movement that propagates to loop 2, which displays anticorrelated dynamics with loop 4 **(Supplementary video 3).** Anticorrelated movement of loop 4 in one monomer and loop 5 in the opposing monomer contributes to two key events in the α₂ subunit: (i) asymmetric conformational gating ^42^, where the interface α-helix, loop 2, and the N-terminal region move toward the active site in one monomer (closed state) and away in the other (open state); and (ii) a seesaw-like motion in which loop 4 and loop 5 move in opposition, reinforcing dimer asymmetry **(Supplementary Fig. 13 and Supplementary video 4**). This alternating activation may support an asymmetric α₂β₂ model, where one chain is catalytic and the other is poised for substrate binding. We believe all these factors together lead to the allosteric coupling between the two subunits. Together, these dynamics suggest reciprocal opening of one active site and closing of the other. Kang et al. proposed that this coupling is driven by active-site disulfide bond formation ^13^. We propose that these structural and dynamic features collectively underlie allosteric coupling between the two α-subunits, although further biochemical, mutational, and structural studies are needed for validation.

In summary, these findings suggest a dynamic and regulated mechanism in α_2_ subunit involving specific structural movements and interactions, ultimately influencing the allosteric regulation of the protein. The flexibility in the protein at different sites is essential for the fidelity, specificity and efficiency of the enzyme.

## 4. Materials and methods

### 4.1. Purification of Mth α-subunit of RNR

The Mth pGEX-4T1 GST-tagged α construct was expressed in *E. coli* BL21(DE3). Cell pellets from 1.5 L culture were resuspended in lysis buffer (50 mM Tris-HCl pH 7.5, 300 mM NaCl, 0.5 mM EDTA, 3% glycerol, 1 mM PMSF) and lysed by sonication. The lysate was clarified by centrifugation at 15,000 rpm for 35 min at 4 °C. The GST-tagged protein was purified using glutathione affinity chromatography, followed by on-column thrombin cleavage overnight at 4 °C. The cleaved α-subunit was eluted, concentrated using a 10 kDa cutoff filter, and further purified by size-exclusion chromatography on a Superdex 200 (16/600) column equilibrated in 25 mM Tris-HCl pH 8.0 and 150 mM NaCl.

Particles of the α-subunit used for cryo-EM structure determination were derived from cryo-EM grids of the ternary NrdEFI complex. Dissociated NrdE particles from the complex, in both apo and holo states, were selected for reconstruction. Briefly, the NrdEFI complex was co-expressed and co-purified using affinity chromatography followed by size-exclusion chromatography. The purified ternary complex was obtained in a buffer containing 25 mM Tris-HCl (pH 8.0), 150 mM NaCl, and 1 mM MnCl₂.

### 4.2. Grid preparation and data acquisition

Cryo-EM data for the purified RNR ternary complex (NrdEFI) in apo and holo states were collected at the CMO1 beamline and the IISER Pune Bio-cluster Cryo-EM facility, respectively. Apo samples were vitrified on Quantifoil Au grids and imaged using a Titan Krios G3 (300 kV) equipped with a K2 Summit detector at a calibrated pixel size of 1.052 Å.For the holo dataset, the purified ternary complex was reconstituted with MgCl₂, ATP, TTP, DTT, and GDP prior to vitrification on Ultrafoil/Quantifoil Au grids using a Vitrobot Mark IV. Data were collected on a Glacios 2 (200 kV) microscope equipped with a Falcon 4 detector and an energy filter at a pixel size of 0.7 Å.

Movies for both datasets were recorded in counting mode, with 40 frames per exposure under optimized cryogenic conditions. Details of grid preparation are provided in Supplementary Section 1.1.

### 4.3. Cryo-EM image processing and Model Refinement

Cryo-EM image processing for both apo and holo α-subunits was carried out in CryoSPARC following motion correction and CTF estimation. After particle picking, 2D classification, and ab initio reconstruction, particles corresponding to the dissociated α-subunit were selected for further refinement. Final 3D reconstructions were obtained through homogeneous/non-uniform refinement and global/local CTF refinement, yielding a 3.0 Å map for the apo α-subunit **(Supplementary figure 2)** and five holo α 3D classes at resolutions ranging from 3.2–3.5 Å **(Supplementary figure 3)** Structural models were generated by docking the AlphaFold3 model of α-subunit into the cryo-EM density maps, followed by flexible fitting and iterative real-space refinement in PHENIX and manual correction in Coot. The map-model FSC was obtained from PHENIX, which estimated the resolution of 3.9 Å for the apo and 8.16 Å for holo1, 8.34 Å for holo2, 7.16 Å for holo3, 7.8 Å for holo4, 7.06 Å for holo5 respectively at FSC of 0.5. The details of image processing and refinement are described in supplementary section 1.2 and 1.3.

### 4.4. Isothermal Titration calorimetry

Interaction studies of α-subunit with nucleotides (dATP, dGTP, dCTP, dTTP, TTP) were performed using MicroCal iTC200 calorimeter. Protein concentration of 6-25 µM and ligand (nucleotide) concentration of 80-150 µM was used for the titration reaction. Protein concentration was estimated at 280 nm, using Denovix DS 11+ Spectrophotometer. The purified protein and nucleotide were resuspended in same buffer containing Tris-HCl -25 mM pH 8, NaCl-150 mM, Glycerol-2 %, MgCl_2_-5 mM, TCEP-1 mM before the experiment. The protein and nucleotide samples were centrifuged at high speed and degassed at 25°C prior to use. Fifteen injections of 2.5 µl each, and 1st injection of 0.4 µl were injected into the cell allowing an equilibration time of 200 secs between injection. All titrations were measured at 25 °C, with filter period of 8 secs, and stirring speed of 600 rpm. For data analysis each of the reaction were blank corrected. Nonlinear curve fitting was done, using one set of sites in Origin software.

### 4.5. Molecular Dynamics simulation

The cryo-EM α-dimer structure was prepared for MD simulations by modeling the missing loop (residues 225–230) from the holo structure and assigning protonation states at pH 7 using CHARMM-GUI. A disulfide bond (Cys157–Cys394) was included in one monomer. Simulations were run in GROMACS (2021.3) with the CHARMM36 force field ^43^. The system was solvated in TIP3P water, neutralized with Na⁺ ions, and equilibrated under NVT and NPT conditions before 500 ns production runs (7 replicates).

Trajectory analysis, including RMSD, RMSF, DCCM, and PCA, was performed using GROMACS and Bio3D in R ^44^. The details are described in supplementary section 1.8.

### 4.6. PDB submitted details

The atomic coordinates of Mth α-subunit in apo and holo form are deposited with the Protein Data Bank/EMDB. The accession code for α-subunit in apo form is 9U4Z with accession codes EMD-63859. The α-subunit in holo form with GDP and TTP (holo1-holo4) is deposited with PDB ID 26LB, 26LC, 26LE,26LF with EMDB nos EMD-80739, EMD-80740, EMD-80741, and EMD-80742 respectively. The α-subunit in holo form with only TTP is deposited with PDB ID 26LL and EMDB nos EMD-80745 (holo5)

## Supporting information

Supplementary doc

## Acknowledgements

This work was supported by DBT-Centre of Excellence Grant (BT/PR15450/COE/34/46/2016). We thank Dr. Christoph Grunder (University of Washington, USA) for providing *M. thermoresistibile* genomic DNA. Computational resources were provided by PARAM Brahma and PARAM Porul under India’s National Supercomputing Mission. We acknowledge support from the National Cryo-EM Facility funded by the DBT B-Life grant (DBT/PR12422/MED/31/287/2014) and thank Vinoth Kumar Kutti for assistance with sample preparation, cryo-EM data collection, and helpful discussions. Pune Biotech Cluster Cryo-EM Facility at IISER Pune is funded by DBT under the Pune Biotech Cluster project (BT/Pune-Biocluster/01/2015). We acknowledge the European Synchrotron Radiation Facility (CM01), Daouda Traore, and the Cryo-EM facility at IISER Pune (DBT-funded), Shubham Kalyane, for support and assistance in data collection. We also thank the Proteomics Facility at NCCS for mass spectrometry support and analysis. SCM and SBC acknowledge a philanthropic grant from Anand Deshpande to Savitribai Phule Pune University, and SCM and LRY acknowledge DST JC Bose Fellowship support (Government of India).

## Author Contributions

LRY and SCM designed the experimental work and Cryo-EM, and their analyses; SBC, MJ and SCM designed the atomistic molecular simulations protocols and their analyses; all authors contributed to writing the manuscript.

## Competing Interest Statement

Authors declare no competing interests.

## Notes

### Competing Interest Statement

The authors have declared no competing interest.

### Summary of Updates

New cryoEM data was collected with RNR catalytic subunit and nucleotide ligand (GDP, TTP, ATP) and the analysis is updated in the modified manuscript.

