## Supplementary doc for "Structural basis of half-site reactivity in the catalytic α-subunit of Class Ib ribonucleotide reductases"

### 1. Materials and methods:

#### 1.1. Grid preparation and data acquisition:

Data for purified RNR ternary complex (NrdEFI) at 2 mg/ml in apo and holo form were acquired at the CMO1 beamline [1] and the Cryo-EM facility of the Pune Bio-cluster at IISER respectively. For apo NrdE, Quantifoil Au grids (300 mesh, R0.6/1) were glow discharged, and the ternary complex was applied at 18 °C and 100% RH using a Vitrobot Mark IV, followed by plunge-freezing in liquid ethane. Data were acquired in counting mode on a Titan Krios G3 (300 kV) equipped with a K2 Summit direct electron detector at a nominal magnification of 130,000× (pixel size 1.052 Å). Movies were recorded at a dose rate of 9.3 $e^{-}$ $pixel^{-1}$ $s^{-1}$, corresponding to 1.05 $e^{-}$ $Å^{-2}$ per frame, with 40 frames collected over a 5s exposure and a single image per hole.

To acquire data for holo NrdE, FPLC-purified ternary (NrdEFI) complex, in buffer containing 25 mM Tris-HCl (pH 8.0), 150 mM NaCl, and 1 mM $MnCl_2$ at a concentration of 2 mg/mL, was used for grid preparation. Prior to grid preparation, the ternary complex was reconstituted with 5 mM $MgCl_2$, 1 mM ATP, 0.1 mM TTP, and 2 mM DTT, and incubated for 10 minutes. This was followed by the addition of 1 mM GDP and within 35 seconds grid was prepared. Ultrafoil Au (300 mesh, R0.6/1) and Quantifoil Au (300 mesh, R1.2/1.3) grids were glow discharged (90 s, 20 mA), and 3.2 µL of holo ternary complex was applied (wait time 12 s; blot force 2; blot time 3.5 s) at 18 °C and 100% RH using a Vitrobot Mark IV, followed by rapid freezing. Grids were immediately used for data collection. Data were acquired in counting mode on a Glacios 2 (200 kV) equipped with a Falcon 4 detector and a 20 eV energy filter at 165,000× magnification (pixel size 0.7 Å). Movies were recorded at a dose rate of 8.7 $e^{-}$ $pixel^{-1}$ $s^{-1}$ (1.3 $e^{-}$ $Å^{-2}$ per frame), with 40 frames collected over a 3s exposure and multiple images per hole. Data for both holo grids were acquired under similar conditions

#### 1.2. Cryo-EM image processing:

For the apo α-subunit, Beam induced motion was corrected using motionCor2 from RELION [2]. Post motion correction micrographs were imported and further processed and analysed using CryoSPARC version v3.3.2 [3]. Patch CTF estimation was performed to estimate CTF parameters. The particles obtained after multiple rounds of 2D classification were subjected to multiple classes of *ab-initio* reconstruction. The class of ab-initio reconstruction that has particles of α dimer were further assessed and duplicate particles at distance of 20 Å were deleted. The particles of apo (#126K) α-subunit obtained were subjected to ab-initio reconstruction to generate the initial 3D model. Homogenous refinement and masked refinement were performed to obtain 3D maps. To obtain a high-resolution map and improve the map quality, global CTF refinement followed by non-uniform refinement was performed by imposing C2 symmetry [4].

For the holo α-subunit, beam-induced motion was corrected using Patch Motion Correction in cryoSPARC v4.7.1, with one frame from the start and two from the end removed and a Fourier crop factor of 1/2 applied. Patch CTF estimation was performed, and micrographs with CTF fits worse than 10 Å were discarded. Particles were picked using blob and template pickers, cleaned using Inspect Picks and Micrograph Junk Detector, merged, and duplicates removed. Extracted particles (box size 288 pixels, Fourier cropped to 256 pixels) were subjected to multiple rounds of 2D classification. Ab initio reconstruction into six classes followed by heterogeneous refinement was used to isolate dissociated α-subunit particles. A total of 783,875 (Quantifoil) and 178,379 (Ultrafoil) particles were selected for further analysis. Patch motion correction was applied to the micrographs using a Fourier crop factor of 3/4, followed by CTF estimation. The particles of dissociated α-subunit were then re-extracted for final map reconstruction. Combined particles underwent ab initio reconstruction, homogeneous refinement, and global and local CTF refinement followed by non-uniform refinement.

Density corresponding to one substrate and one effector molecule was observed. Therefore, a focused mask was generated around the nucleotide-binding region (dilation radius 8 pixels, soft padding 12 pixels), and 3D classification into 20 classes without alignment was performed. Poor-quality class was discarded, and good classes were merged and subjected to a second round of 3D classification into six classes. Model building and refinement were performed for all six classes. Classes containing only effector density were merged, resulting in five distinct classes. These were further refined using global and local CTF refinement, followed by high-resolution local refinement in cryoSPARC. No symmetry was imposed at any stage for holo classes.

The final classes (holo1–holo5) contained ~85k, 95k, 101k, 89k, and 214k particles, respectively. The resolution at the FSC = 0.143 criterion was 3 Å for the apo α structure while the five holo α 3D classes exhibited resolutions ranging from 3.2 to 3.5 Å. A DeepEMhancer-sharpened map, generated from the half maps, was used for model building of the apo α-subunit [5]. For the holo α-subunit, maps for all classes were sharpened in CryoSPARC using a B-factor of −40 $Å^2$. Unsharpened maps were used to demonstrate the flexible N-terminal region. FSC curves for all the data were calculated with half maps using Validation (FSC) in CryoSPARC. Local resolution of the map was estimated using CryoSPARC local resolution estimation job presented using surface Color tool in UCSF Chimera [6].

#### 1.3. Model Refinement:

Final 126K particles for the apo form and 583K for the holo form were used for structure determination in CryoSPARC [3]. $\alpha_2$ subunit model from AlphaFold3 was docked into cryo-EM maps [7] using local optimization in Chimera using fit in map option [6]. The flexible fitting of the α dimer in the cryo-EM map was done using iMODFIT and Namdinator server [8]. The model thus obtained was further refined in real space refine in PHENIX and manually in Coot [9]. Effector (TTP) and substrate (GDP) were incorporated into the model from the monomer library in coot. The nucleotides were manually positioned into the corresponding density and adjusted to optimize fit and stereochemistry. Refinement included minimization, rigid-body refinement, and local grid search, along with application of stereochemical restraints to maintain proper geometry. Secondary structure, Ramachandran and reference model restraints were applied throughout the refinement process. Multiple rounds of refinement in Phenix were performed to obtain a better model with good stereochemistry and optimal fit to the density. The map-model FSC was obtained from PHENIX, which estimated the resolution of 3.9 Å for the apo and 8.16 Å for holo1, 8.34 Å for holo2, 7.16 Å for holo3, 7.8 Å for holo4, 7.06 Å for holo5 respectively at FSC of 0.5.

#### 1.4. 3D variability analysis:

To explore the heterogeneity and flexibility in the apo and holo datasets, CryoSPARC was used for 3D Variability Analysis (3DVA) [10]. For the apo dataset, particles from the best 2D classes were selected for homogeneous refinement. The particles and mask obtained from this homogeneous refinement were subsequently subjected to 3D variability analysis. A total of three components were solved at a filter resolution of 5 Å. For the holo dataset, all holo particles from the 3D classes containing the effector and substrate nucleotide were merged, and the output from local refinement was subjected to 3D variability analysis. A total of three components were solved at a filter resolution of 6 Å. The output from 3DVA was analyzed using the 3DVA Simple Display with 20 frames for each component [10].

#### 1.5. SEC-multiangle light scattering:

110 µl of SEC-purified protein at a concentration of 9.9 mg/ml of Mth α-subunit was subjected to size exclusion chromatography coupled with multi-angle light scattering (SEC-MALS). Superdex 200 10/300 GL column; (GE Healthcare) with a buffer containing 25 mM Tris-Cl pH 7.8, 150 mM NaCl, 1 mM $MgCl_2$ and 2% glycerol at room temperature. BSA (5 mg/mL) was used as a standard, and data was analyzed with Astra software (Wyatt Technologies).

**1.6. Limited proteolysis:**

The purified proteins (α, α with nucleotide, $\alpha_2\beta_2$ complex and BSA) were subjected to trypsin degradation. The detail of $\alpha_2\beta_2$ complex purification is mentioned in Yadav et al 2025 [11]. For α with nucleotide, α-subunit was reconstituted with nucleotide (TTP and GDP) in the presence of 1 mM $MgCl_2$ and 1 mM of DTT and incubated in ice for 5 minutes and further used for assay. Protein concentration was estimated using nanodrop, and a concentration of ~ 20 µg was used for the reaction. The molar ratio of trypsin to protein was 1:50. Protein was incubated with trypsin at 37°C for periods of 3, 7, 30, 60, and 90 minutes. Reaction was stopped by addition of 1 mM PMSF, and the mixture was heated at 80°C for 3 to 5 minutes in SDS loading dye. Proteolysis digests were analysed on 12% SDS PAGE.

**1.7. In-gel digestion and LC-MS/MS proteomic analysis:**

After destaining in 50% methanol, the SDS gel was washed thoroughly with water, and the band of interest at ~ 18 kDa was excised from the gel for further processing. The gel piece was further excised into smaller pieces. The excised gel pieces were destained, reduced, and alkylated as mentioned in the protocol [12]. Briefly, each proteome sample was reduced with 10 mM DTT, alkylated with 50 mM iodoacetamide, and subjected to digestion using trypsin enzyme (1:50). The peptides were extracted using extraction solution containing a combination of ACN, $H_2O$, and trifluoroacetic acid by vortexing and collecting the supernatant for speed vac. After complete drying, the peptides were resuspended in 50 µL of 0.1% TFA. Furthermore, these digested tryptic peptides were desalted using C18 ZipTip tips. The speed vac-dried samples were further reconstituted in water containing 0.1% v/v formic acid. All reagents of LC-MS grade were used in the process.

MS/MS data was acquired in Orbitrap Fusion™ mass spectrometer coupled to an EASY-nLC™ 1200 nano-flow LC system equipped with EASY-Spray PepMap C18 column. For each MS data acquisition, 1 µg desalted tryptic peptides was injected into the Orbitrap Fusion mass spectrometer. Peptides were separated using a 5%–35% solvent B, containing 0.1% V/V formic acid in 80% V/V acetonitrile at a flow rate of 300 nL/min for a 140 min gradient process. The solvent A consisted of 0.1% V/V formic acid in LC-MS grade water. The mass spectra were acquired in positive ionization mode with a positive ionization spray voltage of 2 KV. The MS scan began with an analysis of the MS1 spectrum from the mass range 375–1500 m/z. Analysis was performed using the Orbitrap analyzer at a resolution of 60,000 with an automatic gain control (AGC) target of $4 \times 10^5$ and maximum injection time of 50 msec. MS2 precursors were fragmented by high-energy collision-induced dissociation (HCD) and analyzed using the Orbitrap analyser of the Thermo Scientific Orbitrap Fusion mass spectrometer (NCE 35; AGC $5\times10^4$; maximum injection time 22 ms, resolution 15,000 at 200 m/z).

The MS data was analysed to identify proteins using the Proteome Discoverer software (version 2.2; Thermo Fisher Scientific, Inc.). This was carried out by employing the MS Amanda 2.0 and Sequest HT search engine with a 1% false discovery rate (FDR) and the cut-off criteria of 2 missed cleavages. Database searching included α-subunit from the *Mycobacterium* UniProt reference number G7CEK2. Total protein level analysis was performed using a 10 parts per million precursor ion tolerance. The product ion tolerance used for the data analysis was 0.05 Da. Oxidation of methionine residues (+15.995 Da) was kept as a variable modification, whereas carbamidomethylation of cysteine residues (+57.021 Da) was kept as a static modification. Peptide-spectra matches (PSMs) were adjusted to an FDR of 0.01. PSMs were identified, quantified (using MS/MS fragment intensity), and narrowed down to a 1% peptide FDR and then further narrowed down to a final FDR protein level of 1% for protein-level comparisons. A minimum peptide length of 4 amino acids and complete cleavage from trypsin were considered during analysis. Target decoy-based peptide-spectrum match validation was used to confirm the protein identification with strict FDR of 0.01 for highly confident peptides and a relaxed FDR of 0.05 for moderately confident peptides.

**1.8. Molecular Dynamics simulation:**

The α dimer structure determined from cryo-EM was prepared for MD simulation as follows: the missing loop 2 (residues 225-230) of the apo form was grafted from its holo form and the protonation state of residues were adjusted for pH 7 using CHARMM-GUI [13,14,15]. In the dimer, only one monomer was modelled with the Cys157-Cys394 disulfide bond. MD Simulations were performed using GROMACS version 2021.3 [16,17,18,19] with the CHARMM36 forcefield for the protein [20]. The α dimer was solvated using TIP3P water model in a cubical box with edges at least 15 Å away from the protein surface. 18 $Na^+$ ions were added to neutralize the system. Periodic boundary conditions were applied and energy minimization was performed using the steepest descent algorithm. The LINCS algorithm was used to constrain the bonds with hydrogen atoms [21]. Short-range electrostatics were treated with a 10 Å cut-off, while long-range electrostatics were calculated using the Smooth Particle Mesh Ewald (SPME) method [22]. Position-restraint equilibration was performed first under the NVT ensemble using a modified Berendsen thermostat with 300K reference temperature for 100 ps. This was followed by pressure coupling in the NPT ensemble controlled by Parrinello-Rahman barostat with 1 bar pressure for 1000 ps [23,24]. The unrestrained production simulation was then performed on the well-equilibrated system for 500 ns in 7 replicates using a timestep of 2fs and the coordinates were saved at every 100 ps.

The average and domain-wise root mean square deviation (RMSD) of the protein and root mean square fluctuation (RMSF) of the residues was calculated using standard tools in the GROMACS package. The dynamic cross correlation matrix for Cα atoms over the trajectory was generated using bio3d package in R [25] . The principal component analysis of the Cα atoms across the trajectory was performed in R and the motions along the first principal component axis were visualized.

#### 1.9. PDBFlex analysis:

The PDBFlex database provides clusters of identical structures and their comparative analysis of intrinsic flexibility. The sequence of Mth α-subunit was used as a query to search the database, and the cluster of hits was investigated. Comparison of the α-subunit structure with reported structure of high sequence similarity (69.9%) (Class Ib 6MVE cluster) and low sequence similarity (49.2%) (Class Ia 5R1R cluster) synergistically indicated flexibility in the loop region and N-terminal region [26].

#### 1.10. Cα difference distance matrix:

The distances between all Cα pairs within a chain and between the two chains of the dimer were calculated using a Python script for both the apo and nucleotide-bound forms of the α-subunit dimer. The difference in the Cα distances between the two forms of protein was obtained in the form of a matrix and plotted using R [27]. All figures and images were made in PyMOL and Chimera [6, 28].

#### 1.11. Normal Mode Analysis:

NMA was performed using the command line version of Webnm@ v3, which is a tool that calculates normal modes based on the ENM (elastic network model) [29]. It generated PDB files for different modes (modes 7-12), a correlation matrix, a correlation plot, and mode visualization animations. The raw protein structures as obtained experimentally were first prepared by adding the missing residues and joining the breaks in the bonds using the software Discovery Studio (version 2021) [Biovia, D.S. (2019) Discovery Studio Visualizer. San Diego.] and Coot [9]. The protein structures were energy minimized and then used for normal mode analysis. The analysis with apo and holo α dimer, and α with nucleotides at effector sites did not show any significant difference among the three.

### 2. Results:

#### 2.1. Oligomeric characterisation of Mth α-subunit:

The Mth α-subunit was purified using GST-affinity chromatography and eluted at 78 ml from 16/600 Superdex 200 size exclusion column **(Supplementary Figure 1A and 1B)**. SEC-MALS analysis revealed a major elution peak corresponding to a molecular mass of ~120 kDa, compared with the theoretical 79 kDa of a single polypeptide **(Supplementary Figure 1C)**. This suggests that α exists predominantly as a dimer in solution. The measured value being lower than the theoretical 158 kDa for a dimer might indicate coexistence of both monomer and dimer species. Native gel electrophoresis supported this conclusion, showing two distinct bands corresponding to monomeric (~80 kDa) and dimeric (~160 kDa) species. A subtle stabilization of the dimer on nucleotide binding is observed **(Supplementary Figure 1D)**. Together, these findings from SEC-MALS and native PAGE demonstrate that the α-subunit exists in a concentration-dependent monomer–dimer equilibrium, which shifts toward dimer formation in the presence of nucleotides.

Unlike NrdF2 that exists as an obligate dimer, α-subunit appears to equilibrate between monomer and dimer form [30]. Earlier it has been shown that the α-subunit of mouse has predominant proportion of monomer which is shown to dimerise in presence of effector nucleotide [31] . Similarly, apo α-subunit from *B. subtilis* was largely present in monomeric form while holo form shifts towards dimeric state [32]. Human RNR is also reported to be monomeric in absence of substrate and effector and dimerization is induced upon effector binding [37]. Our observation of monomer- dimer equilibrium is therefore consistent with that reported by other studies. However, although α-subunit exists in monomer and dimer equilibrium, dimerization is necessary for proper functioning, as loop1 of a monomer is known to govern conformation of loop2 of the second monomer for specificity regulation through correct substrate selection [34, 35].

#### 2.2. Cryo-EM reconstruction of apo and holo α-subunit:

The structure of the α dimer has two-fold symmetry, hence C2 symmetry was imposed for apo structure. Local resolution estimation of the map indicated density at a higher resolution in the inner core compared to the outer flexible regions **(Supplementary Figures 2B and 3D).** Density for most of the residues in the core region was clear, and side chains also could be placed in the density **(Supplementary Figures 5).** Assessment of the heatmap of the angular distribution of particles indicated the presence of preferred orientation bias is more in the apo structure compared to the orientation bias observed in the holo structure **(Supplementary**

**Figure 2D and 3G).** The map-model FSC calculated from Phenix software estimated the resolution to be 4 Å apo α and 7-8.5 Å for the holo classes, respectively, at a threshold of 0.5 **(Supplementary Figure 2E and 3F).**

Rebalance analysis was performed to improve angular distribution and enhance map completeness since the cryo-EM reconstruction lacked N-terminal density and exhibited significant orientation bias. spIsoNet, a self-supervised deep-learning framework was also availed to improve the map quality and map completeness, but with no improvement [36]. Despite this effort, the N-terminal region remained unresolved, suggesting intrinsic flexibility or structural disorder. Even for the holo structure despite better angular coverage, the N-terminal density is missing. These findings indicate that the N-terminus is conformationally dynamic.

**2.3. Flexibility at the N-terminus domain of α subunit:**

To confirm this flexibility in the N-terminal region, limited proteolysis and computational analysis was performed. Compared to control protein (BSA), the purified α fraction shows initiation of degradation within 15 mins of incubation at 37 ºC **(Supplementary Figure 7C).** During the initial stages of limited proteolysis, appearance of a discrete fragment of approximately 18 kDa was observed with decrease in intensity of the intact protein band. Qualitatively the data also reveals that when α is complexed with β or in presence of nucleotide, it appears to be relatively resistant to degradation. The presence of degraded band at ~18kDa indicates cleavage towards either N or C termini. This sensitivity of α-subunit at such a low concentration of protease and short time indicates it has regions near termini that are flexible and have a site that is accessible to protease cleavage. Earlier molecular simulation study suggested that more than 10 residues around the cleavage site should be unfolded to provide accessibility to trypsin for cleavage [37].

The ~18 kDa band released on limited proteolysis was excised and analysed by in-gel trypsin digestion followed by QTRAP mass spectrometry **(Supplementary Figure 7D).** Peptide mapping with minimum of 4 amino acids containing peptide and the complete digestion indicated that majority of identified peptides corresponded to the N-terminal region of α-subunit **(Supplementary Figure 7E).** The analysis of peptide obtained indicate loop 1, which is unstructured effector binding site and loop 3 region are among the sites of cleavage. The stability imparted in loop1 due to nucleotide binding may explain the relative resistance of α-subunit in the presence of the nucleotide.

Sequence and structure analysis was performed to understand this flexibility of the N-terminus of α. IDPs typically have higher net charge and lower hydropathy as compared to folded proteins. Comparison of charge-hydropathy of α-subunit with IDPs indicates it to be folded **(Supplementary Figure 7F).** The PONDR plot predicts the probability of a protein region being intrinsically disordered, with regions of high score being disordered and low score as ordered. Although the α-subunit is ordered, the region near residue 150 has a stretch of disordered residue **(Supplementary Figure 7G).** Charge-hydropathy and PONDR plots suggested α-subunit as an ordered protein but with a region of flexibility in the N-terminus [38]. Medusa based on evolutionary information and physico-chemical properties predicts flexibility in the N-terminal region and loop regions [39].

In summary, the collective data underscored the presence of a flexible region in the N-terminal segment of α, likely playing a significant role in its function.

#### 2.4. Normal mode analysis:

In the correlation plot the N-terminal region of one chain is majorly negatively correlated with the core domain of the same chain and the other chain. This region also shows slight to intermediate positive correlation with the N-terminal region of the other chain in the dimer **(Supplementary Figure 11).** With respect to this, one of the modes clearly highlights the unidirectional motion of the N-terminal region towards the two-fold symmetry axis, which eventually leads to synchronous movement of the other N-terminal region away from the two-fold symmetry axis. This further substantiates the anticorrelated movement observed in the α-subunit as reported by 3D variability analysis.

#### 2.5. Resolving continuous flexibility and variability in α-subunit:

3-dimensional variability analysis (3DVA) of the apo α-subunit Cryo-EM dataset using consensus refinement of 126K particle images resolved three major classes of heterogeneity **(Supplementary Figure 12).** The scatter plot showed maximum variability along components 0 and 1, indicating continuous conformational heterogeneity within the sample **(Supplementary Figure 12A).** The first variability component corresponded to twisting and compression motions, predominantly localized to the N-terminal region **(Supplementary Figure 12B).** The second component revealed bending motion along the two-fold symmetry axis, resulting in pronounced N-terminal density changes and reflecting intrinsic flexibility within the dimer **(Supplementary Figure 12C).** The third component showed lateral displacement of the α monomers relative to the two-fold symmetry axis **(Supplementary Figure 12D)**.

Similarly, 3DVA of the holo α-subunit dataset (merge class with effector and substrate holo1-holo4) using 370K particle images also resolved three major classes of heterogeneity, with maximum variability observed along components 1 and 2, indicating continuous conformational heterogeneity within the sample **(Supplementary Figure 14A).** The first component revealed inward movement of the N-terminal region toward the two-fold symmetry axis, directed toward the loop 1 region of the other monomer. The second component revealed upward displacement of the N-terminal region toward the active site **(Supplementary Figure 14B–D).** The third component showed movement toward the two-fold symmetry axis and upward movement toward the active site. In the holo structure, since density is present in the N-terminal substrate-bound chain, movement of the N-terminal toward the symmetry axis and also toward the active site is seen.

Notably, both apo and holo structures exhibited synchronous movement between loop 2 and the N-terminal region, bringing these regions into closer proximity. This anticorrelated breathing motion alternated between the two monomers of the dimer, suggesting a coordinated opening-and-closing mechanism of the active sites that may facilitate substrate docking in one protomer concomitant with product release in the other **(Supplementary Video 5).**

**2.6. Comparison of α-subunit with other known structures:**

The superposition of Mth α-subunit with reported structures shows high structural similarity with both Class Ia and Ib RNRs, except for the longer N-terminal cone domain present in Class Ia. **Supplementary Table 2** shows cross structure statistics among different reported structures. Superposition of Mth α dimer with structures of Class Ib α-subunit of *Salmonella typhimurium* (PDB ID: 1PEM) and that of *Bacillus subtilis* (PDB ID: 6MVE) yielded RMSD of 2.6 Å and 3 Å respectively **(Supplementary Figure 15A).** The sequence identity of Mth α-subunit was higher with *S. typhimurium* (72.4%) than *B. subtilis* (49.3%). Similarly, comparison with the Class Ia structure of *E. coli* (PDB ID 2R1R) yielded an RMSD of 3.8 Å and minimal sequence identity of 26.9% **(Supplementary Figure 15B and 15C).** As expected, the RMSD for the dimer of *B. subtilis* (PDB ID: 6CGL and 6CGN) was above 20 Å **(Supplementary Figure 15D),** as these structures were determined in inhibitory conditions leading to formation of non-canonical dimer [40].

**Supplementary Figures and Legends:**

**Supplementary Figure 1:**

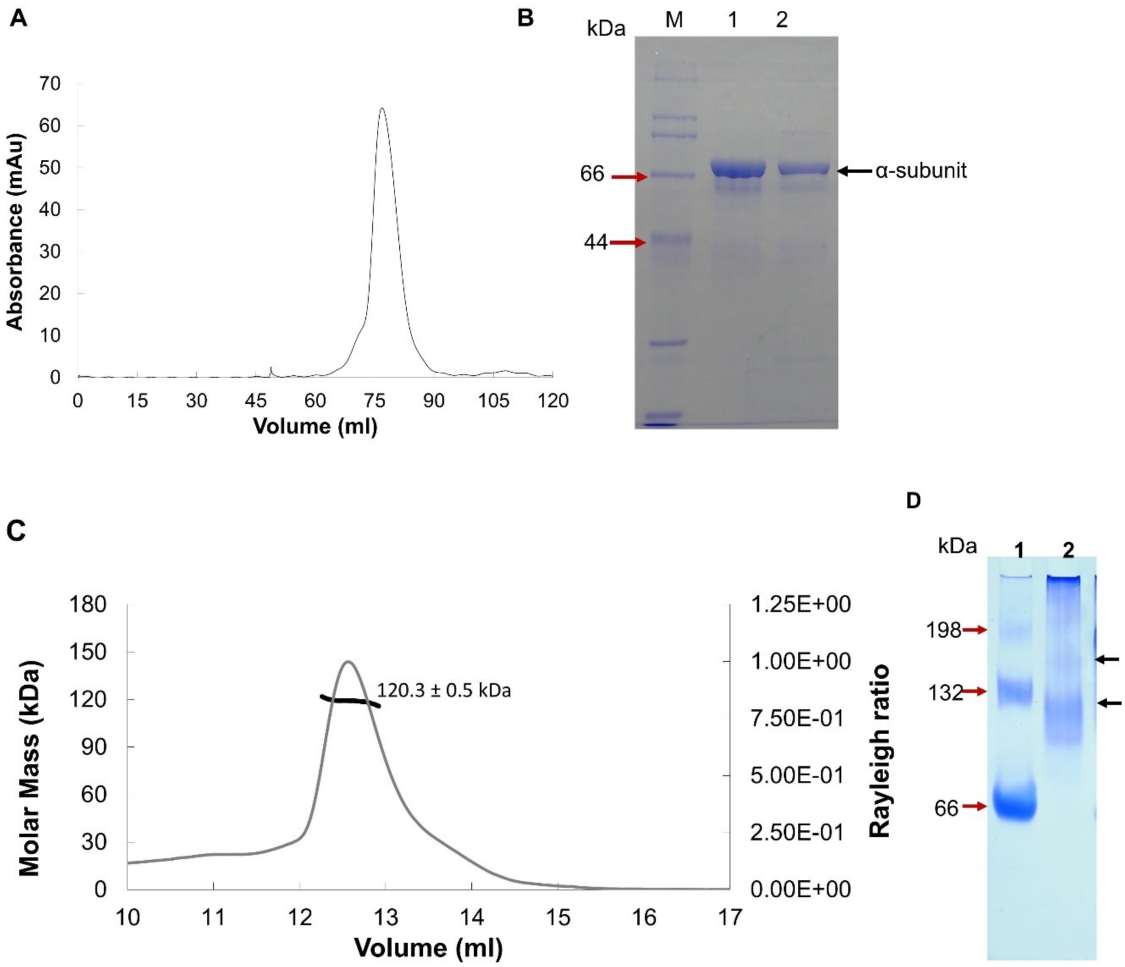


**Supplementary Figure 1: Oligomeric characterization of Mth α subunit:**

**A.** Size exclusion chromatography profile of purified α-subunit with a Superdex 200 16/600 column. **B.** SEC purified fraction loaded on 12% SDS gel showing sample purity. **C.** SEC MALS analysis of α-subunit in Superdex 200 10/30 column. **D.** Native gel electrophoresis in 8% gel: lane1-BSA, 2- α-subunit. Red arrow indicates monomer, dimer and trimer species of BSA while black arrow indicates monomer and dimer species of α-subunit

**Supplementary Figure 2:**

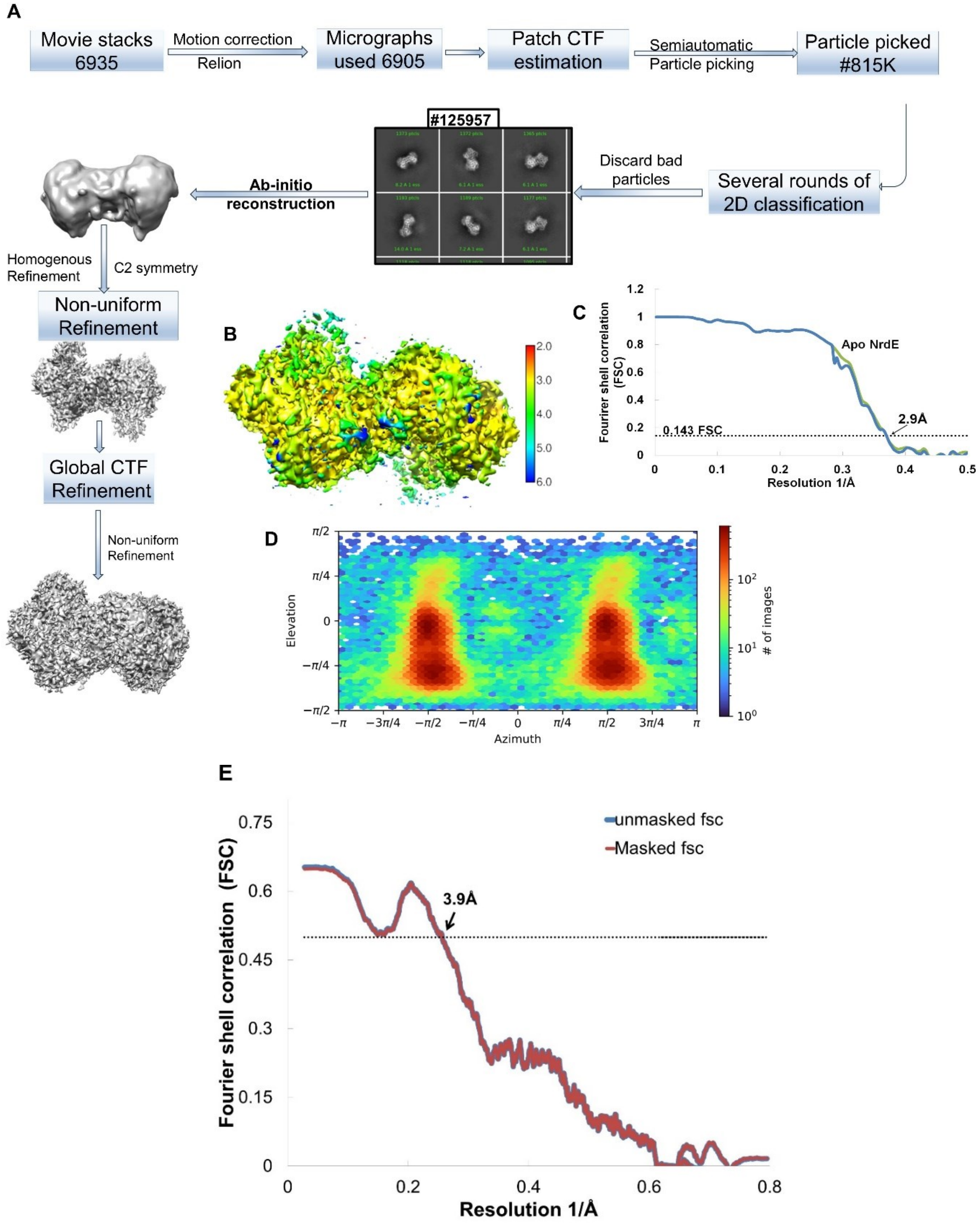


**Supplementary Figure 2: Single-particle cryo-EM processing workflow and reconstructions of the Mth α-subunit in apo form:**

**A.** Flow chart showing the image-processing workflow for the cryo-EM data of apo α-subunit.

**B.** Local resolution estimation with the color scale bar shown on its right indicates resolution in Å. As can be seen, in most parts of the core structure, resolution is close to 3 Å, whereas on the periphery of the structure, the resolution is poor. **C.** The "gold standard" FSC between two independent halves of the map. The dotted line represents the 0.143 FSC cut-off, which indicates a nominal resolution of 3.0 Å calculated using CryoSPARC mask (where green (FSC with tight mask) blue (FSC with noise substitution). **D.** Angular distributions calculated in CryoSPARC for particle projections of all particles used for the final three-dimensional reconstruction. The heatmap shows the number of particles for each viewing angle. **E.** Map model of the FSC curve of apo α-subunit with (red) and without mask (blue). The dotted line represents the 0.5 FSC cut-off, which indicates a resolution of 3.9 Å obtained from Phenix real space refinement.

**Supplementary Figure 3:**

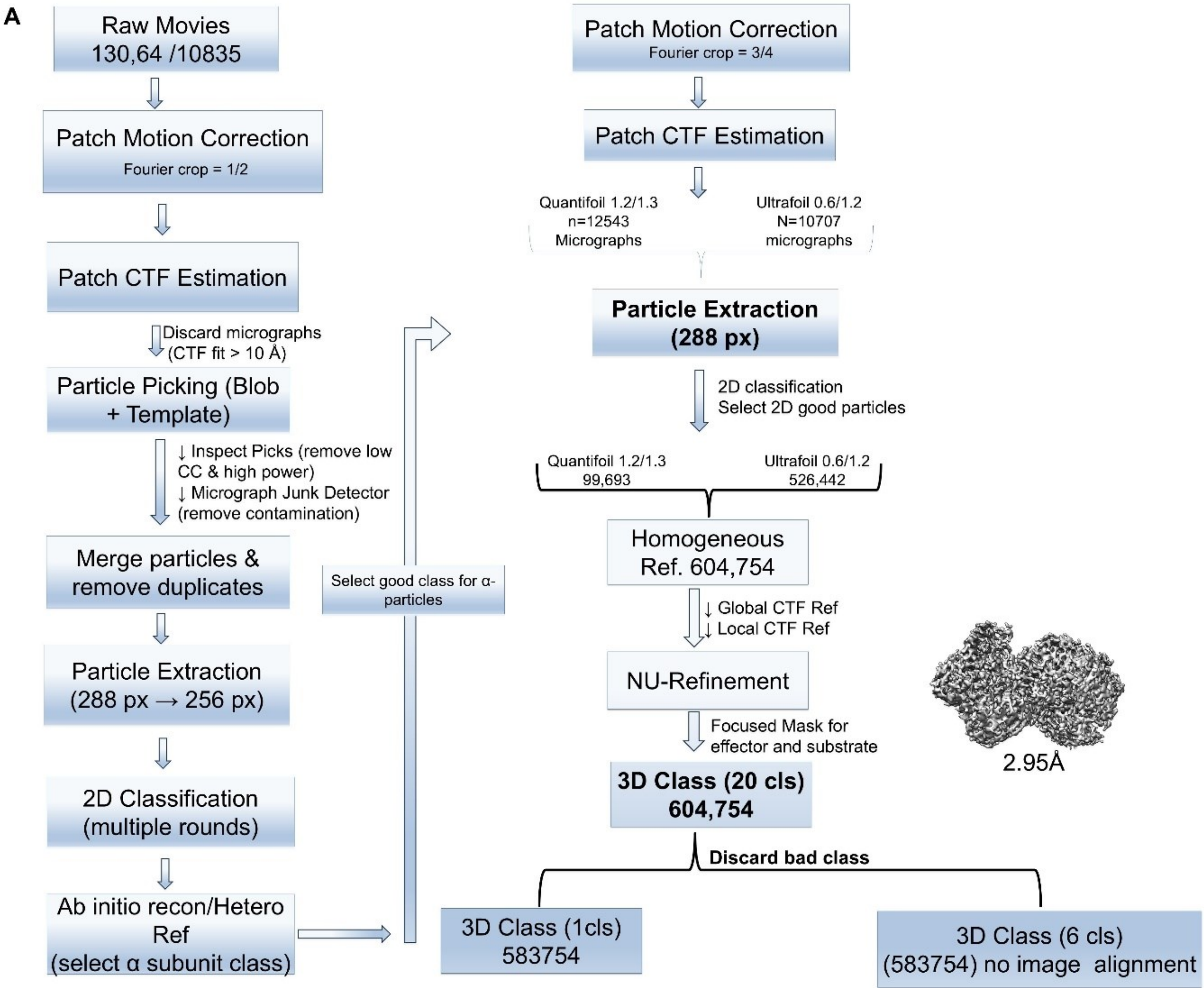

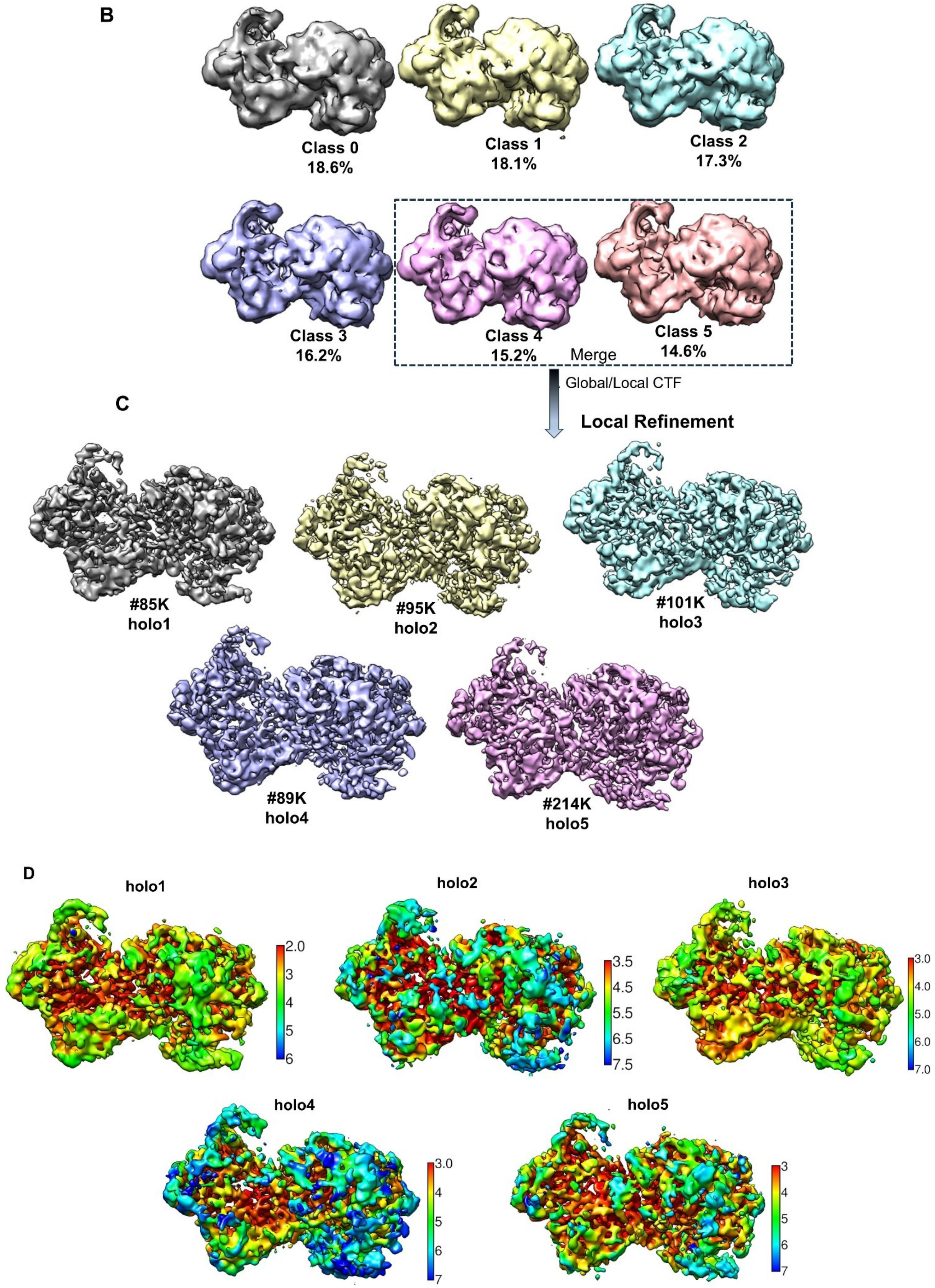
B
Class 0
18.6%
Class 1
18.1%
Class 2
17.3%
Class 3
16.2%
Class 4
15.2%
Class 5
14.6%
Merge
Global/Local CTF
C
Local Refinement
#85K
holo1
#95K
holo2
#101K
holo3
#89K
holo4
#214K
holo5
D
holo1
holo2
holo3
holo4
holo5

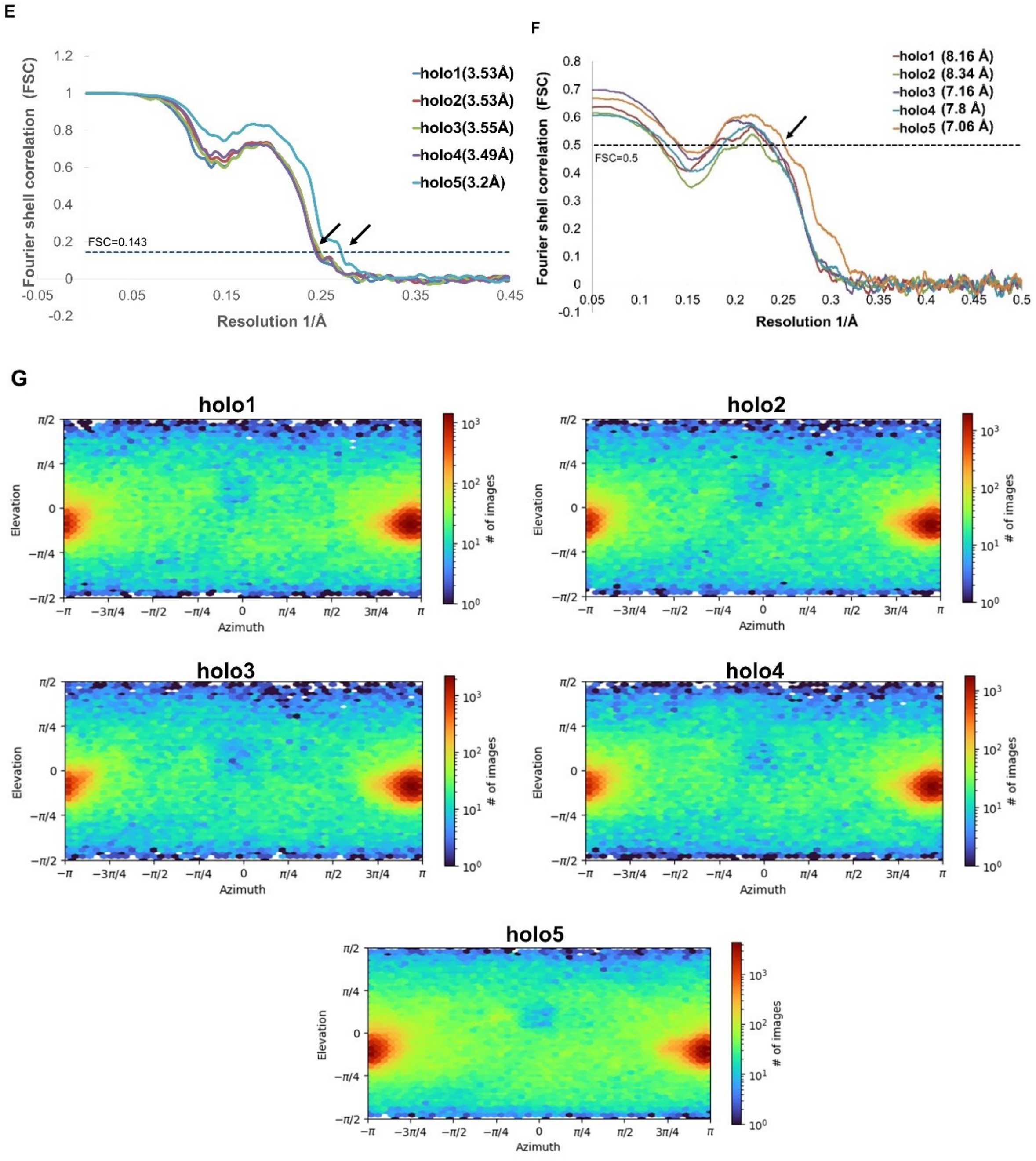


**Supplementary Figure 3: Cryo-EM structure determination workflow of Mth α-subunit in the holo form with TTP and GDP:**

**A.** Flow chart showing the image-processing pipeline for the cryo-EM data of the holo α-subunit. **B.** Six unsharp map of 3D classes obtained without alignment after focused classification. **C.** CryoSPARC sharpened map using a B-factor of −40 Å² for all final five 3D classes used for structure determination. **D.** Local resolution estimation of all 3D class after high resolution refinement with the color scale bar shown on its right indicates resolution in Å. **E.** The “gold standard” FSC between two independent halves of the map indicates a resolution 3.22-3.55Å. The dotted (black) line represents the 0.143 FSC cut-off mask with resolution of each of the 3D classes is mentioned in the bracket. **F.** Map model FSC curves of holo α-subunit for all 3D classes indicates resolution in Å as shown in bracket as obtained from Phenix real space refinement. The dotted line represents the resolution at 0.5 FSC cut-off. **G.** Angular distributions calculated in CryoSPARC for particle projections of all particles used for the final three-dimensional reconstruction. Heatmap shows the number of particles for each viewing angle for each of the 3D class.

**Supplementary Figure 4:**

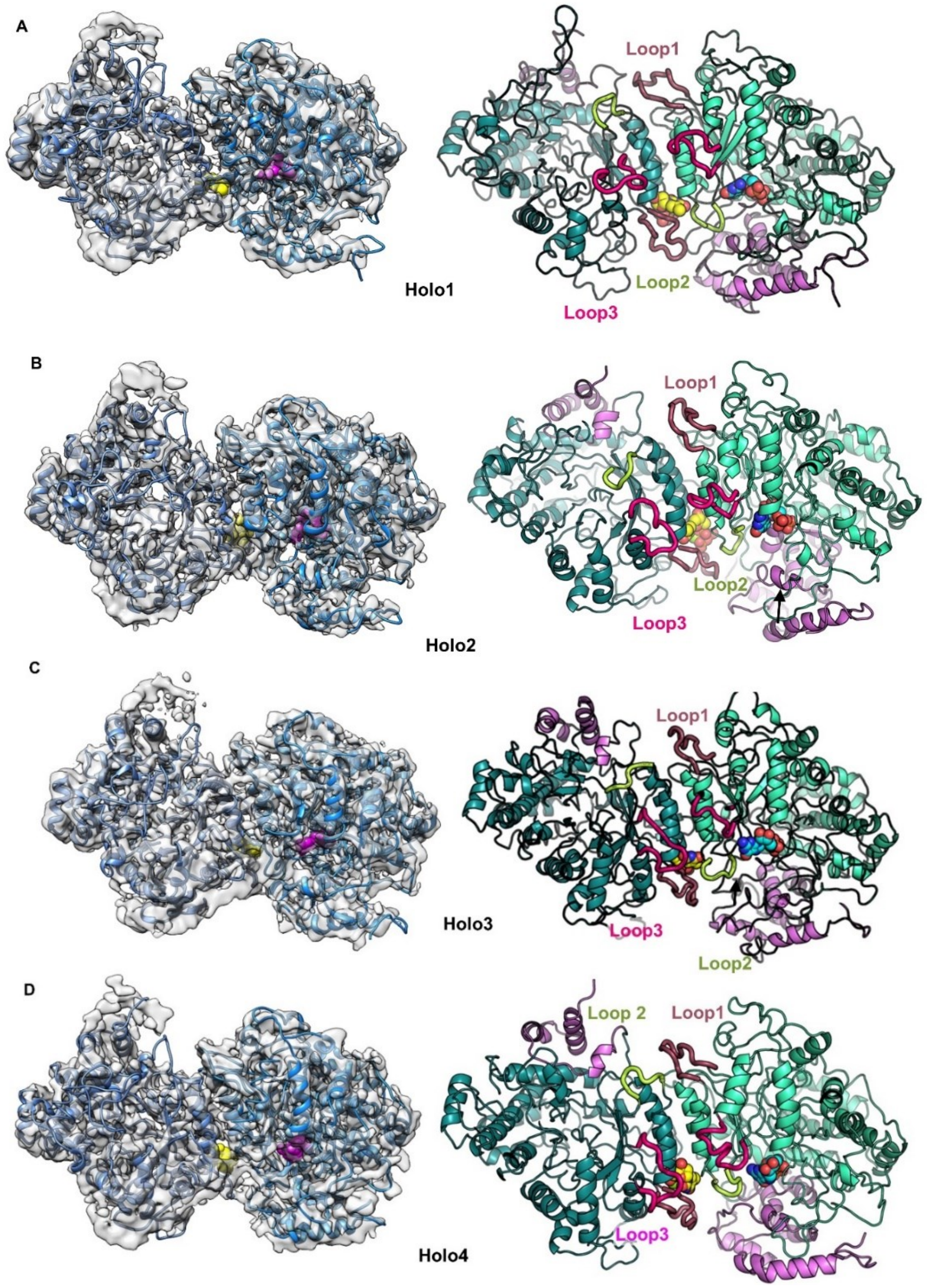

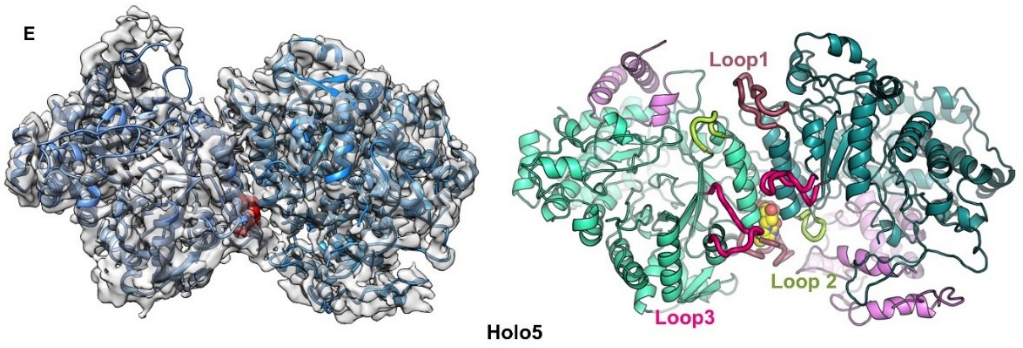


**Supplementary Figure 4: Overview of structures of different 3D class of Mth α-subunit in holo form:** Map-model overlay (left panel) and structure (right panel) of these five-3D class (holo1-holo5) of holo α-subunit is shown **(A-E):** A CryoSPARC sharpened map shown as a transparent surface with a model fitted colored in blue. The map contour is 0.13 as displayed in Chimera. The effector site and active site regions are shown in yellow and blue spheres respectively for the model. The N-terminal region is shown in violet colour with ~100 residue missing in effector bound chain. Loop1, 2 and 3 is highlighted in brown, green, and pink color respectively.

**Supplementary Figure 5:**

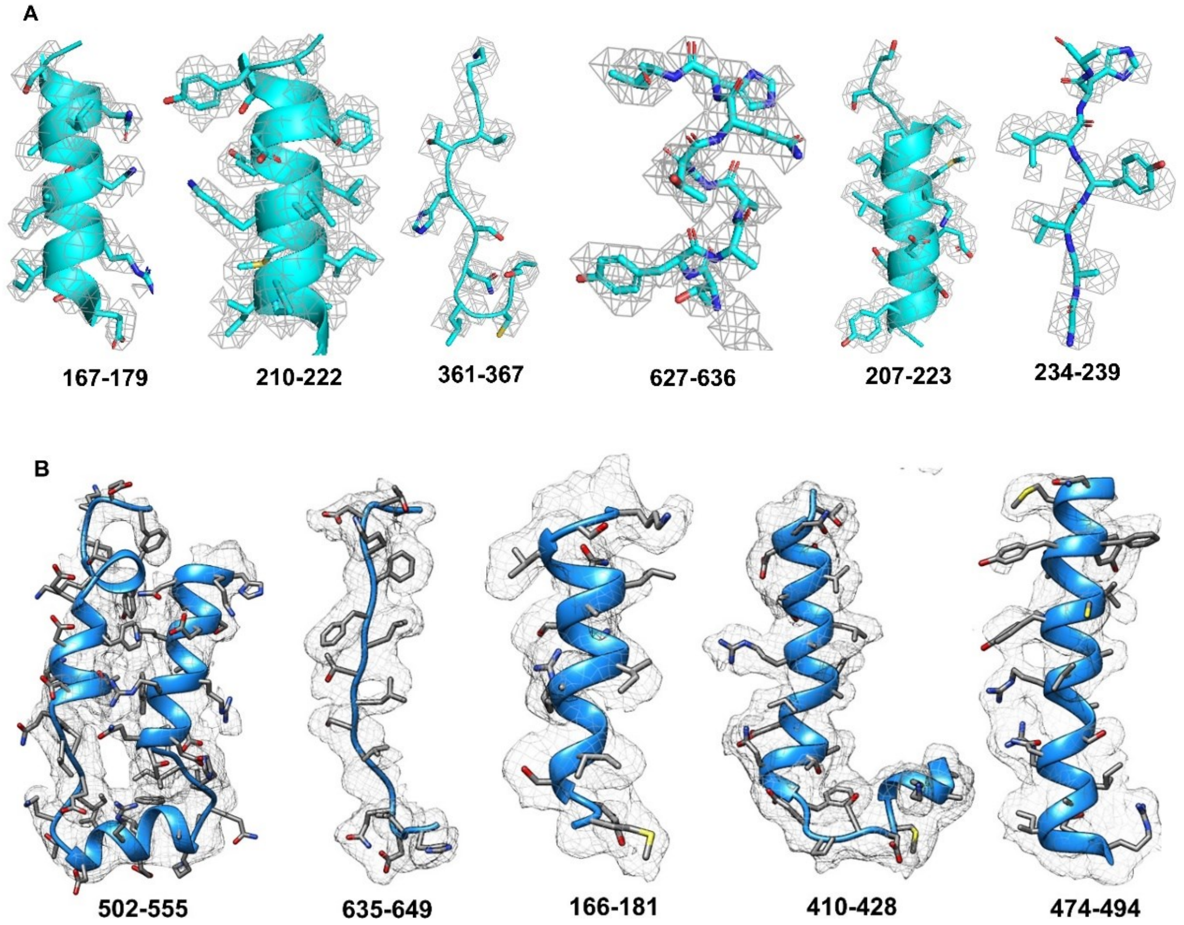

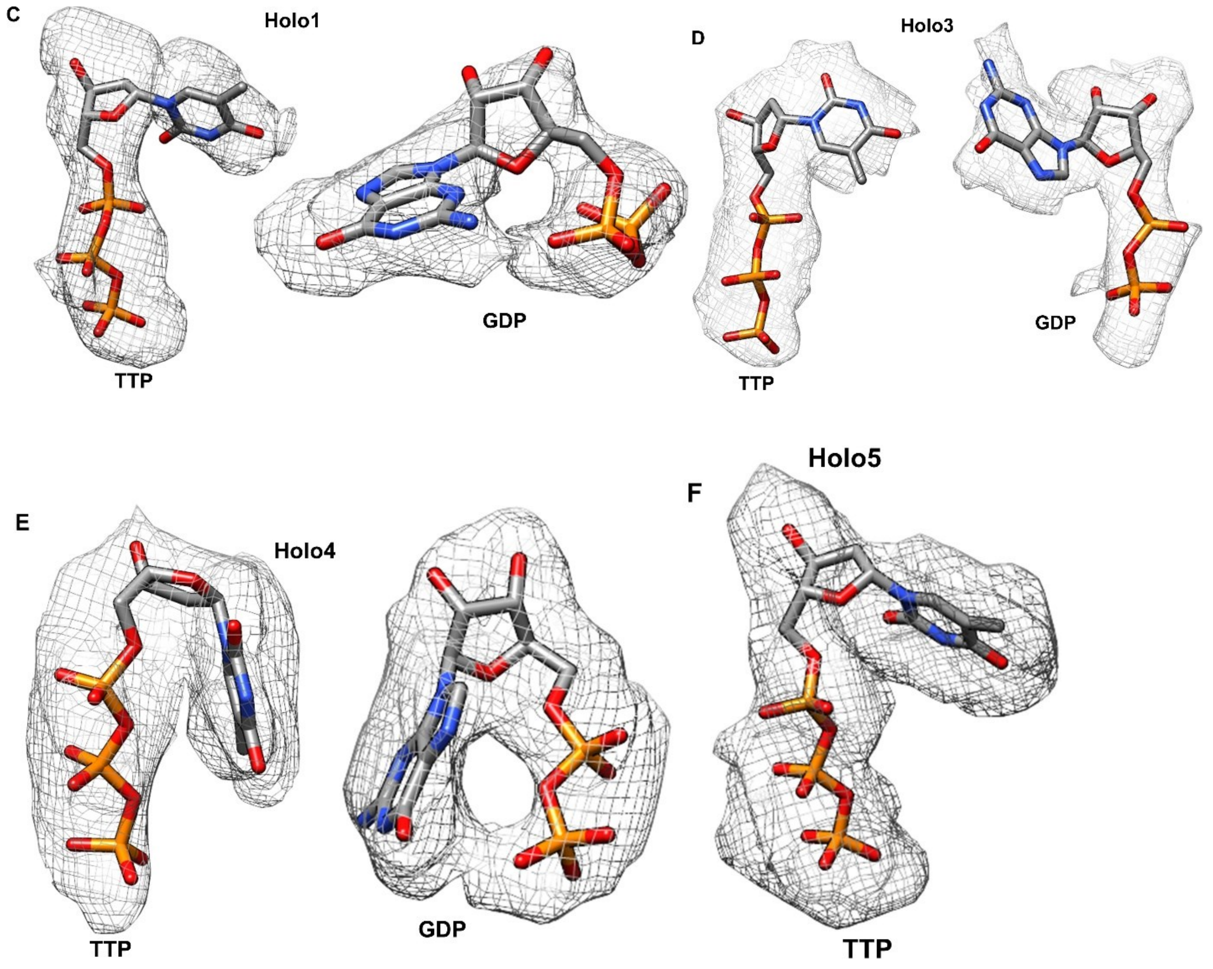


**Supplementary Figure 5: Representative local cryo-EM density:** Local cryo-EM density with structure, as visualized in UCSF Chimera is shown as cartoon and side chain shown as sticks. Representative regions of the α-subunit map (grey surface) demonstrating side chain density in local regions with residue range listed below the image**.** As demonstrated, the main chain as well as the side chains fit well in the experimental density in most parts of the structure **A.** The DeepEMhancer sharpened map of apo α-subunit was contoured at 0.01 σ. The different local map was contoured at level 1e-05**.** Color zone tool of Chimera with a radius cutoff of 1 Å was used to map the density to the atomic model. The cut-off radius was chosen to provide an optimal balance between specificity and completeness in mapping density around the atomic coordinates. **B.** The CryoSPARC sharpened map of holo3 α-subunit contoured at level 0.1 σ. The different local maps were contoured at level ranging from 0.09-0.13. Color zone tool of Chimera with a radius cutoff of 2 Å was used to map the density to the atomic model.

CryoSPARC-sharpened maps were used to visualize the densities corresponding to TTP and GDP in different classes of the holo α-subunit. For map visualization the Color Zone tool in

Chimera with a radius cutoff of 2 Å was used. Subtraction of the protein-only map from the original sharpened map yielded the nucleotide difference density used for visualization of bound nucleotides **C.** Holo1 density for TTP and GDP nucleotide contoured at map level 0.05 and 0.15 respectively, **D.** Holo3 density for TTP and GDP nucleotide contoured at map level 0.1, **E.** Holo4 density for TTP and GDP nucleotide contoured at map level 0.16, **F.** Holo5 density for TTP contoured at map level 0.07

**Supplementary Figure 6:**

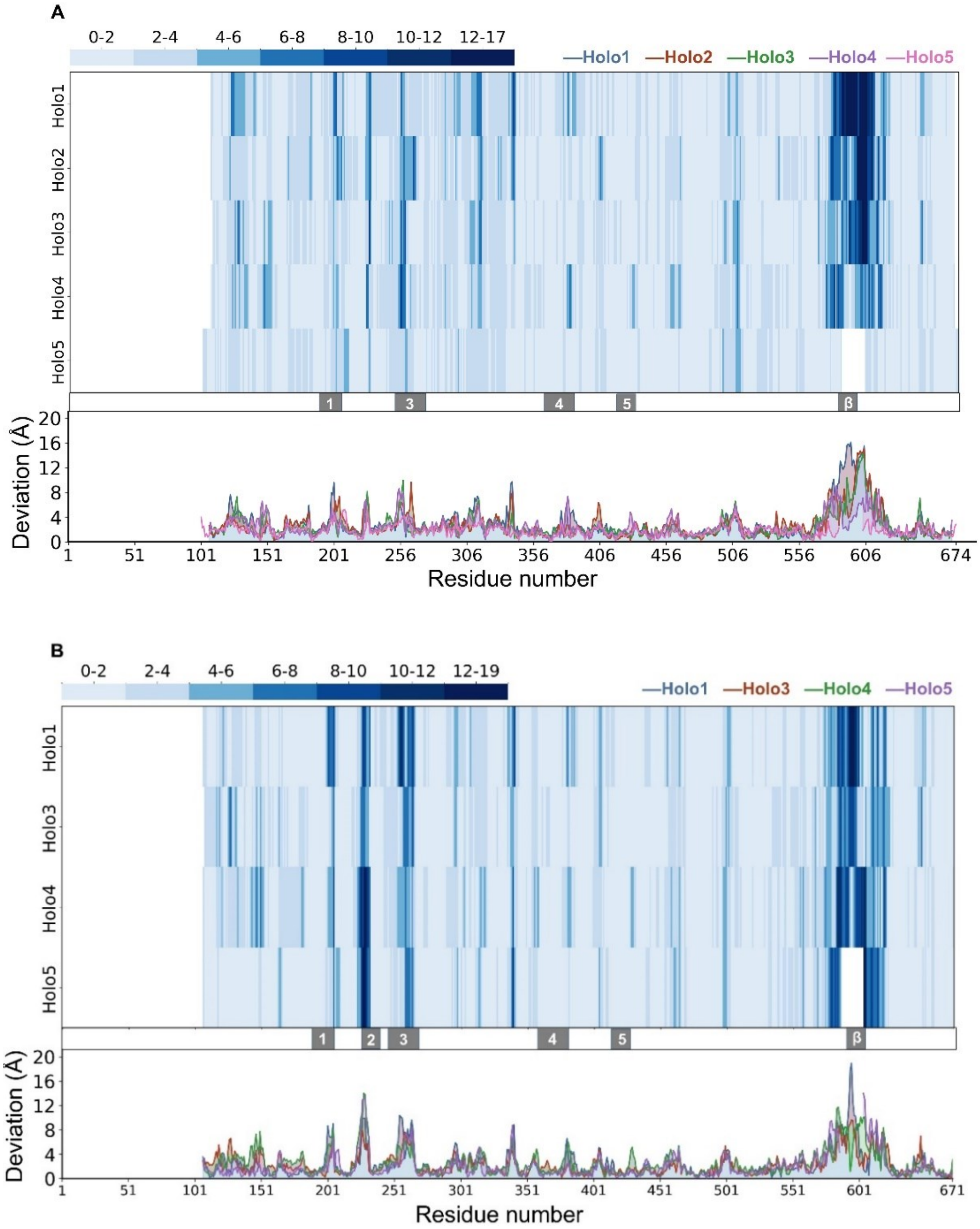

**Supplementary Figure 6: Per residue deviation of effector-bound chain of holo structures.** The blue, red, green, purple and pink lines in the line-plot corresponds to Holo 1, Holo 2, Holo 3, Holo 4 and Holo 5 respectively. The blue color bar at the top shows deviation range of the heatmap in Å. The grey regions labelled as 1, 2, 3, 4, 5 and β represents Loop 1, Loop 2, Loop 3, Loop 4, Loop 5 and β-hairpin loop regions respectively across the x-axis showing residue number. **A.** Per residue deviation of effector-bound chain of holo structures w.r.t. chain A of apo structure. This plot does not include Loop 2 region (226-230) as it was missing in apo NrdE. **B.** Per residue deviation of effector-bound chain of holo structures w.r.t. reference: Holo2 structure. White color space indicates missing region in the model.

**Supplementary Figure 7:**

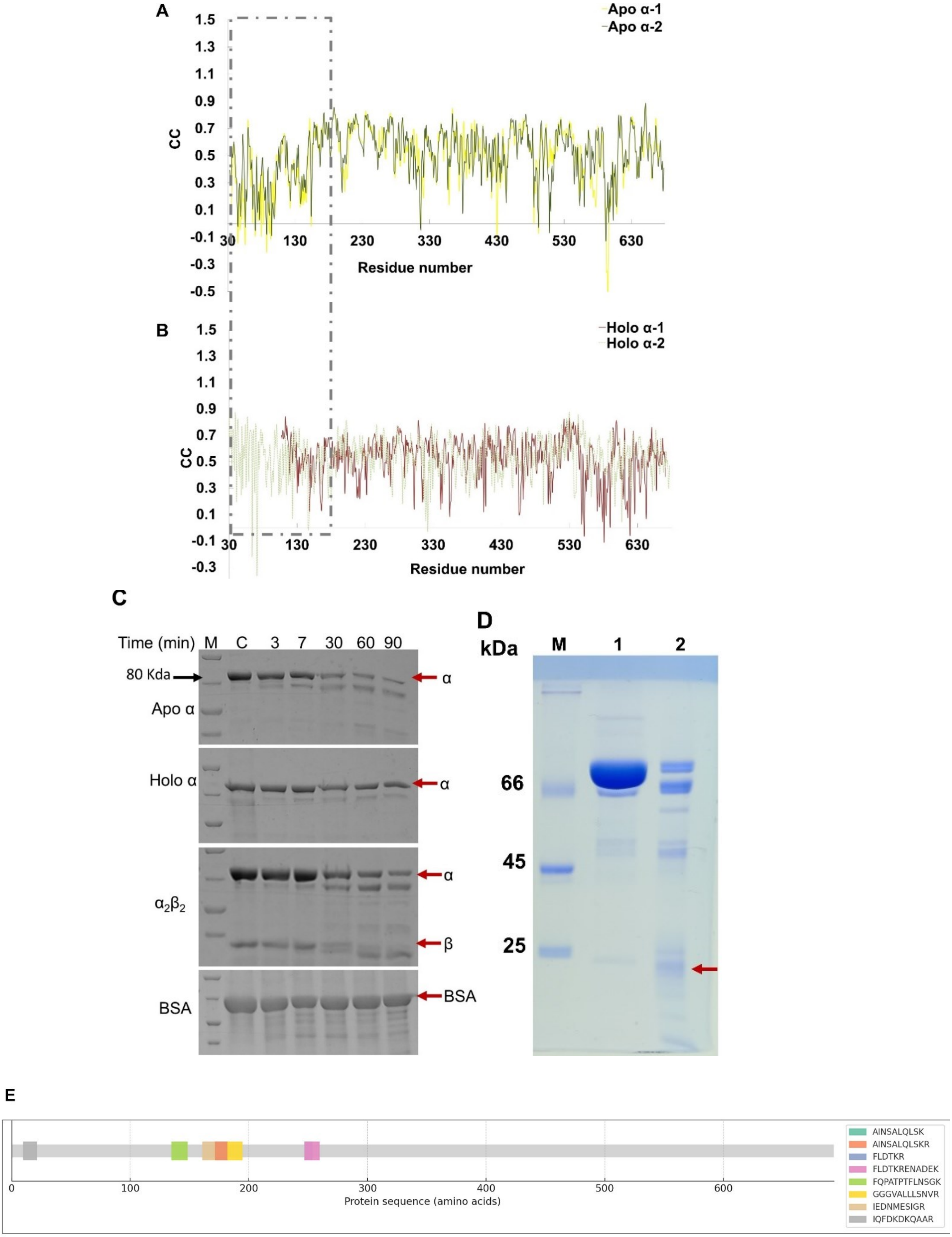

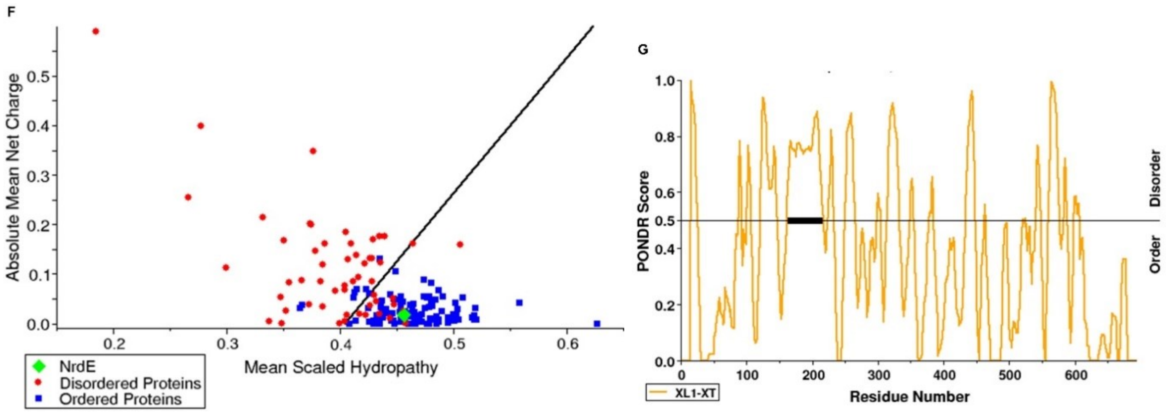


**Supplementary Figure 7: Flexibility in the N-terminus of α-subunit:**

Correlation graphs showing the average correlation coefficient (CC) between the model and map, with regions of low correlation at the N-terminus highlighted by dotted boxes. For the apo α-subunit, the DeepEMhancer-sharpened map was used for refinement, whereas for the holo α-subunit, the cryoSPARC-sharpened map was used. **A.** Correlation between the two monomers of the apo structure, indicating poor correlation at the N-terminus.**B.** Correlation between the substrate-bound and effector-bound monomers of the holo structure, showing absent density at the N-terminus of the effector-bound (red) and the presence of N-terminal density in the substrate-bound monomer (green). **C.** Limited proteolysis of α-subunit was performed under different condition mentioned and the sample was electrophoresed on 10% SDS gel. **D.** Trypsin digestion of α-subunit for 0 and 90 minutes to obtain ~18 kDa band for mass spectrometry as indicated by red arrow. **E.** Protein sequence coverage by mass spectrometry**.** The grey bar represents the full-length protein sequence with different peptides hits obtained highlighted in different colours and the sequence of peptides shown in the right of the image. **F.** Charge-hydropathy plot with average net charge of a protein against its mean scaled hydropathy. **G.** PONDR plot**.** Black thick line around residues no 200 highlights the long disorder stretch that may be susceptible to degradation.

**Supplementary Figure 8:**

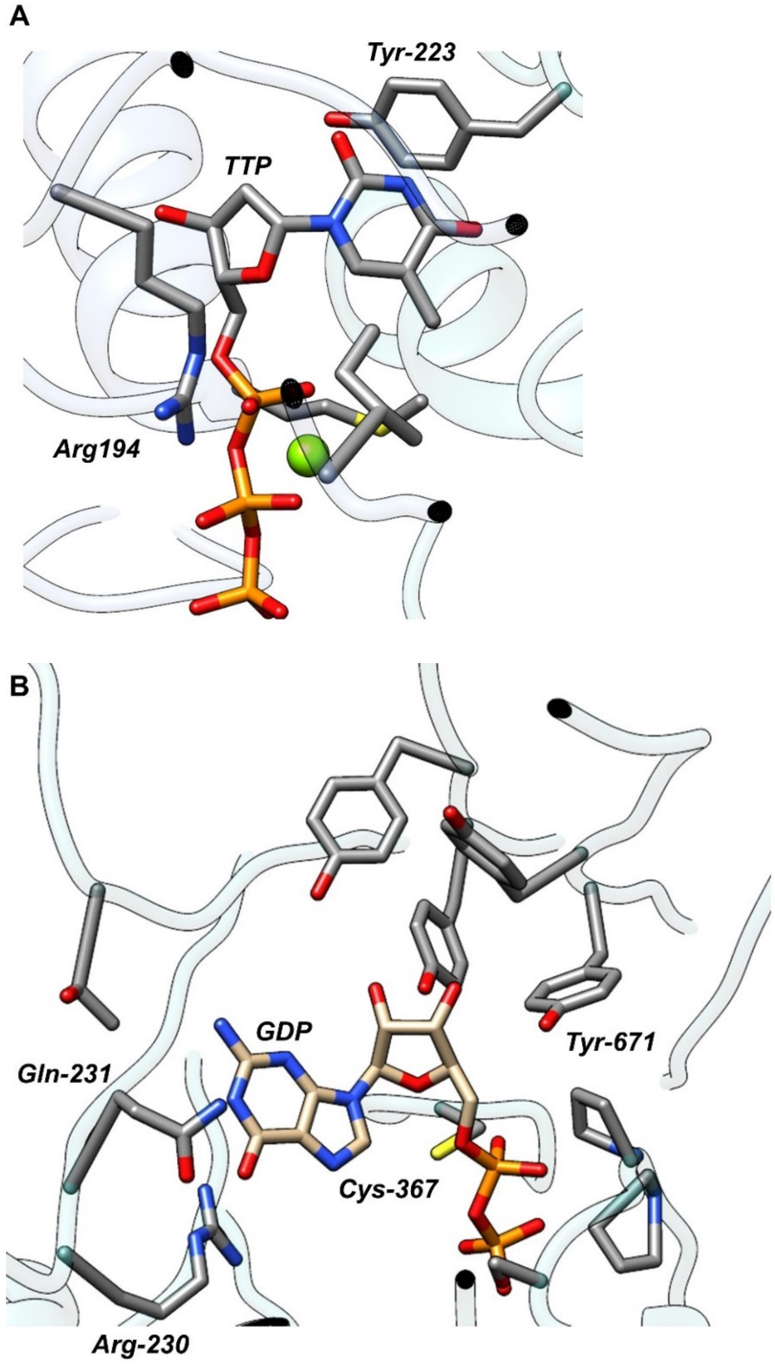


**Supplementary Figure 8:**

**Active and effector sites**.: **A.** Effector TTP bound at the allosteric site. **B.** Substrate GDP bound at the catalytic site in a representative holo3 class structure. The protein is shown in cartoon representation, with residues in the vicinity of the bound ligands displayed as sticks.

**Supplementary Figure 9:**

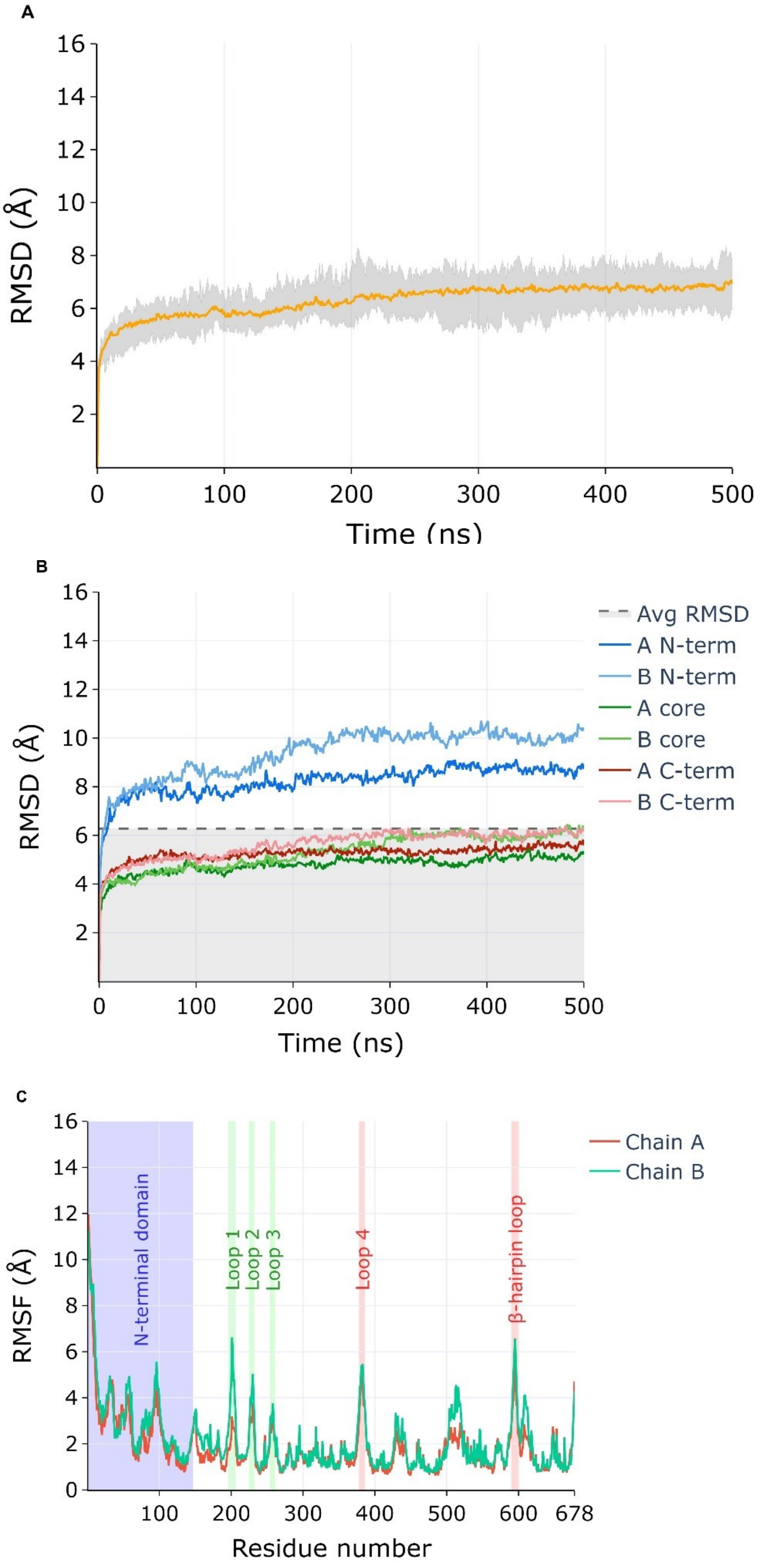

**Supplementary Figure 9: Fluctuations observed during 500 ns of MD simulation:**

**A.** The average RMSD of α dimer is represented at different time frames. The grey region depicts the maximum and minimum RMSD at each time frame across the replicates, **B.** The domain specific RMSD for N-terminal (blue), core (green) and C-terminal (red) domain over the simulation time. The lighter shades of respective colours represent chain A whereas darker shades represent chain B of the dimer. The dotted grey line marks the average RMSD of the dimer, **C.** The root mean square fluctuations of residues of chain A (orange) and chain B (green) of the α dimer. The region marked in red highlights major fluctuating residues (Loop 4 and β-hairpin loop) while regions marked in blue and green represent the N-terminal domain and loop 1, 2 and 3 respectively.

**Supplementary Figure 10:**

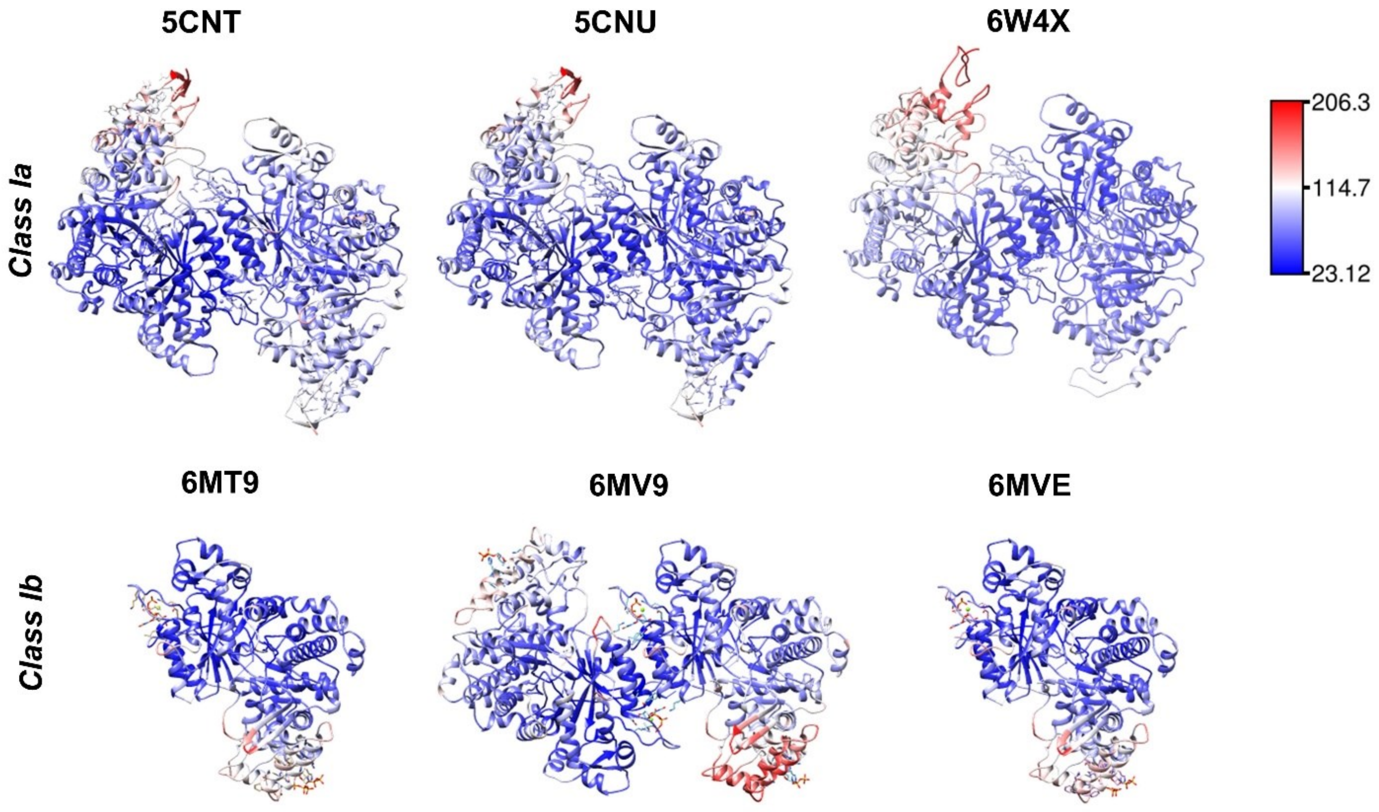


**Supplementary Figure 10: Disordered N-terminal depicted by high B-factors:**

The average B factor as a function of residue for Class Ia (PDB ID 5CNT, 5CNU, 6W4X) and Class Ib *B. subtilis (*PDB ID 6MV9, 6MT9, 6MVE). The colour bar indicates the B factor value from blue to red.

**Supplementary Figure 11:**

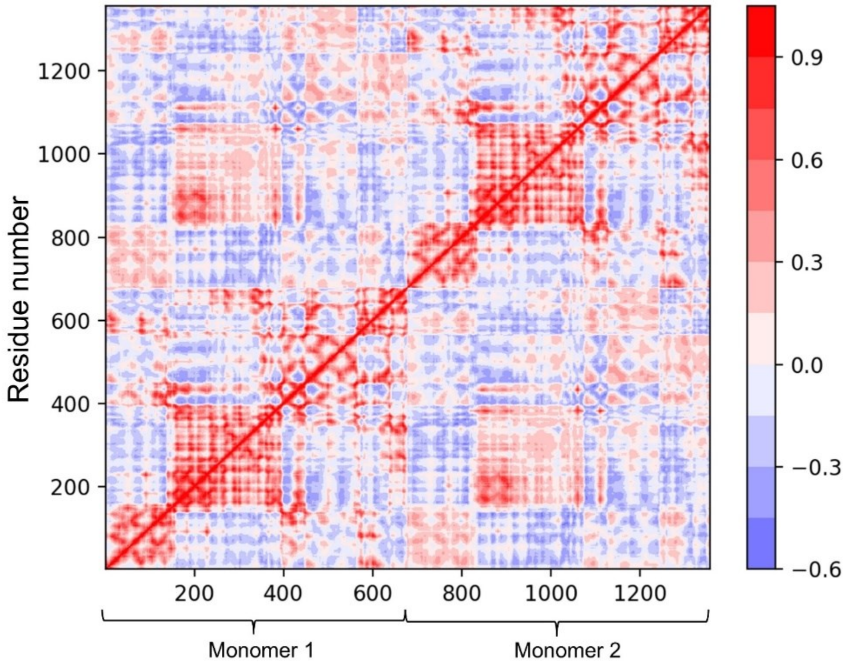


**Supplementary Figure 11: Correlations deduced from Normal Mode Analysis:**

The plot shows positive (red) and negative (blue) correlations between residues as obtained after normal mode analysis. The residue numbers range from 1 to 1356 (1-678 representing monomer 1 and 679-1356 representing monomer 2).

**Supplementary Figure 12:**

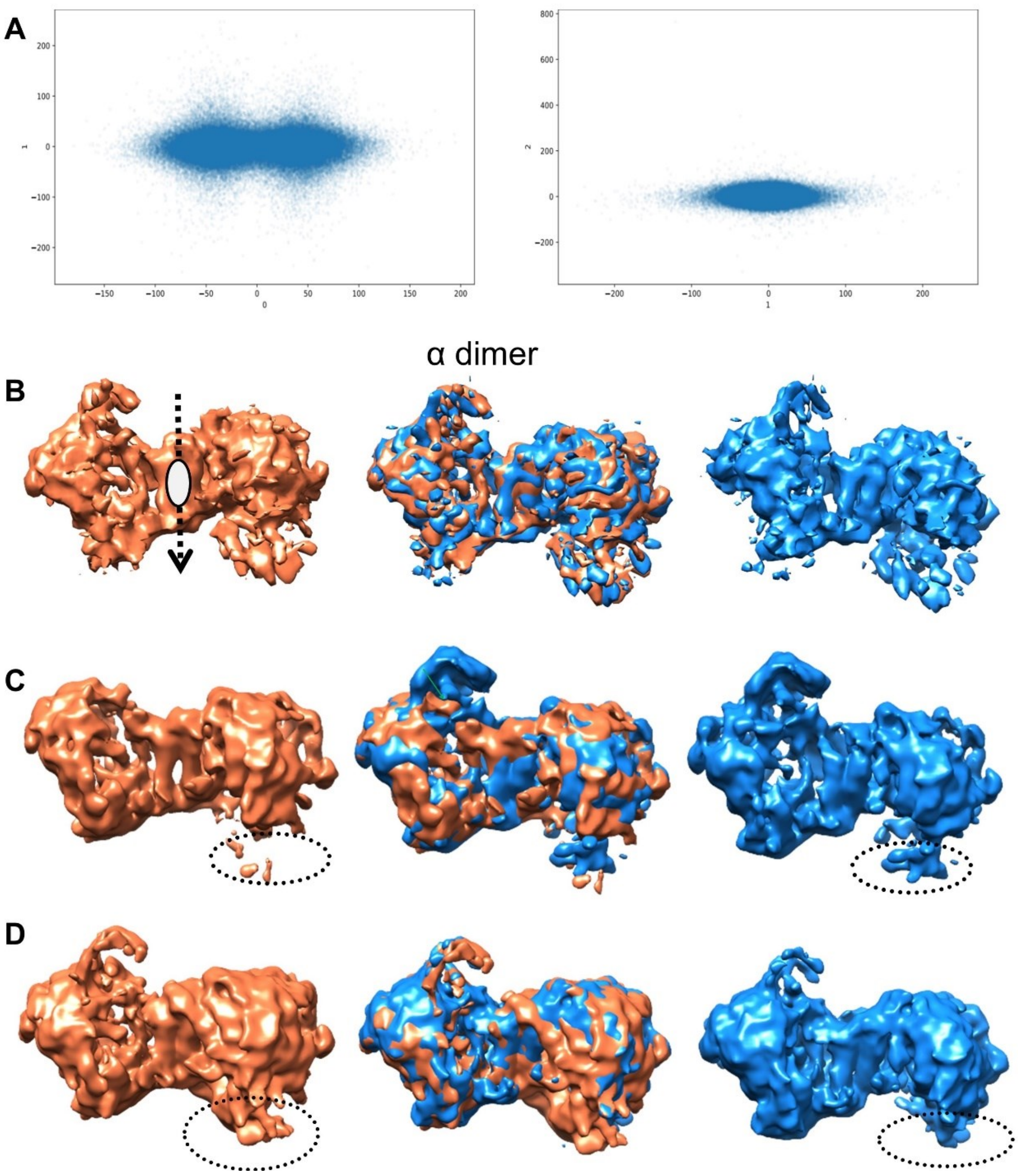


**Supplementary Figure 12: 3D Variability analysis of the apo α subunit:**

**A.** Scatter plot obtained indicated maximum variability along components 0 and 1 present in the sample. **B.** Variability Component 0. The identified 2-fold axis is indicated by the white ellipsoid, **C.** Variability Component 1, and **D.** Variability Component 2. The dotted circle indicates variability observed in the N-terminal region of α subunit.

**Supplementary Figure 13:**

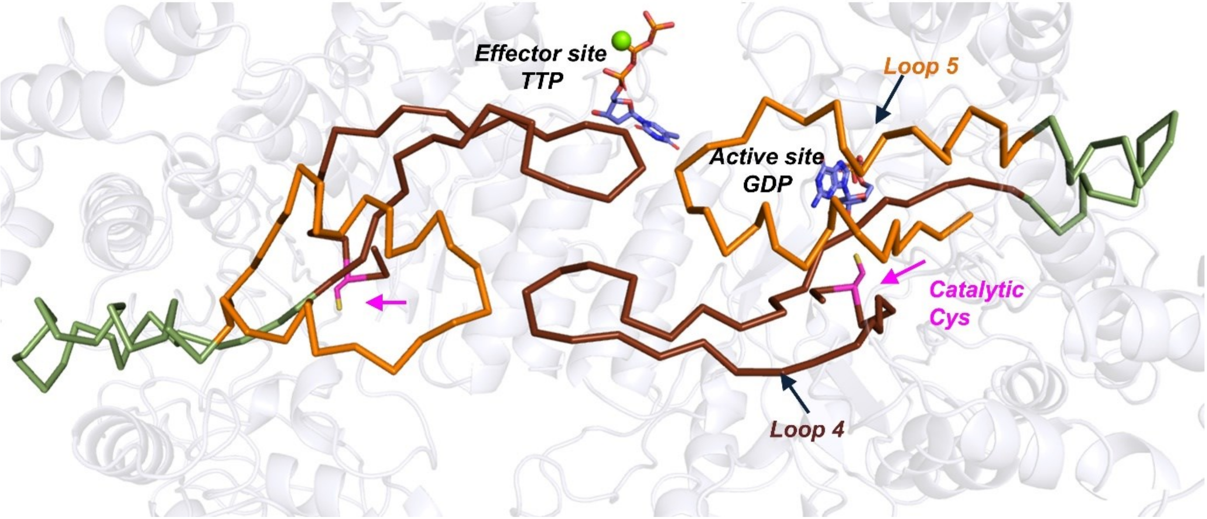


**Supplementary Figure 13: Seesaw motion in the dimer of α dimer**

Anticorrelated movement was observed in loop4 of one monomer and loop5 of another monomer as observed in 3DVA. Ribbon representation of loop 4 is shown in brown, loop 5 in orange, and the connecting region between loops 4 and 5 is shown in green. The catalytic residue is depicted in pink and indicated by a pink arrow.

**Supplementary Figure 14:**

A

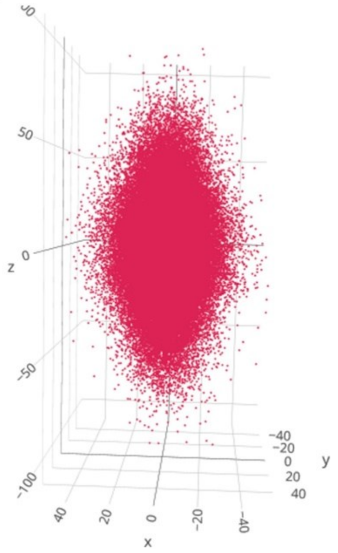

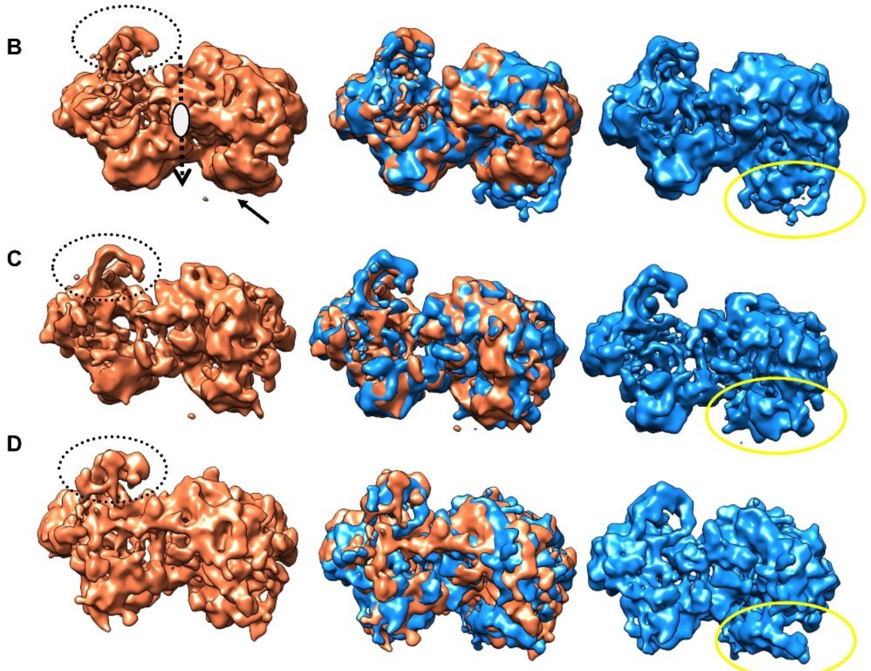


**Supplementary Figure 14: 3D Variability analysis of the holo α subunit:**

A. Scatter plot obtained indicated maximum variability along components 1 and 2 present in the sample. B. Variability Component 0. The identified 2-fold axis is indicated by the white ellipsoid, C. Variability Component 1, and D. Variability Component 2. The black dotted circle indicates variability observed in the N-terminal region of effector bound α subunit. The yellow circle indicates variability observed in the N-terminal region of substrate bound α subunit.

**Supplementary Figure 15:**

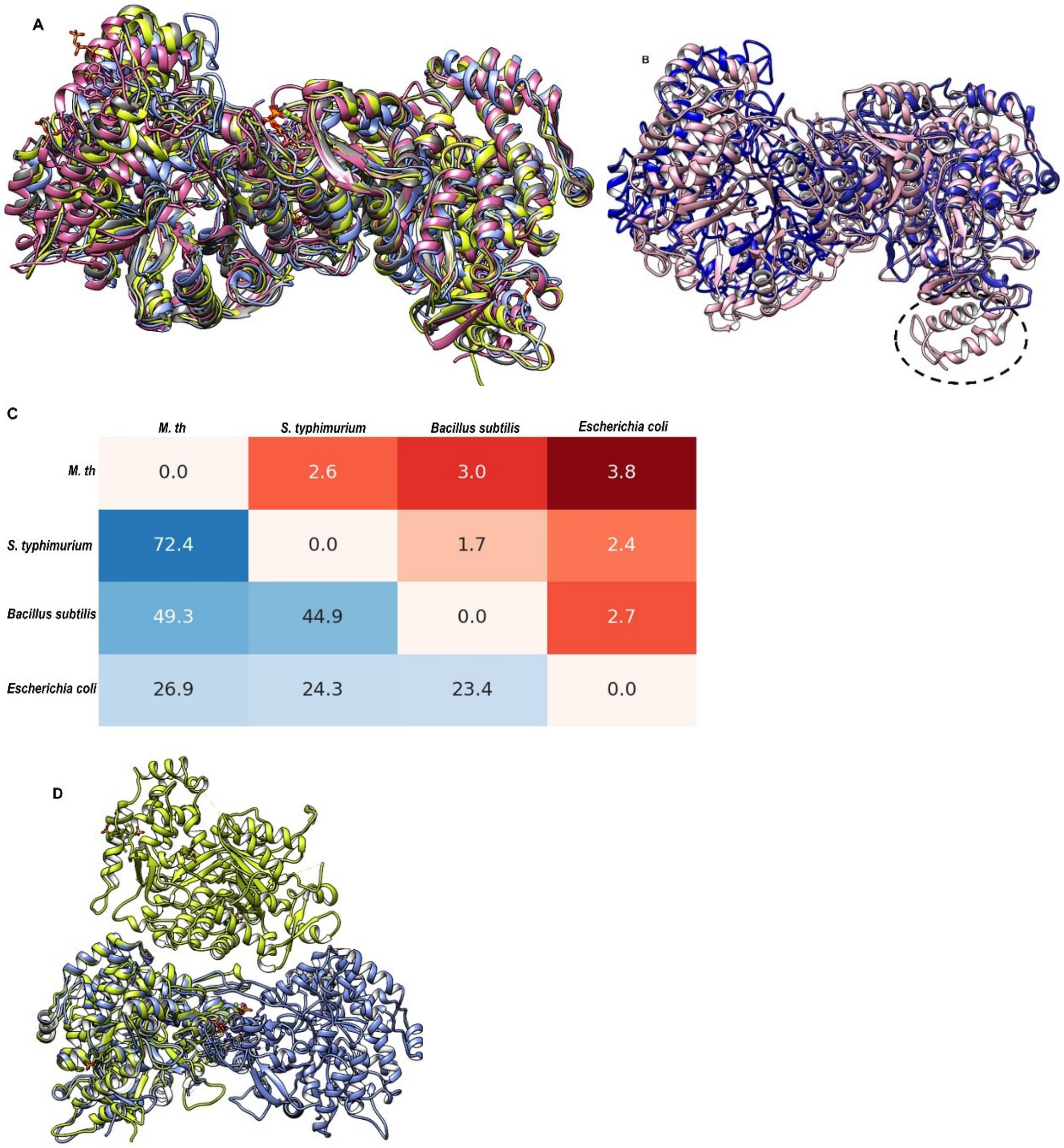


**Supplementary Figure 15: Structural superposition of α-subunit with structure determined in inhibitory and non-inhibitory conditions:**

Structural superposition of **A**. Mth α-subunit (blue) with Class Ib *S. typhimurium* 2BQ1 (green), *S. typhimurium* holo structure 1PEU (grey), and *B. subtilis* holo structure 6MVE (pink), **B.** Mth α-subunit (blue) with Class Ia 2R1R *E. coli* (pink), dotted circle represents the N-terminal cone domain. **C.** The heatmap shows the values of RMSD (red) and percent identity (blue) between Mth α-subunit and the representative structures from *S. typhimurium* (1PEM), *B. subtilis* (6MVE), and *E. coli* (2R1R), **D.** Mth α-subunit (blue) with non-canonical dimer 6CGL *B. subtilis* (green).

**Supplementary Table 1**: Cross-structure statistics of different cryoEM structures determined of α-subunit in holo and apo of Mth.

| Cross-structure statistics RMSD (Å) | | | | | | |
|---|---|---|---|---|---|---|
| **Structure** | **holo1** | **holo2** | **holo3** | **holo4** | **holo5** | **apo** |
| **holo1** | | 2.4 | 2.3 | 2.5 | 2.1 | 2.9 |
| **holo2** | 2.4 | | 2.2 | 2.5 | 2.1 | 2.8 |
| **holo3** | 2.3 | 2.2 | | 2.4 | 1.9 | 2.7 |
| **holo4** | 2.5 | 2.5 | 2.4 | | 2.3 | 3.0 |
| **holo5** | 2.1 | 2.1 | 1.9 | 2.3 | | 2.2 |
| **apo** | 2.9 | 2.8 | 2.7 | 3.0 | 2.2 | |

**Supplementary Table 2**: Cross-structure statistics of different structures determined of α-subunit of Class Ia and 1b. Data are shown for *E. coli* (EC), *S.tyhimurium* (ST) and *B. subtilis* (BS) along with PDB ID and chain name.\

| | | Mth | ST | ST | ST | ST | ST | BS | BS | BS | BS | BS | BS | BS | BS | EC |
|---|---|---|---|---|---|---|---|---|---|---|---|---|---|---|---|---|
| | | Mth | 1PEM_A | 1PEO_A | 1PEQ_A | 1PEU_A | 2BQ1_AB | 6CGL_AB | 6CGM_A | 6CGN_A | 6MT9_A | 6MV9_AB | 6MVE_A | 6MW3_AB | 6MYX_AB | 9DB2_AB |
| Mth | Mth | 0.0 | 2.4 | 2.5 | 2.5 | 2.5 | 2.4 | 24.9 | 2.6 | 2.7 | 2.6 | 2.8 | 2.6 | 3.7 | 3.9 | 3.5 |
| ST | 1PEM_A | 2.5 | 0.0 | 0.3 | 0.3 | 0.3 | 0.5 | 1.0 | 1.0 | 1.1 | 0.9 | 0.9 | 0.9 | 1.5 | 1.5 | 1.3 |
| ST | 1PEO_A | 2.5 | 0.3 | 0.0 | 0.2 | 0.2 | 0.5 | 1.0 | 1.0 | 1.1 | 1.0 | 0.8 | 0.9 | 1.5 | 1.5 | 1.3 |
| ST | 1PEQ_A | 2.5 | 0.3 | 0.2 | 0.0 | 0.3 | 0.5 | 1.0 | 1.0 | 1.1 | 0.9 | 0.8 | 0.9 | 1.5 | 1.5 | 1.4 |
| ST | 1PEU_A | 2.5 | 0.3 | 0.2 | 0.3 | 0.0 | 0.5 | 1.0 | 1.0 | 1.1 | 1.0 | 0.8 | 1.0 | 1.5 | 1.5 | 1.3 |
| ST | 2BQ1_AB | 2.4 | 0.5 | 0.5 | 0.5 | 0.5 | 0.0 | 25.2 | 1.0 | 1.1 | 1.1 | 1.6 | 1.1 | 2.4 | 2.5 | 2.3 |
| BS | 6CGL_AB | 24.9 | 1.0 | 1.0 | 1.0 | 1.0 | 25.2 | 0.0 | 0.4 | 0.5 | 0.5 | 25.0 | 0.5 | 26.0 | 25.7 | 25.2 |
| BS | 6CGM_A | 2.9 | 1.0 | 1.0 | 1.0 | 1.0 | 1.0 | 0.4 | 0.0 | 0.3 | 0.5 | 0.5 | 0.5 | 1.2 | 1.2 | 2.4 |
| BS | 6CGN_A | 3.0 | 1.1 | 1.1 | 1.1 | 1.1 | 1.1 | 0.5 | 0.3 | 0.0 | 0.7 | 0.6 | 0.7 | 1.2 | 1.3 | 2.1 |
| BS | 6MT9_A | 2.8 | 0.9 | 1.0 | 0.9 | 1.0 | 1.1 | 0.5 | 0.5 | 0.7 | 0.0 | 0.3 | 0.1 | 1.2 | 1.1 | 1.9 |
| BS | 6MV9_AB | 2.8 | 0.9 | 0.8 | 0.8 | 0.8 | 1.6 | 25.0 | 0.5 | 0.6 | 0.3 | 0.0 | 0.3 | 1.5 | 1.5 | 2.8 |
| BS | 6MVE_A | 2.8 | 0.9 | 0.9 | 0.9 | 1.0 | 1.1 | 0.5 | 0.5 | 0.7 | 0.1 | 0.3 | 0.0 | 1.2 | 1.1 | 1.9 |
| BS | 6MW3_AB | 3.7 | 1.5 | 1.5 | 1.5 | 1.5 | 2.4 | 26.0 | 1.2 | 1.2 | 1.2 | 1.5 | 1.2 | 0.0 | 0.8 | 3.2 |
| BS | 6MYX_AB | 3.9 | 1.5 | 1.5 | 1.5 | 1.5 | 2.5 | 25.7 | 1.2 | 1.3 | 1.1 | 1.5 | 1.1 | 0.8 | 0.0 | 3.3 |
| EC | 9DB2_AB | 3.5 | 1.3 | 1.3 | 1.4 | 1.3 | 2.3 | 25.2 | 2.4 | 2.1 | 1.9 | 2.8 | 1.9 | 3.2 | 3.3 | 0.0 |
| EC | 1R1R_AB | 4.1 | 1.3 | 1.4 | 1.5 | 1.4 | 2.7 | 25.9 | 2.2 | 2.0 | 1.9 | 3.0 | 1.9 | 2.9 | 3.1 | 1.3 |
| EC | 2R1R_AB | 3.8 | 1.3 | 1.3 | 1.4 | 1.3 | 2.5 | 25.9 | 2.3 | 2.0 | 1.9 | 2.9 | 1.8 | 3.0 | 3.1 | 1.1 |
| EC | 2X0X_AB | 4.0 | 1.3 | 1.3 | 1.5 | 1.4 | 2.9 | 25.7 | 2.4 | 2.1 | 2.1 | 3.2 | 2.1 | 3.5 | 3.8 | 1.4 |
| EC | 2XAK_AB | 4.0 | 1.3 | 1.4 | 1.5 | 1.4 | 2.8 | 25.8 | 2.4 | 2.1 | 2.1 | 3.1 | 2.1 | 3.5 | 3.8 | 1.4 |
| EC | 2XAP_AB | 4.1 | 1.3 | 1.4 | 1.5 | 1.4 | 2.9 | 25.8 | 2.4 | 2.1 | 2.1 | 3.2 | 2.1 | 3.6 | 3.8 | 1.5 |
| EC | 2XAV_AB | 4.1 | 1.3 | 1.4 | 1.5 | 1.4 | 2.9 | 25.8 | 2.5 | 2.1 | 2.1 | 3.5 | 2.1 | 3.5 | 3.7 | 1.5 |
| EC | 2XAW_AB | 3.9 | 1.3 | 1.4 | 1.5 | 1.4 | 2.7 | 25.7 | 2.5 | 2.1 | 2.1 | 3.3 | 2.1 | 3.5 | 3.8 | 1.3 |
| EC | 2XAX_AB | 4.1 | 1.3 | 1.4 | 1.5 | 1.4 | 2.9 | 25.8 | 2.5 | 2.1 | 2.1 | 3.5 | 2.1 | 3.6 | 3.8 | 1.5 |
| EC | 2XAY_AB | 4.1 | 1.3 | 1.3 | 1.5 | 1.4 | 2.9 | 25.8 | 2.4 | 2.1 | 2.1 | 3.2 | 2.1 | 3.5 | 3.7 | 1.4 |
| EC | 2XAZ_AB | 4.1 | 1.3 | 1.4 | 1.5 | 1.4 | 2.9 | 25.8 | 2.4 | 2.1 | 2.1 | 3.2 | 2.1 | 3.5 | 3.8 | 1.4 |
| EC | 2XO4_AB | 4.2 | 1.3 | 1.4 | 1.5 | 1.4 | 2.9 | 25.9 | 2.4 | 2.0 | 1.9 | 3.2 | 1.8 | 3.4 | 3.6 | 1.5 |
| EC | 2XO5_AB | 4.0 | 1.3 | 1.3 | 1.4 | 1.3 | 2.8 | 25.8 | 2.3 | 2.0 | 1.8 | 3.1 | 1.8 | 3.4 | 3.6 | 1.4 |
| EC | 3R1R_AB | 3.8 | 1.3 | 1.3 | 1.4 | 1.3 | 2.6 | 25.8 | 2.4 | 2.0 | 1.8 | 3.0 | 1.8 | 3.3 | 3.5 | 1.2 |
| EC | 3UUS_AB | 3.6 | 1.3 | 1.3 | 1.5 | 1.3 | 2.3 | 25.7 | 2.5 | 2.0 | 1.9 | 2.8 | 1.9 | 3.2 | 3.4 | 1.1 |
| EC | 4ERM_AB | 3.5 | 1.3 | 1.3 | 1.5 | 1.3 | 2.3 | 25.5 | 2.5 | 2.0 | 1.9 | 2.7 | 1.9 | 3.1 | 3.3 | 1.0 |
| EC | 4ERP_AB | 4.2 | 1.3 | 1.4 | 1.5 | 1.4 | 3.0 | 25.8 | 2.5 | 2.2 | 1.9 | 3.1 | 1.9 | 3.3 | 3.5 | 1.5 |
| EC | 4R1R_AB | 4.2 | 1.3 | 1.4 | 1.5 | 1.4 | 3.0 | 25.9 | 2.4 | 1.9 | 1.9 | 3.2 | 1.8 | 3.4 | 3.6 | 1.5 |
| EC | 5CNS_AB | 3.8 | 1.5 | 1.5 | 1.7 | 1.6 | 2.6 | 25.7 | 2.3 | 1.7 | 1.7 | 3.1 | 1.7 | 3.5 | 3.8 | 0.9 |
| EC | 5CNT_AB | 3.8 | 1.5 | 1.6 | 1.7 | 1.6 | 2.6 | 25.6 | 2.3 | 1.8 | 1.8 | 3.0 | 1.8 | 3.6 | 3.8 | 0.8 |
| EC | 5CNU_AB | 3.7 | 1.5 | 1.6 | 1.7 | 1.6 | 2.6 | 25.6 | 2.4 | 1.8 | 1.8 | 3.0 | 1.8 | 3.6 | 3.8 | 0.8 |
| EC | 5CNV_AB | 3.8 | 1.6 | 1.6 | 1.8 | 1.7 | 2.7 | 25.9 | 2.4 | 1.8 | 1.9 | 3.1 | 2.0 | 3.5 | 3.8 | 0.9 |
| EC | 5R1R_AB | 4.0 | 1.3 | 1.4 | 1.5 | 1.4 | 2.7 | 25.7 | 2.4 | 2.1 | 1.9 | 3.1 | 1.9 | 3.4 | 3.6 | 1.3 |
| EC | 6R1R_AB | 3.9 | 1.3 | 1.4 | 1.5 | 1.4 | 2.7 | 25.8 | 2.4 | 2.1 | 1.9 | 3.1 | 1.8 | 3.3 | 3.5 | 1.2 |
| EC | 6W4X_AB | 3.7 | 1.4 | 1.4 | 1.6 | 1.4 | 2.3 | 25.3 | 2.4 | 2.2 | 2.0 | 2.7 | 2.0 | 2.8 | 2.9 | 0.8 |
| EC | 7R1R_AB | 4.1 | 1.3 | 1.4 | 1.5 | 1.3 | 2.7 | 25.8 | 2.4 | 2.1 | 1.9 | 3.1 | 1.8 | 3.4 | 3.5 | 1.2 |
| EC | 8VHN_AB | 3.7 | 1.3 | 1.4 | 1.5 | 1.4 | 2.4 | 25.7 | 2.4 | 2.0 | 1.9 | 2.9 | 1.9 | 3.1 | 3.3 | 1.0 |
| EC | 8VHO_AB | 3.6 | 1.3 | 1.4 | 1.5 | 1.4 | 2.4 | 25.5 | 2.4 | 2.0 | 1.8 | 2.8 | 1.8 | 3.1 | 3.3 | 0.9 |
| EC | 8VHP_AB | 3.7 | 1.6 | 1.6 | 1.7 | 1.6 | 2.6 | 25.8 | 2.5 | 1.7 | 1.8 | 3.1 | 1.8 | 3.7 | 4.0 | 0.9 |
| EC | 8VHQ_AB | 3.8 | 1.3 | 1.3 | 1.5 | 1.4 | 2.6 | 25.8 | 2.4 | 2.0 | 1.9 | 2.9 | 1.8 | 3.3 | 3.4 | 1.2 |
| EC | 8VHR_AB | 3.8 | 1.3 | 1.4 | 1.5 | 1.4 | 2.5 | 25.8 | 2.4 | 1.9 | 1.9 | 2.9 | 1.9 | 3.1 | 3.3 | 1.2 |
| EC | 8VHU_AB | 3.5 | 1.4 | 1.4 | 1.5 | 1.4 | 2.3 | 25.4 | 2.4 | 2.0 | 1.9 | 2.8 | 1.9 | 3.2 | 3.4 | 0.9 |

| | | EC | EC | EC | EC | EC | EC | EC | EC | EC | EC | EC | EC | EC |
|---|---|---|---|---|---|---|---|---|---|---|---|---|---|---|
| | | 1R1R_AB | 2R1R_AB | 2X0X_AB | 2XAK_AB | 2XAP_AB | 2XAV_AB | 2XAW_AB | 2XAX_AB | 2XAY_AB | 2XAZ_AB | 2XO4_AB | 2XO5_AB | 3R1R_AB |
| Mth | Mth | 4.1 | 3.8 | 4.0 | 4.0 | 4.1 | 4.1 | 3.9 | 4.1 | 4.1 | 4.1 | 4.2 | 4.0 | 3.8 |
| ST | 1PEM_A | 1.3 | 1.3 | 1.3 | 1.3 | 1.3 | 1.3 | 1.3 | 1.3 | 1.3 | 1.3 | 1.3 | 1.3 | 1.3 |
| ST | 1PEO_A | 1.4 | 1.3 | 1.3 | 1.4 | 1.4 | 1.4 | 1.4 | 1.4 | 1.3 | 1.4 | 1.4 | 1.3 | 1.3 |
| ST | 1PEQ_A | 1.5 | 1.4 | 1.5 | 1.5 | 1.5 | 1.5 | 1.5 | 1.5 | 1.5 | 1.5 | 1.5 | 1.4 | 1.4 |
| ST | 1PEU_A | 1.4 | 1.3 | 1.4 | 1.4 | 1.4 | 1.4 | 1.4 | 1.4 | 1.4 | 1.4 | 1.4 | 1.3 | 1.3 |
| ST | 2BQ1_AB | 2.7 | 2.5 | 2.9 | 2.8 | 2.9 | 2.9 | 2.7 | 2.9 | 2.9 | 2.9 | 2.9 | 2.8 | 2.6 |
| BS | 6CGL_AB | 25.9 | 25.9 | 25.7 | 25.8 | 25.8 | 25.8 | 25.7 | 25.8 | 25.8 | 25.8 | 25.9 | 26 | 25.8 |
| BS | 6CGM_A | 2.2 | 2.3 | 2.4 | 2.4 | 2.4 | 2.5 | 2.5 | 2.5 | 2.4 | 2.4 | 2.4 | 2.3 | 2.4 |
| BS | 6CGN_A | 2.0 | 2.0 | 2.1 | 2.1 | 2.1 | 2.1 | 2.1 | 2.1 | 2.1 | 2.1 | 2.0 | 2.0 | 2.0 |
| BS | 6MT9_A | 1.9 | 1.9 | 2.1 | 2.1 | 2.1 | 2.1 | 2.1 | 2.1 | 2.1 | 2.1 | 1.9 | 1.8 | 1.8 |
| BS | 6MV9_AB | 3.0 | 2.9 | 3.2 | 3.1 | 3.2 | 3.5 | 3.3 | 3.5 | 3.2 | 3.2 | 3.2 | 3.1 | 3.0 |
| BS | 6MVE_A | 1.9 | 1.8 | 2.1 | 2.1 | 2.1 | 2.1 | 2.1 | 2.1 | 2.1 | 2.1 | 1.8 | 1.8 | 1.8 |
| BS | 6MW3_AB | 2.9 | 3.0 | 3.5 | 3.5 | 3.6 | 3.5 | 3.5 | 3.6 | 3.5 | 3.5 | 3.4 | 3.4 | 3.3 |
| BS | 6MYX_AB | 3.1 | 3.1 | 3.8 | 3.8 | 3.8 | 3.7 | 3.8 | 3.8 | 3.7 | 3.8 | 3.6 | 3.6 | 3.5 |
| EC | 9DB2_AB | 1.3 | 1.1 | 1.4 | 1.4 | 1.5 | 1.5 | 1.3 | 1.5 | 1.4 | 1.4 | 1.5 | 1.4 | 1.2 |
| EC | 1R1R_AB | 0.0 | 0.4 | 0.8 | 0.8 | 0.8 | 0.7 | 0.8 | 0.7 | 0.6 | 0.7 | 0.8 | 0.8 | 0.7 |
| EC | 2R1R_AB | 0.4 | 0.0 | 0.7 | 0.6 | 0.7 | 0.6 | 0.7 | 0.7 | 0.6 | 0.6 | 0.7 | 0.7 | 0.5 |
| EC | 2X0X_AB | 0.8 | 0.7 | 0.0 | 0.2 | 0.1 | 0.3 | 0.3 | 0.2 | 0.2 | 0.2 | 0.2 | 0.2 | 0.4 |
| EC | 2XAK_AB | 0.8 | 0.6 | 0.2 | 0.0 | 0.2 | 0.3 | 0.2 | 0.3 | 0.2 | 0.2 | 0.2 | 0.2 | 0.3 |
| EC | 2XAP_AB | 0.8 | 0.7 | 0.1 | 0.2 | 0.0 | 0.2 | 0.4 | 0.2 | 0.2 | 0.2 | 0.2 | 0.3 | 0.4 |
| EC | 2XAV_AB | 0.7 | 0.6 | 0.3 | 0.3 | 0.2 | 0.0 | 0.4 | 0.2 | 0.2 | 0.2 | 0.2 | 0.4 | 0.5 |
| EC | 2XAW_AB | 0.8 | 0.7 | 0.3 | 0.2 | 0.4 | 0.4 | 0.0 | 0.4 | 0.4 | 0.4 | 0.4 | 0.2 | 0.3 |
| EC | 2XAX_AB | 0.7 | 0.7 | 0.2 | 0.3 | 0.2 | 0.2 | 0.4 | 0.0 | 0.2 | 0.2 | 0.2 | 0.3 | 0.5 |
| EC | 2XAY_AB | 0.6 | 0.6 | 0.2 | 0.2 | 0.2 | 0.2 | 0.4 | 0.2 | 0.0 | 0.1 | 0.2 | 0.3 | 0.4 |
| EC | 2XAZ_AB | 0.7 | 0.6 | 0.2 | 0.2 | 0.2 | 0.2 | 0.4 | 0.2 | 0.1 | 0.0 | 0.2 | 0.3 | 0.4 |
| EC | 2XO4_AB | 0.8 | 0.7 | 0.2 | 0.2 | 0.2 | 0.2 | 0.4 | 0.2 | 0.2 | 0.2 | 0.0 | 0.3 | 0.5 |
| EC | 2XO5_AB | 0.8 | 0.7 | 0.2 | 0.2 | 0.3 | 0.4 | 0.2 | 0.3 | 0.3 | 0.3 | 0.3 | 0.0 | 0.3 |
| EC | 3R1R_AB | 0.7 | 0.5 | 0.4 | 0.3 | 0.4 | 0.5 | 0.3 | 0.5 | 0.4 | 0.4 | 0.5 | 0.3 | 0.0 |
| EC | 3UUS_AB | 1.0 | 0.7 | 0.9 | 0.8 | 0.9 | 0.9 | 0.7 | 0.9 | 0.9 | 0.9 | 0.9 | 0.7 | 0.6 |
| EC | 4ERM_AB | 1.1 | 0.8 | 1.0 | 0.9 | 1.0 | 1.1 | 0.8 | 1.1 | 1.0 | 1.0 | 1.0 | 0.9 | 0.8 |
| EC | 4ERP_AB | 0.8 | 0.8 | 0.8 | 0.8 | 0.8 | 0.7 | 0.9 | 0.8 | 0.7 | 0.8 | 0.8 | 0.8 | 0.8 |
| EC | 4R1R_AB | 0.7 | 0.7 | 0.3 | 0.3 | 0.3 | 0.3 | 0.5 | 0.3 | 0.3 | 0.3 | 0.3 | 0.4 | 0.5 |
| EC | 5CNS_AB | 1.3 | 1.1 | 0.8 | 0.9 | 0.9 | 1.0 | 0.8 | 1.0 | 0.9 | 0.9 | 0.9 | 0.8 | 0.9 |
| EC | 5CNT_AB | 1.3 | 1.0 | 0.9 | 0.9 | 0.9 | 1.0 | 0.8 | 1.0 | 0.9 | 0.9 | 0.9 | 0.8 | 0.8 |
| EC | 5CNU_AB | 1.4 | 1.2 | 1.0 | 1.0 | 1.0 | 1.1 | 1.0 | 1.1 | 1.1 | 1.0 | 1.1 | 1.0 | 0.9 |
| EC | 5CNV_AB | 1.2 | 1.0 | 0.8 | 0.8 | 0.8 | 0.9 | 0.8 | 0.8 | 0.8 | 0.8 | 0.8 | 0.8 | 0.8 |
| EC | 5R1R_AB | 0.8 | 0.6 | 0.3 | 0.3 | 0.4 | 0.4 | 0.2 | 0.4 | 0.4 | 0.3 | 0.4 | 0.2 | 0.2 |
| EC | 6R1R_AB | 0.7 | 0.5 | 0.4 | 0.3 | 0.4 | 0.4 | 0.3 | 0.4 | 0.4 | 0.4 | 0.4 | 0.3 | 0.2 |
| EC | 6W4X_AB | 1.3 | 1.3 | 1.7 | 1.6 | 1.7 | 1.6 | 1.6 | 1.6 | 1.6 | 1.6 | 1.7 | 1.6 | 1.5 |
| EC | 7R1R_AB | 0.6 | 0.5 | 0.3 | 0.3 | 0.4 | 0.4 | 0.3 | 0.4 | 0.3 | 0.3 | 0.4 | 0.3 | 0.2 |
| EC | 8VHN_AB | 0.8 | 0.6 | 0.7 | 0.7 | 0.8 | 0.8 | 0.7 | 0.8 | 0.7 | 0.7 | 0.8 | 0.7 | 0.6 |
| EC | 8VHO_AB | 0.8 | 0.6 | 0.8 | 0.7 | 0.8 | 0.8 | 0.7 | 0.8 | 0.7 | 0.7 | 0.8 | 0.7 | 0.6 |
| EC | 8VHP_AB | 1.5 | 1.3 | 1.0 | 1.1 | 1.1 | 1.2 | 1.0 | 1.2 | 1.1 | 1.1 | 1.1 | 1.1 | 1.1 |
| EC | 8VHQ_AB | 0.7 | 0.5 | 0.5 | 0.5 | 0.6 | 0.6 | 0.5 | 0.6 | 0.5 | 0.5 | 0.6 | 0.5 | 0.4 |
| EC | 8VHR_AB | 0.6 | 0.4 | 0.6 | 0.6 | 0.6 | 0.6 | 0.6 | 0.6 | 0.5 | 0.6 | 0.6 | 0.6 | 0.5 |
| EC | 8VHU_AB | 1.0 | 0.7 | 0.9 | 0.9 | 1.0 | 1.0 | 0.9 | 1.0 | 1.0 | 0.9 | 1.0 | 0.9 | 0.7 |

| | | EC | EC | EC | EC | EC | EC | EC | EC | EC | EC | EC | EC | EC | EC | EC | EC | EC | EC |
|---|---|---|---|---|---|---|---|---|---|---|---|---|---|---|---|---|---|---|---|
| | | 3UUS_AB | 4ERM_AB | 4ERP_AB | 4R1R_AB | 5CNS_AB | 5CNT_AB | 5CNU_AB | 5CNV_AB | 5R1R_AB | 6R1R_AB | 6W4X_AB | 7R1R_AB | 8VHN_AB | 8VHO_AB | 8VHP_AB | 8VHQ_AB | 8VHR_AB | 8VHU_AB |
| Mth | Mth | 3.6 | 3.5 | 4.2 | 4.2 | 3.8 | 3.8 | 3.7 | 3.8 | 4.0 | 3.9 | 3.7 | 4.1 | 3.7 | 3.6 | 3.7 | 3.8 | 3.8 | 3.5 |
| ST | 1PEM_A | 1.3 | 1.3 | 1.3 | 1.3 | 1.5 | 1.5 | 1.5 | 1.6 | 1.3 | 1.3 | 1.4 | 1.3 | 1.3 | 1.3 | 1.6 | 1.3 | 1.3 | 1.4 |
| ST | 1PEO_A | 1.3 | 1.3 | 1.4 | 1.4 | 1.5 | 1.6 | 1.6 | 1.6 | 1.4 | 1.4 | 1.4 | 1.4 | 1.4 | 1.4 | 1.6 | 1.3 | 1.4 | 1.4 |
| ST | 1PEQ_A | 1.5 | 1.5 | 1.5 | 1.5 | 1.7 | 1.7 | 1.7 | 1.8 | 1.5 | 1.5 | 1.6 | 1.5 | 1.5 | 1.5 | 1.7 | 1.5 | 1.5 | 1.5 |
| ST | 1PEU_A | 1.3 | 1.3 | 1.4 | 1.4 | 1.6 | 1.6 | 1.6 | 1.7 | 1.4 | 1.4 | 1.4 | 1.3 | 1.4 | 1.4 | 1.6 | 1.4 | 1.4 | 1.4 |
| ST | 2BQ1_AB | 2.3 | 2.3 | 3.0 | 3.0 | 2.6 | 2.6 | 2.6 | 2.7 | 2.7 | 2.7 | 2.3 | 2.7 | 2.4 | 2.4 | 2.6 | 2.6 | 2.5 | 2.3 |
| BS | 6CGL_AB | 25.7 | 25.5 | 25.8 | 25.9 | 25.7 | 25.6 | 25.6 | 25.9 | 25.7 | 25.8 | 25.3 | 25.8 | 25.7 | 25.5 | 25.8 | 25.8 | 25.8 | 25.4 |
| BS | 6CGM_A | 2.5 | 2.5 | 2.5 | 2.4 | 2.3 | 2.3 | 2.4 | 2.4 | 2.4 | 2.4 | 2.4 | 2.4 | 2.4 | 2.4 | 2.5 | 2.4 | 2.4 | 2.4 |
| BS | 6CGN_A | 2.0 | 2.0 | 2.2 | 1.9 | 1.7 | 1.8 | 1.8 | 1.8 | 2.1 | 2.1 | 2.2 | 2.1 | 2.0 | 2.0 | 1.7 | 2.0 | 1.9 | 2.0 |
| BS | 6MT9_A | 1.9 | 1.9 | 1.9 | 1.9 | 1.7 | 1.8 | 1.8 | 1.9 | 1.9 | 1.9 | 2.0 | 1.9 | 1.9 | 1.8 | 1.8 | 1.9 | 1.9 | 1.9 |
| BS | 6MV9_AB | 2.8 | 2.7 | 3.1 | 3.2 | 3.1 | 3.0 | 3.0 | 3.1 | 3.1 | 3.1 | 2.7 | 3.1 | 2.9 | 2.8 | 3.1 | 2.9 | 2.9 | 2.8 |
| BS | 6MVE_A | 1.9 | 1.9 | 1.9 | 1.8 | 1.7 | 1.8 | 1.8 | 2.0 | 1.9 | 1.8 | 2.0 | 1.8 | 1.9 | 1.8 | 1.8 | 1.8 | 1.9 | 1.9 |
| BS | 6MW3_AB | 3.2 | 3.1 | 3.3 | 3.4 | 3.5 | 3.6 | 3.6 | 3.5 | 3.4 | 3.3 | 2.8 | 3.4 | 3.1 | 3.1 | 3.7 | 3.3 | 3.1 | 3.2 |
| BS | 6MYX_AB | 3.4 | 3.3 | 3.5 | 3.6 | 3.8 | 3.8 | 3.8 | 3.8 | 3.6 | 3.5 | 2.9 | 3.5 | 3.3 | 3.3 | 4.0 | 3.4 | 3.3 | 3.4 |
| EC | 9DB2_AB | 1.1 | 1.0 | 1.5 | 1.5 | 0.9 | 0.8 | 0.8 | 0.9 | 1.3 | 1.2 | 0.8 | 1.2 | 1.0 | 0.9 | 0.9 | 1.2 | 1.2 | 0.9 |
| EC | 1R1R_AB | 1.0 | 1.1 | 0.8 | 0.7 | 1.3 | 1.3 | 1.4 | 1.2 | 0.8 | 0.7 | 1.3 | 0.6 | 0.8 | 0.8 | 1.5 | 0.7 | 0.6 | 1.0 |
| EC | 2R1R_AB | 0.7 | 0.8 | 0.8 | 0.7 | 1.1 | 1.0 | 1.2 | 1.0 | 0.6 | 0.5 | 1.3 | 0.5 | 0.6 | 0.6 | 1.3 | 0.5 | 0.4 | 0.7 |
| EC | 2X0X_AB | 0.9 | 1.0 | 0.8 | 0.3 | 0.8 | 0.9 | 1.0 | 0.8 | 0.3 | 0.4 | 1.7 | 0.3 | 0.7 | 0.8 | 1.0 | 0.5 | 0.6 | 0.9 |
| EC | 2XAK_AB | 0.8 | 0.9 | 0.8 | 0.3 | 0.9 | 0.9 | 1.0 | 0.8 | 0.3 | 0.3 | 1.6 | 0.3 | 0.7 | 0.7 | 1.1 | 0.5 | 0.6 | 0.9 |
| EC | 2XAP_AB | 0.9 | 1.0 | 0.8 | 0.3 | 0.9 | 0.9 | 1.0 | 0.8 | 0.4 | 0.4 | 1.7 | 0.4 | 0.8 | 0.8 | 1.1 | 0.6 | 0.6 | 1.0 |
| EC | 2XAV_AB | 0.9 | 1.1 | 0.7 | 0.3 | 1.0 | 1.0 | 1.1 | 0.9 | 0.4 | 0.4 | 1.6 | 0.4 | 0.8 | 0.8 | 1.2 | 0.6 | 0.6 | 1.0 |
| EC | 2XAW_AB | 0.7 | 0.8 | 0.9 | 0.5 | 0.8 | 0.8 | 1.0 | 0.8 | 0.2 | 0.3 | 1.6 | 0.3 | 0.7 | 0.7 | 1.0 | 0.5 | 0.6 | 0.9 |
| EC | 2XAX_AB | 0.9 | 1.1 | 0.8 | 0.3 | 1.0 | 1.0 | 1.1 | 0.8 | 0.4 | 0.4 | 1.6 | 0.4 | 0.8 | 0.8 | 1.2 | 0.6 | 0.6 | 1.0 |
| EC | 2XAY_AB | 0.9 | 1.0 | 0.7 | 0.3 | 0.9 | 0.9 | 1.1 | 0.8 | 0.4 | 0.4 | 1.6 | 0.3 | 0.7 | 0.7 | 1.1 | 0.5 | 0.5 | 1.0 |
| EC | 2XAZ_AB | 0.9 | 1.0 | 0.8 | 0.3 | 0.9 | 0.9 | 1.0 | 0.8 | 0.3 | 0.4 | 1.6 | 0.3 | 0.7 | 0.7 | 1.1 | 0.5 | 0.6 | 0.9 |
| EC | 2XO4_AB | 0.9 | 1.0 | 0.8 | 0.3 | 0.9 | 0.9 | 1.1 | 0.8 | 0.4 | 0.4 | 1.7 | 0.4 | 0.8 | 0.8 | 1.1 | 0.6 | 0.6 | 1.0 |
| EC | 2XO5_AB | 0.7 | 0.9 | 0.8 | 0.4 | 0.8 | 0.8 | 1.0 | 0.8 | 0.2 | 0.3 | 1.6 | 0.3 | 0.7 | 0.7 | 1.1 | 0.5 | 0.6 | 0.9 |
| EC | 3R1R_AB | 0.6 | 0.8 | 0.8 | 0.5 | 0.9 | 0.8 | 0.9 | 0.8 | 0.2 | 0.2 | 1.5 | 0.2 | 0.6 | 0.6 | 1.1 | 0.4 | 0.5 | 0.7 |
| EC | 3UUS_AB | 0.0 | 0.5 | 1.2 | 1.0 | 0.8 | 0.8 | 0.9 | 0.8 | 0.7 | 0.7 | 1.4 | 0.7 | 0.6 | 0.6 | 0.9 | 0.6 | 0.7 | 0.6 |
| EC | 4ERM_AB | 0.5 | 0.0 | 1.3 | 1.1 | 1.0 | 0.9 | 1.0 | 0.9 | 0.9 | 0.8 | 1.3 | 0.9 | 0.5 | 0.5 | 1.1 | 0.7 | 0.7 | 0.7 |
| EC | 4ERP_AB | 1.2 | 1.3 | 0.0 | 0.8 | 1.4 | 1.4 | 1.6 | 1.4 | 0.8 | 0.8 | 1.6 | 0.8 | 1.1 | 1.1 | 1.7 | 0.9 | 0.9 | 1.1 |
| EC | 4R1R_AB | 1.0 | 1.1 | 0.8 | 0.0 | 1.0 | 1.0 | 1.2 | 0.9 | 0.4 | 0.4 | 1.7 | 0.4 | 0.8 | 0.9 | 1.2 | 0.6 | 0.7 | 1.0 |
| EC | 5CNS_AB | 0.8 | 1.0 | 1.4 | 1.0 | 0.0 | 0.2 | 0.3 | 0.3 | 0.8 | 0.8 | 1.2 | 0.9 | 0.7 | 0.7 | 0.3 | 0.7 | 0.9 | 1.0 |
| EC | 5CNT_AB | 0.8 | 0.9 | 1.4 | 1.0 | 0.2 | 0.0 | 0.3 | 0.3 | 0.8 | 0.8 | 1.1 | 0.9 | 0.7 | 0.7 | 0.3 | 0.8 | 0.9 | 0.9 |
| EC | 5CNU_AB | 0.9 | 1.0 | 1.6 | 1.2 | 0.3 | 0.3 | 0.0 | 0.4 | 0.9 | 0.9 | 1.2 | 1.0 | 0.7 | 0.7 | 0.3 | 0.8 | 1.0 | 0.9 |
| EC | 5CNV_AB | 0.8 | 0.9 | 1.4 | 0.9 | 0.3 | 0.3 | 0.4 | 0.0 | 0.7 | 0.8 | 1.2 | 0.8 | 0.7 | 0.7 | 0.4 | 0.7 | 0.8 | 1.0 |
| EC | 5R1R_AB | 0.7 | 0.9 | 0.8 | 0.4 | 0.8 | 0.8 | 0.9 | 0.7 | 0.0 | 0.1 | 1.5 | 0.2 | 0.6 | 0.6 | 1.0 | 0.4 | 0.5 | 0.8 |
| EC | 6R1R_AB | 0.7 | 0.8 | 0.8 | 0.4 | 0.8 | 0.8 | 0.9 | 0.8 | 0.1 | 0.0 | 1.5 | 0.1 | 0.5 | 0.5 | 1.0 | 0.4 | 0.5 | 0.8 |
| EC | 6W4X_AB | 1.4 | 1.3 | 1.6 | 1.7 | 1.2 | 1.1 | 1.2 | 1.2 | 1.5 | 1.5 | 0.0 | 1.5 | 1.2 | 1.2 | 1.2 | 1.5 | 1.4 | 1.3 |
| EC | 7R1R_AB | 0.7 | 0.9 | 0.8 | 0.4 | 0.9 | 0.9 | 1.0 | 0.8 | 0.2 | 0.1 | 1.5 | 0.0 | 0.6 | 0.6 | 1.1 | 0.4 | 0.5 | 0.8 |
| EC | 8VHN_AB | 0.6 | 0.5 | 1.1 | 0.8 | 0.7 | 0.7 | 0.7 | 0.7 | 0.6 | 0.5 | 1.2 | 0.6 | 0.0 | 0.1 | 0.8 | 0.5 | 0.5 | 0.7 |
| EC | 8VHO_AB | 0.6 | 0.5 | 1.1 | 0.9 | 0.7 | 0.7 | 0.7 | 0.7 | 0.6 | 0.5 | 1.2 | 0.6 | 0.1 | 0.0 | 0.8 | 0.5 | 0.5 | 0.6 |
| EC | 8VHP_AB | 0.9 | 1.1 | 1.7 | 1.2 | 0.3 | 0.3 | 0.3 | 0.4 | 1.0 | 1.0 | 1.2 | 1.1 | 0.8 | 0.8 | 0.0 | 0.9 | 1.0 | 1.1 |
| EC | 8VHQ_AB | 0.6 | 0.7 | 0.9 | 0.6 | 0.7 | 0.8 | 0.8 | 0.7 | 0.4 | 0.4 | 1.5 | 0.4 | 0.5 | 0.5 | 0.9 | 0.0 | 0.3 | 0.7 |
| EC | 8VHR_AB | 0.7 | 0.7 | 0.9 | 0.7 | 0.9 | 0.9 | 1.0 | 0.8 | 0.5 | 0.5 | 1.4 | 0.5 | 0.5 | 0.5 | 1.0 | 0.3 | 0.0 | 0.7 |
| EC | 8VHU_AB | 0.6 | 0.7 | 1.1 | 1.0 | 1.0 | 0.9 | 0.9 | 1.0 | 0.8 | 0.8 | 1.3 | 0.8 | 0.7 | 0.6 | 1.1 | 0.7 | 0.7 | 0.0 |

**Supplementary Videos:**

**Supplementary video 1: PCA of active site motion:**

Visualization of the trajectory in backbone representation along principal component axis 1 showing significant movement of the active site pocket residues highlighted in red

**Supplementary video 2: Loop 2 Arg230 and N-terminal Guides Substrate Positioning:**

Morphing demonstrates substrate movement from external binding site to active site (holo4→holo3)

**Supplementary video 3: PCA of central helices and loop2 motion:**

Visualization of the trajectory along principal component axis 1 showing coordinated movement of the four central helices (magenta) and the loop 2 (orange) (backbone representation)

**Supplementary video 4: Loop4 and loop5 anticorrelation Drives Asymmetry**

Morphing in the order of apo→holo5→holo4→holo1 to demonstrate anticorrelation in loop4 and loop5 in opposite chain of dimer

**Supplementary video 5: 3DVA:**

Continuous heterogeneity of Apo α-subunit observed along components 2. The component resolves the anticorrelated movement among each monomer and also highlight the role of central α-helix in closing and opening the active site pocket via loop2
